# Fine-mapping of *SNCA* in REM sleep behavior disorder and overt synucleinopathies

**DOI:** 10.1101/756528

**Authors:** Lynne Krohn, Richard YJ Wu, Karl Heilbron, Jennifer A. Ruskey, Sandra B. Laurent, Cornelis Blauwendraat, Armaghan Alam, Isabelle Arnulf, Michele T.M. Hu, Yves Dauvilliers, Birgit Högl, Mathias Toft, Kari Anne Bjørnarå, Ambra Stefani, Evi Holzknecht, Christelle Charley Monaca, Abril Beatriz, Giuseppe Plazzi, Elena Antelmi, Luigi Ferini-Strambi, Peter Young, Anna Heidbreder, Valérie Cochen De Cock, Brit Mollenhauer, Friederike Sixel-Döring, Claudia Trenkwalder, Karel Sonka, David Kemlink, Michela Figorilli, Monica Puligheddu, Femke Dijkstra, Mineke Viaene, Wolfang Oertel, Marco Toffoli, Gian Luigi Gigli, Mariarosaria Valente, Jean-François Gagnon, Mike A. Nalls, Andrew B. Singleton, 23andMe Research Team, Alex Desautels, Jacques Y. Montplaisir, Paul Cannon, Owen A. Ross, Bradley F. Boeve, Nicolas Dupré, Edward A. Fon, Ronald B. Postuma, Lasse Pihlstrøm, Guy A. Rouleau, Ziv Gan-Or

## Abstract

**Objective:** REM-sleep behavior disorder (RBD) is a prodromal synucleinopathy, as >80% will eventually convert to overt synucleinopathy. We performed an in-depth analysis of the *SNCA* locus to identify RBD-specific risk variants.

**Methods:** Full sequencing and genotyping of *SNCA* was performed in isolated/idiopathic RBD (iRBD, n=1,076), Parkinson’s disease (PD, n=1,013), and dementia with Lewy bodies (DLB, n=415), and in control subjects (n=6,155). A replication cohort from 23andMe of PD patients with probable RBD (pRBD) was also analyzed (cases n=1,782, controls n=131,250). Adjusted logistic regression models and meta-analyses were performed. Effects on conversion rate were analyzed in 432 RBD patients with available data using Kaplan-Meier survival analysis.

**Results:** A 5’-region *SNCA* variant (rs10005233) was associated with iRBD (OR=1.43, *p*=1.1E-08), which was replicated in pRBD. This variant is in linkage disequilibrium (LD) with other 5’ risk variants across the different synucleinopathies. An independent iRBD-specific suggestive association (rs11732740) was detected at the 3’ of *SNCA* (OR=1.32, *p*=4.7E-04, not statistically significant after Bonferroni correction). Homozygous carriers of both iRBD-specific SNPs were at highly increased risk for iRBD (OR=5.74, *p*=2E-06). The known top PD-associated variant (3’ variant rs356182) had an opposite direction of effect in iRBD compared to PD.

**Interpretation:** There is a distinct pattern of association at the *SNCA* locus in RBD as compared to PD, with an opposite direction of effect at the 3’ of *SNCA*. Several 5’ *SNCA* variants are associated with iRBD and with pRBD in overt synucleinopathies, and may suggest a cognitive component to this region.

## INTRODUCTION

Rapid eye movement (REM) sleep behavior disorder (RBD) is characterized by lack of muscle atonia during REM sleep, causing patients to enact their dreams. Population-wide prevalence of RBD is estimated to be approximately 0.5-1%.^1–3^ Isolated (also referred to as ‘idiopathic’) RBD (iRBD) is a prodromal condition for synucleinopathies, neurodegenerative disorders pathologically characterized by α-synuclein deposition. Within an average of 10-15 years, more than 80% of iRBD cases will convert to Parkinson’s disease (PD, most of whom will eventually develop PD with dementia, PDD^4^), dementia with Lewy bodies (DLB), or in fewer cases multiple system atrophy (MSA).^5–9^ The time between iRBD onset or diagnosis to conversion is highly variable; some iRBD patients convert very rapidly, while others may convert decades after the diagnosis of iRBD,^10^ and the mechanisms affecting the risk for RBD and rate of conversion are mostly unknown.^11^

In recent years, preliminary studies suggested that the genetics of iRBD only partially overlap with those of PD or DLB.^12^ Similar to PD, iRBD has been associated with risk variants in *GBA*,^13^ to the *LRRK2* protective haplotype^14^ and to *TMEM175* variants (Krohn et al., under revision). However, no association has been found with other key PD or DLB risk variants in *LRRK2*,^15^ *MAPT* haplotypes,^16^ and the *APOE* ε4 risk haplotype.^17^ Thus far, the role of the *SNCA* gene in iRBD has not been thoroughly studied. *SNCA* encodes α-synuclein, the main protein component of Lewy-bodies and neurites in synucleinopathies.^18^ Interestingly, in different forms of synucleinopathies, there are different, reportedly independent variants in the *SNCA* locus that have been associated with the risk for the disease. In PD, multiple independent *SNCA* association signals have been identified, but a downstream (3’) *SNCA* single nucleotide polymorphism (SNP) rs356182 has consistently been the top genome-wide association study (GWAS) signal in all large-scale meta-analyses, with a secondary independent association at the 5’ region of *SNCA*.^19,20^ However, the top 3’ PD-associated variant is not associated with DLB, in which a strong association has been demonstrated with a SNP at the 5’ region of *SNCA* (rs7681440).^21^ The latter is in linkage disequilibrium (LD) with the secondary signal in PD. Another marker in LD with this 5’ region signal was associated with probable RBD (pRBD, determined by a validated questionnaire) in PD patients in a previous study.^22^ Other SNPs in the *SNCA* locus have been suggested to be involved in PDD^23^ and Alzheimer’s disease with Lewy body pathology (ADLBV, a variant of AD which also demonstrates diffuse Lewy body pathology).^24^ In MSA, contradicting results have been reported regarding the *SNCA* locus, and the largest analysis thus far refuted the previously reported association.^25^

In the current study, we aimed to thoroughly analyze the *SNCA* locus in RBD. By using a combination of full *SNCA* sequencing and comprehensive SNP genotyping in the largest genetic cohort of iRBD to date, as well as in PD and DLB patients with and without pRBD, we examined *SNCA* association with RBD risk and conversion.

## METHODS

### Population

#### Discovery cohort

The discovery cohort included unrelated, consecutively recruited iRBD (n=1,076) and PD patients (n=733), and controls (n=6,155) of European ancestry (determined for all cases and controls using HapMap v.3 in hg19/GRCh37 and principal component [PC] analysis). iRBD refers to cases who were diagnosed with RBD only, without another defined neurodegenerative synucleinopathy. The control group was composed of elderly (n=225, 63.5±8 years) and young controls (n=650, 36±7 years) of European origin collected in Montreal (52% men), and elderly controls from CARTaGENE (n=5,245, 55±8 years, 41% men), a registry that collects clinical data and DNA in Canada (https://www.cartagene.qc.ca/en/home).^26^ Since the analyzed variant frequencies were similar across these control groups and the low prevalence of iRBD leaves little likelihood of contamination, they were combined for analysis. Homogeneity of the cohorts was confirmed using PC analysis after merging, and PCs were included in statistical tests to account for unknown differences. Variability in sex and age were also taken into account and adjusted for in the statistical analysis.

RBD was diagnosed with video polysomnography (vPSG) according to the International Classification of Sleep Disorders, version 2 (ICSD-2) criteria.^27^ Additional data including age at onset and diagnosis of RBD, eventual phenoconversion to an overt synucleinopathy, and rate of phenoconversion were available for a subset of samples (n=432). PD was diagnosed by movement disorder specialists according to the UK Brain Bank Criteria^28^ without excluding patients who had relatives with PD (to 2015), or International Parkinson Disease and Movement Disorders Society criteria (after 2015).^29^ All study participants signed informed consent forms, and the study protocol was approved by the institutional review boards.

#### Replication cohorts

For analysis of probable RBD (pRBD) in PD and DLB, data was available for a total of 2,450 synucleinopathy patients with pRBD, 774 patients without pRBD, and 131,250 controls. These include a European cohort from 23andMe (PD+pRBD n=1,782, controls n=131,250), Montreal cohort (n=183 with pRBD [PD+pRBD], n=243 without pRBD [PD-pRBD]), a PD cohort from Oslo, Norway (PD+pRBD n=123, PD-pRBD n=147), a PD cohort from the Parkinson’s Progression Marker Initiative^30^ (PPMI, PD+pRBD n=106, PD-pRBD n=276) and a DLB cohort from the Mayo Clinic (DLB+pRBD n=256, DLB-pRBD n=108). Association results including the samples from the Oslo cohort were reported for three *SNCA* risk SNPs in a previous publication.^22^ RBD was assessed by the RBD screening questionnaire (RBDSQ) in the Oslo and PPMI cohorts, and by the RBD1Q question in the Montreal PD, 23andMe, and DLB cohorts. Both questionnaires have high sensitivity and specificity in PD.^31,32^

### Genetic Analysis

#### Single nucleotide polymorphism analysis

DNA was extracted using a standard salting out protocol. iRBD and PD cohorts from Montreal, and DLB cohort from the Mayo Clinic were genotyped using the OmniExpress-24 v1.2 chip (Illumina Inc., approximately 700,000 SNPs) with added NeuroX GWAS custom SNPs, comprised of over 24,000 SNPs associated with neurological diseases. CARTaGENE controls were genotyped using the Infinuim Global Screening Array (GSA, Illumina). Data was converted to PLINK^33^ format and merged using only SNPs genotyped on both platforms (n=~160k). Oslo and PPMI PD samples were genotyped as previously described.^22,34^

Quality control (QC) was performed on the variant level (SNPs excluded if genotype quality<95%, missingness>5%, divergent call rates between cases and controls *p*<1E-04, or departure from the Hardy-Weinberg equilibrium *p*<1E-04) and on the sample level (samples excluded if genotype data showed missingness in>5%, abnormal heterozygosity, conflicting sex assignment, cryptic relatedness pihat>0.125, or non-European ancestry). QC was separately performed on the cohorts and then performed again after merging. Sample QC including ancestry and relatedness confirmed homogeneity across the cohorts, and PCs were calculated on pre-imputed genome-wide merged data of pruned SNPs (pairwise R^2^>0.5) and minor allele frequency (MAF)>0.05 for use in statistical analyses. The merged samples were imputed with the Michigan Imputation Server^26^ using the Haploytype Reference Consortium^35^ r1.1 2016 reference panel and filtered for imputation quality>0.8. Post imputation, a total of 1,862 SNPs in the *SNCA* locus (defined as +/−500kb around the PD top GWAS hit, rs356182) were obtained.

In the 23andMe cohort, DNA extraction and genotyping were performed on saliva samples by the National Genetics Institute (NGI), a CLIA licensed clinical laboratory and a subsidiary of Laboratory Corporation of America. Samples were genotyped on one of five genotyping platforms. The v1 and v2 platforms were variants of the Illumina HumanHap550+ BeadChip, including about 25,000 custom SNPs selected by 23andMe, with a total of about 560,000 SNPs. The v3 platform was based on the Illumina OmniExpress+ BeadChip, with custom content to improve the overlap with the 23andMe v2 array, with a total of about 950,000 SNPs. The v4 platform was a fully customized array, including a lower redundancy subset of v2 and v3 SNPs with additional coverage of lower-frequency coding variation, and about 570,000 SNPs. The v5 platform is an Illumina Infinium Global Screening Array (~640,000 SNPs) supplemented with ~50,000 SNPs of custom content. Samples that failed to reach 98.5% call rate were re-analyzed. Those who did not reach a sufficient call rate were excluded. All individuals included in the analyses provided informed consent and answered surveys online according to the 23andMe human subject protocol, which was reviewed and approved by Ethical & Independent Review Services, a private institutional review board (http://www.eandireview.com). QC was performed similar to the procedure outlined above.

#### Full sequencing of SNCA

The coding and 3’ and 5’ untranslated regions (UTR) of *SNCA* were also sequenced in iRBD patients (n=1,076) and controls (n=910) using Molecular Inversion Probes (MIPs) designed, targeted, and amplified as previously described.^36^ Targeting probes are detailed in Table S1 and the full protocol is available upon request. The MIPs library was sequenced using Illumina HiSeq 4000 platform at the McGill University and Genome Québec Innovation Centre. Sequencing data processing was done by Burrows-Wheeler Alignment (BWA), Genome Analysis Toolkit^37^ (GATK v3.8) for post-alignment adjustments and variant calling, and ANNOVAR^38^ for annotation. Variant frequencies were extracted from two public databases: gnomAD^39^ and PDgene.^40^ Variants were filtered for minimum depth of coverage at 30x, Hardy Weinberg equilibrium *p*>0.001, genotype quality >90%, and missingness <10% in both variants and samples.

### Statistical Analysis

To analyze the association between *SNCA* SNPs and iRBD, case-control logistic regression models were used, with the top 3 PCs (number of PCs required for adjustment were determined by scree plot), sex, and age as covariates. The number of independent tests for multiple testing correction was determined according to pruned number of *SNCA* SNPs with MAF>0.05 by R^2^ > 0.5 (n=191) to avoid an overly stringent significance threshold by correcting separately for SNPs that are in high LD and represent the same haplotype. This method set the Bonferroni-corrected significance threshold at *p*<2.6E-04, which is in line with previously established thresholds in the same *SNCA* region.^34^ Logistic regression was performed in a stepwise forward method^41^ as previously described^34^ to account for linkage disequilibrium (LD) and ensure independence of significant hits, followed by a combined model including all significantly associated signals to ensure independence. LD was calculated and visualized for top hits and known synucleinopathy risk variants in Haploview.^42^ Variants which passed the aforementioned significance threshold were replicated using 23andMe PD+pRBD versus controls including the covariates sex, age, and PCs 1-5. Next, top risk SNPs for iRBD and overt synucleinopathies were analyzed in the PD and DLB cohorts, comparing patients with and without pRBD (PD+/−pRBD, DLB+/−pRBD). Each cohort was analyzed separately, followed by a meta-analysis with the R package metafor,^43^ using logistic regression adjusted for sex, age, and PCs 1-3. The association of rare variants with RBD risk was evaluated in the MIPs sequencing data using optimized sequence Kernel association test (SKAT-O).^44^

To examine whether *SNCA* variants affect the rate of conversion of RBD, we estimated both duration from RBD onset to conversion onset and RBD diagnosis to conversion diagnosis. Data on conversion was available for 432 RBD patients (converted n=237). Top synucleinopathy-associated SNPs and 3’ and 5’ UTR variants with MAF >0.01 from the sequencing data were included in the analysis. The association between SNPs and rate of conversion was examined using Kaplan-Meier survival analysis. The Bonferroni-corrected p value threshold was *p*<0.006, accounting for 8 variants. Analyses were performed using R version 3.5.1.

## RESULTS

### *SNCA* variants included in the analysis

To perform the analysis on high quality GWAS data, a total of 1,862 SNPs in the *SNCA* locus were included in the final analysis (Table S2). From the targeted sequencing of *SNCA*, 53 rare variants were found and included in the SKAT-O analyses, as well as 3 common variants included in conversion analyses (Table S3, including coding, intronic and untranslated region variants). Of the 53 rare variants, 4 were nonsynonymous; p.K97R, p.P117S (rs145138372), and p.A124T (rs1358566725) were each found in a single iRBD patient, and p.N122S (rs749476922) was found in two controls. None were previously associated with PD or any other condition. Each variant is quite rare (MAF < 1e-04 on gnomAD).

### Risk for RBD is primarily associated with 5’ *SNCA* SNPs

Two independent signals, one at the 5’ (rs10005233, OR=1.43, 95%CI=1.27–1.62, *p*=1.1e-08) and one at the 3’ (rs11732740, OR=1.32, 95%CI=1.13-1.53, *p*=4.7e-04) of *SNCA*, were associated with risk for iRBD (Table 1, Figure 2), yet only the 5’ SNP remained statistically significant after Bonferroni correction. Analysis without the Cartagene control samples that were done on a different platform, yielded almost identical results (data not shown). Homozygous carriers of both SNPs were associated with highly increased risk for iRBD (OR=5.74, 95%CI=2.81-11.72, *p*=2E-06), with a gradual increase in risk dependent on the number of alleles with risk variants in these two SNPs (Table 2). The association of the *SNCA* 5’ variant with iRBD was then examined in the 23andMe cohort of PD+pRBD patients (n=1,782) and controls (n=131,250), and was replicated (OR=1.15, 95%CI=1.08-1.23, *p*=4.6E-05). This 5’ variant (rs10005233) is in LD with the previously published top signal for DLB, rs7681440 (R^2^=0.94, D’=0.99),^21^ as well as the secondary PD 5’ GWAS signal rs763443 (R^2^=0.78, D’=0.89)^20^ and a variant previously associated with ADLBV rs2583988 (R^2^=0.40, D’=0.99),^24^ suggesting that one haplotype may drive this association across all synucleinopathies (Figure 1A). The second independent iRBD-associated signal at the 3’ of *SNCA*, rs11732740, did not reach Bonferroni-corrected statistical significance. Notably, this SNP is not in LD with the top GWAS 3’ SNP associated with PD, rs356182^19,20^ (R^2^=0.003, D’=0.14) but is in partial LD with another, independent 3’ PD risk SNP rs2870004, which was recently reported^34^ (R^2^=0.04, D’=0.81).

**Table 1.** *SNCA* variants associated with REM Sleep Behavior Disorder.

| **SNP** | **rs10005233** | **rs11732740** |
| --- | --- | --- |
| Alleles, effect/reference | T/C | G/A |
| GRCh37/hg19 base position | 4:90743331 | 4:90423029 |
| **Analysis, n cases/controls = 1,031/5,864** |  |  |
| Effect allele frequency cases/controls | 0.58 / 0.49 | 0.21 / 0.17 |
| Phred-Scaled CADD | 4.71 | 4.67 |
| **Stepwise Conditional Analysis** |  |  |
| Conditioned on | NA | rs10005233 |
| OR (95% CI) | 1.43 (1.27 – 1.62) | 1.32 (1.13 - 1.53) |
| *p* | 1.11E-08 | 0.00047 |
| **Combined Model** |  |  |
| OR (95% CI) | 1.42 (1.26-1.61) | 1.32 (1.13-1.53) |
| *p* | 2.73E-08 | 0.0005 |

SNP: single nucleotide polymorphism. CADD: combined annotation dependent depletion. OR: odds ratio. CI: confidence interval. NA: not applicable.

**Table 2.** Gradual increase in risk for iRBD in carriers of the top 5’ and 3’ *SNCA* variants.

| **Number of alleles with risk SNPs in rs10005233 and in rs11732740<sup>a</sup>** | **% of carriers in RBD (n=1,076)** | **% of carriers in controls (n=5,897)** | **OR (95% CI)<sup>b</sup>** | ***p*-value<sup>b</sup>** |
| --- | --- | --- | --- | --- |
| **0** | 12.6% | 18.6% | reference | reference |
| **1** | 33.6% | 41.4% | 1.40 (1.08-1.81) | 0.012 |
| **2** | 39.0% | 30.8% | 2.08 (1.60-2.70) | 3.6E-08 |
| **3** | 12.6% | 8.5% | 2.45 (1.75-3.42) | 1.75E-07 |
| **4** | 2.0% | 0.8% | 5.74 (2.81-11.72) | 2E-06 |

<sup>a</sup> 0 – Non-carriers of both risk SNPs; 1 – Carriers of one risk variant, either in rs10005233 or rs11732740; 2 – Carriers of two risk variants, either one in each SNP, or homozygous carriers of one of the SNPs and non-carriers of the second SNP; 3 – Homozygous carriers of one of the risk SNPs and heterozygous carriers of the second risk SNP; 4 – Homozygous carriers of both risk SNPs
<sup>b</sup> Logistic regression adjusted for covariates as detailed in methods.
iRBD, isolated REM-sleep behavior disorder; OR, odds ratio; CI, confidence interval.

**Figure 1.**
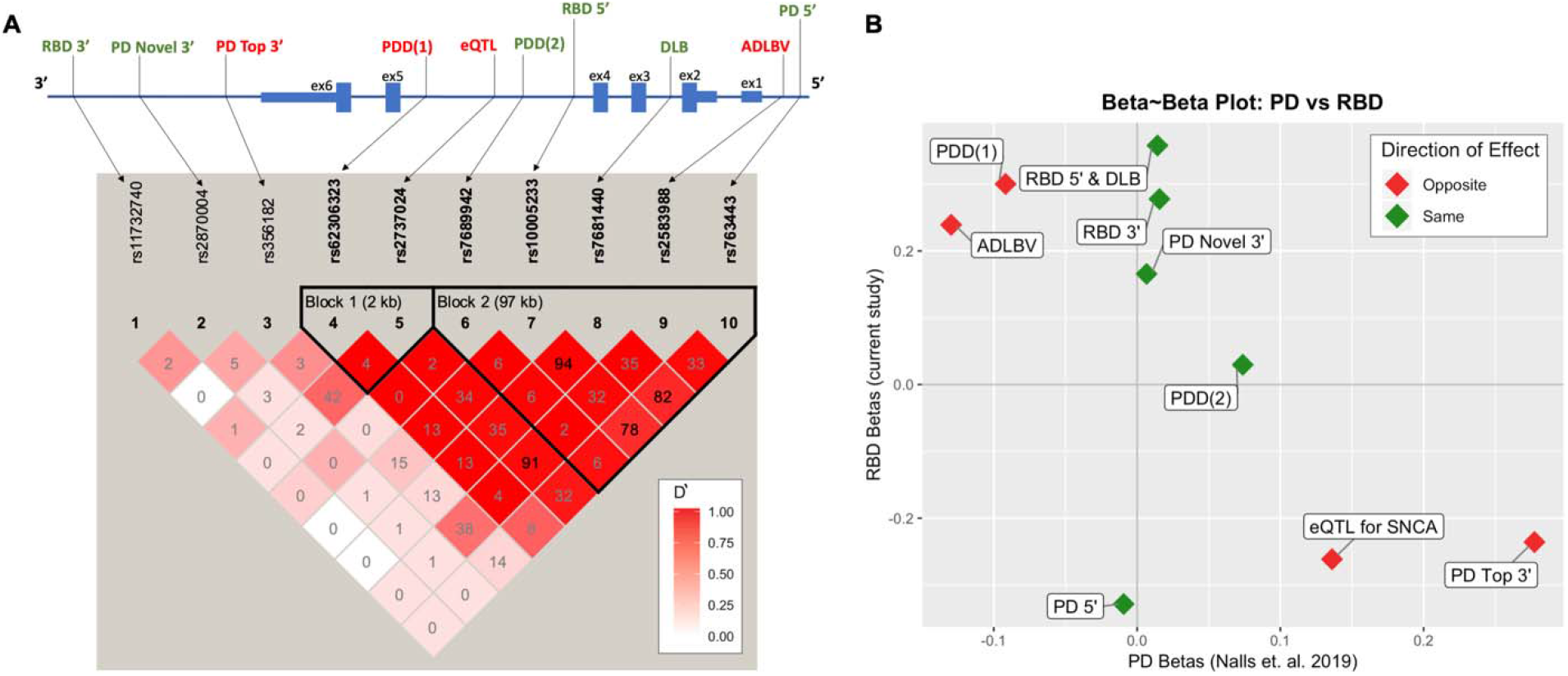
Schematic of top *SNCA* genetic signals across synucleinopathies with a PD to RBD comparison. **A)** A schematic of the *SNCA* region representing the location of top signals for synucleinopathies and linkage disequilibrium (LD) plot by Haploview. In the LD plot, numbers represent R^2^ values and deeper color shade represents strength of D’. Of note, when R^2^ is low and D’ is high, variants are still in strong LD, as the low R^2^ is a result of differences in allele frequencies, yet high D’ means that the less common SNP is in most cases (or all cases if D’=1) appears on the same allele as the more common SNP. Top RBD signal rs10005322 is in moderate to high LD with all 5’ *SNCA* SNPs associated with synucleinopathy. The second RBD signal, 3’ rs11732740, is independent and not in LD with previously reported 3’ or 5’ risk variants. Synucleinopathy risk variants are located in the promoter and regulatory regions, with a strong LD block in the 5’ end. **B)** A beta-beta plot comparing *SNCA* betas found in the current study (y-axis) to betas from the latest PD GWAS (x-axis). Points in red represent different direction of effect between iRBD and PD, while green represents effect in the same direction. Interestingly, the strongest risk variants for PD at the 3’ of the gene are less common in RBD (marked as “PD Top 3’” and “eQTL for *SNCA*”).

**Figure 2.**
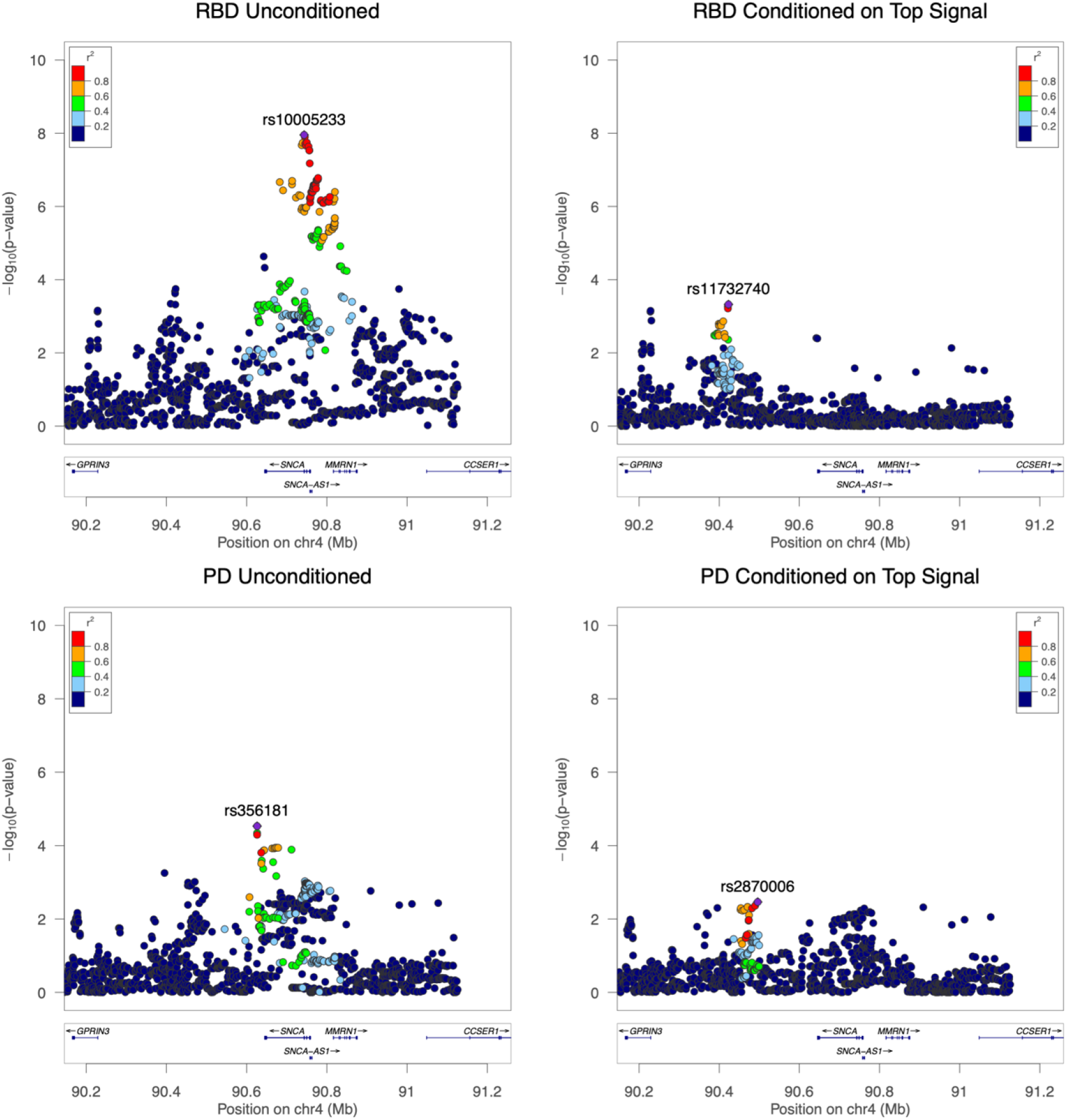
The *SNCA* locus with association results for RBD and PD. Regional LocusZoom Manhattan plots show the top iRBD risk association signal (top left) and the second highest iRBD signal (top right), conditioned on the first. The bottom panels shoe the top PD hits in our PD cohort, demonstrating the different pattern of association between iRBD and PD in the *SNCA* locus. Conditional analysis assures independence of association signals, as shown by the disappearance of the LD block with the RBD top hit rs10005233 on the conditioned plot on the top right. In both iRBD plots, a less significant but notable signal is shown in the region of reported top hit for PD, rs356182, with an opposite direction of effect than the PD signal (OR=0.79, 95% CI=0.70–0.91, *p*=6e-04, in iRBD vs Controls).

To determine whether iRBD was associated with rare *SNCA* variants (MAF<0.01), all rare variants (detailed in Table S3) were analyzed together using SKAT-O (burden), followed by separate analyses of nonsynonymous, synonymous, and UTR variants with the same method. None of the analyses suggested association between any of the types of rare *SNCA* variants and iRBD.

### SNPs previously linked to synucleinopathies and their association with RBD

To examine whether *SNCA* variants previously associated with synucleinopathies are also associated with RBD, we examined the top hits from studies performed on PD,^19,20,34^ PDD,^23^ DLB,^21^ and ADLBV.^24^ Table 3 details the association of these variants with risk for iRBD in our cohort, and the LD with the iRBD 5’ risk variant. Of importance, the effect of several variants was in an opposite direction of effect in iRBD compared to PD. Figure 1B compares the effect sizes (betas) of these top hits between iRBD and PD, based on the most recent meta-analysis for PD.^45^ The top PD signal rs356182-G (3’) is associated with increased risk for PD (OR=1.32, 95%CI=1.30-1.35, *p*=3.9E-194), yet decreased risk for RBD (OR=79, 95%CI=0.70-0.91, *p=*6E-04 in iRBD, Table 3). Likewise, the ADLBV risk variant rs2583988-C is associated with decreased risk for PD (OR=0.88, 95%CI=0.86-0.90, *p*=3.2E-36) and increased risk for iRBD (OR=1.27, 95%CI=1.10-1.48, *p*=0.001), although both without statistical significance after Bonferroni correction in iRBD. Finally, iRBD cases are more likely to carry the DLB-associated haplotype rs62306323-T and rs7689942-C, compared to the reverse which marks the PDD haplotype (Table 3).^23^

**Table 3.** *SNCA* SNPs previously associated with different forms of synucleinopathies and their association with RBD in the current study.

| SNP | Reference | EA | EAF<br>RBD/controls | LD (R<sup>2</sup>/D') |  | Association with RBD in<br>the current study |  |
| --- | --- | --- | --- | --- | --- | --- | --- |
|  |  |  |  | rs10005233 (5') | rs11732740 (3') | OR (95% CI) | p |
| Parkinson's disease |  |  |  |  |  |  |  |
| rs356182 | Top GWAS hit (3')<br>*Chang et. al. 2017* | G | 0.30 / 0.34 | 0.20 / 0.60 | 0.00 / 0.14 | 0.79 (0.70 – 0.91) | 6E-04 |
| rs763443 | GWAS signal (5')<br>*Nalls. et. al. 2014* | C | 0.58 / 0.50 | 0.79 / 0.91 | 0.01 / 0.23 | 1.38 (1.22 – 1.57) | 4E-07 |
| rs2870004 | Novel risk locus (3')<br>*Pihlstrøm et. al. 2018* | T | 0.23 / 0.225 | 0.07 / 0.26 | 0.04 / 0.81 | 1.18 (1.022 – 1.37) | 0.024 |
| rs2737024 | eQTL for SNCA<br>*McClymont et. al. 2018* | G | 0.22 / 0.27 | 0.42 / 1.0 | 0.01 / 0.32 | 0.77 (0.66 – 0.89) | 4E-04 |
| Parkinson's disease with dementia |  |  |  |  |  |  |  |
| rs7689942 | *Guella et. al. 2016* | T | 0.06 / 0.06 | 0.07 / 1.0 | 0.00 / 0.27 | 1.03 (0.80 – 1.33) | 0.80 |
| rs62306323 | *Guella et. al. 2016* | T | 0.14 / 0.12 | 0.15 / 1.0 | 0.00 / 0.20 | 1.35 (1.13 – 1.63) | 0.001 |
| Dementia with Lewy bodies |  |  |  |  |  |  |  |
| rs7681440 | Top GWAS hit (5')<br>*Guerreiro et. al. 2018* | G | 0.58 / 0.50 | 0.94 / 0.99 | 0.00 / 0.00 | 1.42 (1.25 – 1.61) | 3E-08 |
| Alzheimer's disease with Lewy body pathology |  |  |  |  |  |  |  |
| rs2583988 | *Linnertz et. al. 2014* | C | 0.78 / 0.74 | 0.40 / 0.99 | 0.01 / 0.29 | 1.27 (1.10 – 1.48) | 0.001 |

SNP: single nucleotide polymorphism. EA: effect allele. EAF: effect allele frequency. LD: linkage disequilibrium. OR: odds ratio. GWAS: genome-wide association study. eQTL: expression quantitative trait loci.

To further compare the frequencies of *SNCA* variants in RBD and PD, we analyzed the Montreal PD cohort (n=733, controls n=6,155) in the same step-wise, conditional manner in which we analyzed iRBD. Contrary to iRBD and the PD+pRBD replication cohort, the strongest associated SNP with PD in our cohort is downstream (3’). The top signal is rs356181 (Figure 2, OR=1.62, 95%CI=1.13–1.41, *p*=9.9E-05), which is in LD with the top PD GWAS hit, rs356182^15^ (R^2^=0.63, D’=0.99, Figure 1A). After conditioning on the top signal, a second 3’ signal was identified at rs2870006 (OR=0.84, 95%CI=0.75-0.95, *p*=0.0034), also in LD with previously found 3’ signal at rs2870004^34^ (R^2^=0.21, D’=0.99), representing the first independent replication of this association. These SNPs were not in LD with the iRBD two top signals (Table 3), and at separate location than the top iRBD signals (Figure 2).

### Role of *SNCA* in synucleinopathies with probable RBD

We further investigated the association of 5’ and 3’ *SNCA* SNPs with pRBD by comparing allele frequencies of the iRBD associated variants identified in the current study, as well as established synucleinopathy risk-associated SNPs, in DLB and PD patients with and without pRBD. Bonferroni corrected significance threshold was set at *p*<0.01. Results are detailed in Figure 3.

**Figure 3.**
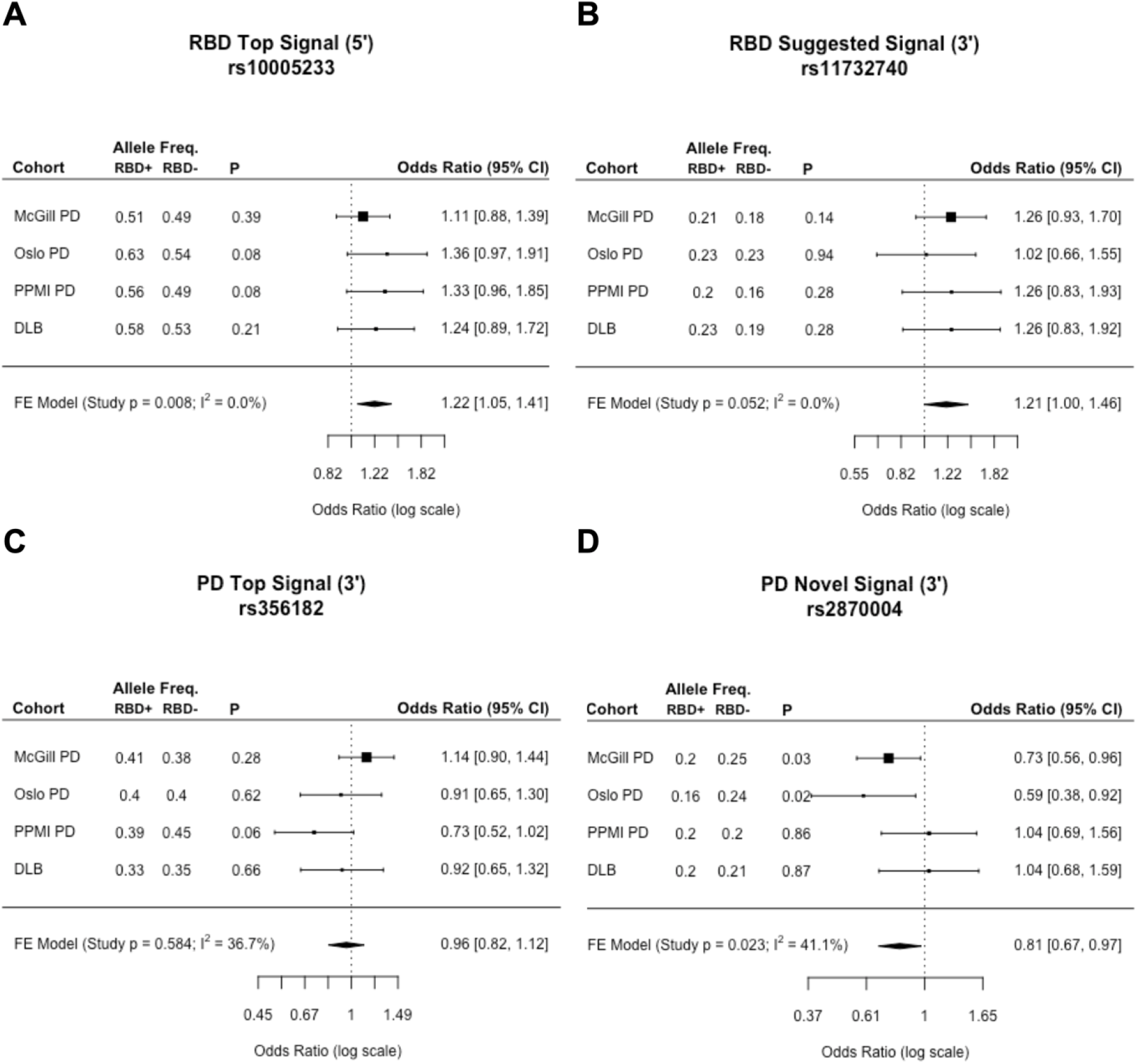
Association between synucleinopathy risk loci in *SNCA* with pRBD in PD and DLB. The forest plots represent the odds ratios and 95% confidence intervals for association of top synucleinopathy risk loci in synucleinopathy with pRBD (PD+pRBD and DLB+pRBD) versus synucleinopathy without RBD (PD-pRBD and DLB-pRBD), as well as results from fixed effect meta-analyses. **A)** The RBD top hit rs10005233 represents all synucleinopathy 5’ variants (rs7681440 and rs763443) because of high LD. 5’ risk variants are associated with synucleinopathies+pRBD, possibly suggesting an RBD-driven signal. **B)** The RBD potential secondary signal shows a trend towards association to synucleinopathy with pRBD, without statistical significance, yet marginal (*p*=0.052). **C)** The PD top GWAS hit rs356182 shows no apparent associations to synucleinopathy with or without pRBD. **D)** Secondary 3’ PD variant rs2870004 is significantly associated with PD without pRBD, suggesting a PD specific signal (FE meta-analysis excluding DLB: OR = 0.76, CI=0.62–0.94, p=0.009). This pattern is not present in DLB, thus is not significant in synucleinopathies overall. Freq.: frequency. OR: odds ratio. CI: confidence interval. FE: fixed effect. pRBD: probable RBD. Syn: Synucleinopathy.

#### 5’ variants

The 5’ variants associated with iRBD, DLB, and PD (rs10005233, rs7681440, and rs763443 respectively) are in high LD with each other (R^2^>0.78, D’>0.9) and are thus represented by the RBD variant rs10005233 in this analysis. We found that 5’ variants are significantly associated with synucleinopathy+pRBD (Fixed Effect [FE] meta-analysis OR=1.22, 95%CI=1.05-1.41, *p*=0.008, Figure 3).

#### 3’ variants

Two independent 3’ risk variants for PD and the 3’ variant associated with pRBD were analyzed. Unlike the 5’ variants, these are independent signals (Figure 1A). Recently reported novel 3’ risk locus for PD, rs2870004,^34^ was significantly associated with pRBD (meta-analysis OR=0.76, CI=0.62–0.94, *p*=0.009), however showed no differences in allele frequencies of DLB+pRBD versus DLB-pRBD, suggesting that this may be a PD-specific signal. Results are inconclusive for the top PD GWAS variant rs356182, showing discrepant distributions of allele frequencies across these cohorts. Similar to the 5’ iRBD risk variant, the secondary iRBD signal rs11732740 shows consistently increased allele frequencies in synucleinopathy+pRBD, however without reaching statistical significance (*p*=0.052, Figure 3B).

### *SNCA* variants and conversion to overt synucleinopathies

Next, we examined whether the top 5’ and 3’ *SNCA* variants for synucleinopathy risk, as well as 3’ and 5’ UTR variants from the sequencing data, were associated with rate of conversion, defined as time from onset or diagnosis from iRBD to diagnosis of an overt neurodegenerative disease in 237 patients who have converted. It must be noted that phenoconversion data was gathered from a cohort where 53% converted to PD, 40% to DLB, and 5% to MSA (2% to “other”), however at random a smaller number of DLB converters had genetic data available, creating a PD-skewed cohort for these analyses. For this reason, we could not perform analysis of variant association with type of conversion.

Top synucleinopathy risk variants from the GWAS data and three common 5’ and 3’ UTR SNPs from the sequencing data were analyzed for association with rate of conversion. One variant, 5’ UTR rs2583986, was associated with faster conversion to overt synucleinopathies (Figure 4, Kaplan-Meier survival analysis *p*=0.0029), when analyzing the rate of conversion from reported onset of RBD to diagnosed onset of overt synucleinopathy. Homozygous carriers of this variant (n=4) had converted within 3.5 ± 1.97 years, Heterozygous carriers (n=30) within 6.2 ± 2.71 years, and non-carriers (n=76) had converted within 9.4 ± 2.05 years. We then performed the same calculation using age at diagnosis to conversion rather than age at onset, and found that this variant lost its significance (*p*>0.05). Interestingly, this variant is in LD with the top risk SNP for iRBD (R^2^=0.38, D’=0.99), however the wild type allele (slower conversion) is correlated with the iRBD risk allele (rs10005233-T). No other *SNCA* variants were associated with rate of conversion from either onset or diagnosis of iRBD.

**Figure 4.**
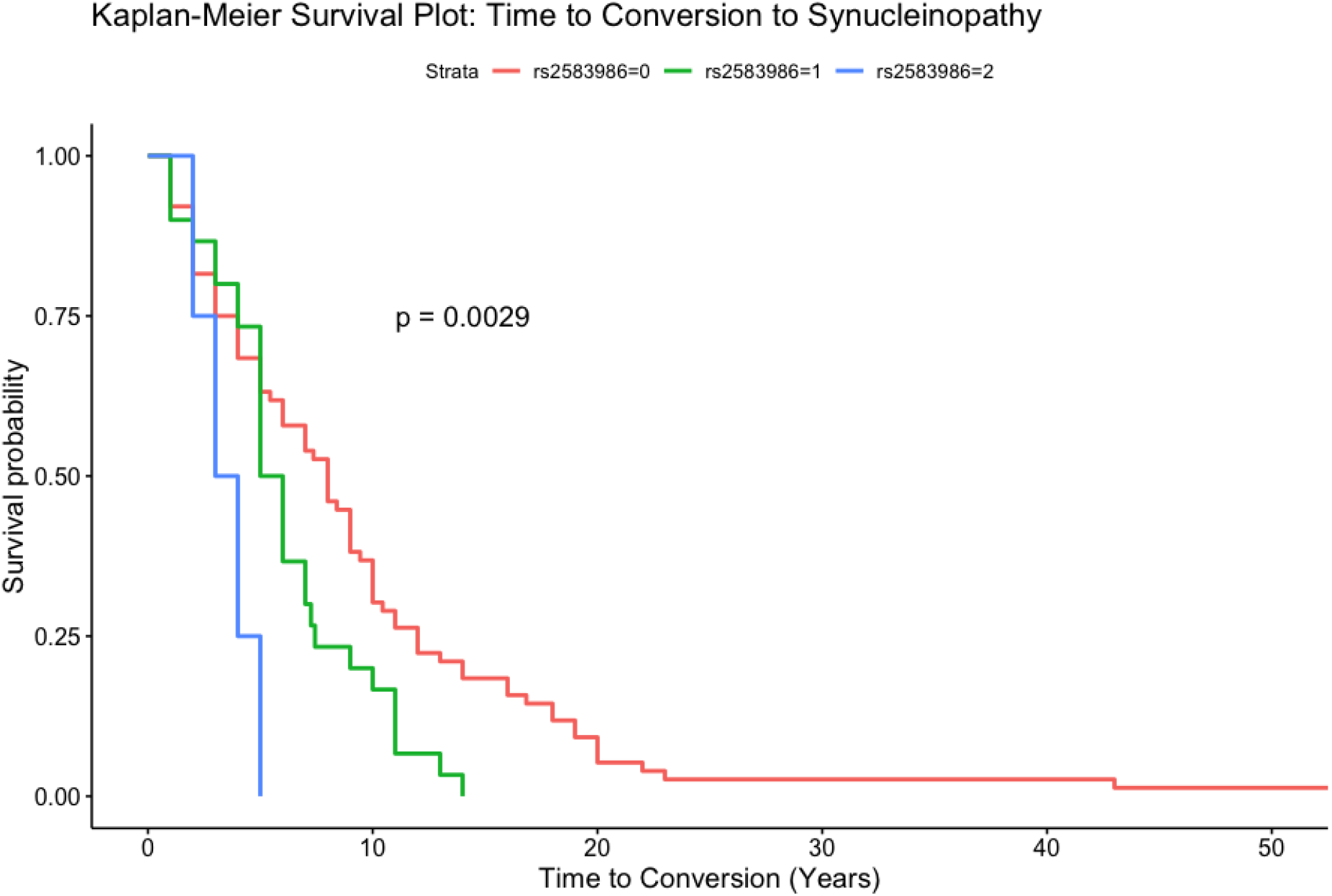
The 5’ UTR SNP rs2583986 and conversion to overt synucleinopathy. Kaplan-Meier survival plot demonstrating a possible effect of rs2583986 genotype of conversion from iRBD to overt synucleinopathy. Wild type carriers (n=76), had mean time to conversion (TTC)=9.4±2.05 years, heterozygous carriers (n=30), had mean TTC=6.2±2.71 years; and homozygous carriers (n=4), had mean TTC=3.5±1.97 years. These results should be interpreted with much caution due to the limitations mentioned in the discussion, including the small numbers and the lack of confidence in patients’ report on RBD onset. Of note, when we performed the same analysis from diagnosis of iRBD to conversion, there was no association between this SNP and conversion.

## DISCUSSION

In the current study we utilized the largest genetic cohort of patients with iRBD published to date, and large replication cohorts with data on pRBD, to perform a fine-mapping study of the *SNCA* locus in RBD. Our results demonstrate that there is a distinct pattern in RBD compared to the most common synucleinopathy, PD. The top association with *SNCA* in RBD is a SNP in the 5’ region of the gene, contrary to the 3’ SNP top association with PD. Additionally, the 5’ RBD variant is associated with susceptibility to pRBD in PD and DLB, suggesting that the 5’ association previously reported in PD and DLB may be driven by the subgroup of patients with RBD. Interestingly, the 3’ PD SNP shows an opposite direction of effect in iRBD, further highlighting the genetic differences. A very high risk for iRBD (OR=5.7) was noted for homozygous carriers of both the 5’ and 3’ iRBD risk variants. We also report a variant (in LD with 5’ risk SNPs) that is potentially associated with rate of conversion to overt synucleinopathy, yet this association has several limitations discussed below, and requires replication.

We have identified two major associations in RBD, a primary association at the 5’ side of *SNCA* and a suggestive secondary association at the 3’ of the gene, both distinct from those identified in our cohort of PD patients (Figure 2). Interestingly, the 5’ top RBD-associated SNP (rs10005233) is in nearly complete LD with the top risk locus for DLB^17^ and is in LD with all 5’ risk SNPs for synucleinopathy (Figure 1A, Table 3), including the 5’ risk variant for PD. This may suggest that all of these associations could be driven by the same variant. The highest CADD score for the variants that we studied was at rs7681440 (11.21), but the association can be driven by other variants. This hypothesis requires functional studies to identify the specific variant or combination of variants that drive these associations. Of note, it is possible that *SNCA* 5’ risk variants in PD and DLB may be driven by the sub-population of patients susceptible to RBD. This is supported by our analyses on PD and DLB cohorts for which we have data on pRBD, where 5’ synucleinopathy risk alleles are significantly associated with synucleinopathy+pRBD. It is important to note that the secondary 3’ RBD-associated SNP, rs11732740 (which did not pass multiple testing correction), is not in LD with neither the PD nor DLB associated SNPs in previous GWASs, and therefore it may be specific to the RBD subtype. This RBD 3’ risk allele is consistently elevated in synucleinopathy with RBD (Figure 3B), however without statistical significance (*p*=0.052) and remains speculative.

The mechanism underlying the association of *SNCA* variants remains unknown. As shown in Figure 1A, the top variants associated with synucleinopathies are concentrated in promoter and regulatory regions of *SNCA*, with a strong LD block towards the 5’ region. The 5’ section of intron four (chr4:90737000–90743400, including RBD variant rs10005233) has been identified as a conserved regulatory region containing a ~160bp CT-rich region.^46^ This region is marked by four distinct haplotypes, one of which is associated with Lewy body pathology and increased α-synuclein levels in Alzheimer’s disease. This haplotype is tagged by rs2298728-A, which is in strong LD with the RBD risk allele rs10005233-T (R^2^=0.06, D’=1, R^2^ is low due to differences in allele frequencies, but the less common SNP always appear on the same allele as the more common SNP). In contrast, a variant in the 3’ end of the intron-four enhancer region rs356168-A has been associated with increased levels of α-synuclein in human iPS cells,^47^ and is in LD with the top signal for PD rs356182-A (R^2^=0.50, D’=0.91). Finally, the top RBD variant rs10005233 has been linked to novel alternative 3′-end *SNCA* isoforms (PB.1016.383, PB.1016.384), associated with a truncated open reading frame prediction.^48^ These findings suggest potential functional effects of some of these variants, but they need to be further replicated and studied.

The independent associations of the 5’ and 3’ variants across the different synucleinopathies may suggest phenotype-specific effects in the regulatory regions on opposing ends of the gene. Although the pathogenic mechanisms of the *SNCA* gene and its encoded protein α-synuclein in synucleinopathies are still not fully understood, our findings suggest that the 5’ region of *SNCA* might affect cognitive components of synucleinopathy. This region is associated with PDD, DLB and ADLBV, which are characterized by cognitive impairment, and now also with RBD, which is known to be the strongest risk factor for rapid and severe cognitive decline in PD.^49^ The differential effects of the 5’ and 3’ SNPs may be due to differential effects in different parts of the brain, and future functional and clinical studies of these regulatory regions will be essential for understanding the pathological mechanisms underlying RBD and these synucleinopathies.

Previous genetic findings also supported an only partially overlapping genetic background between iRBD, PD and DLB.^12^ While *GBA* variants, implicated in both PD^50^ and DLB,^21,51^ were also strongly associated with RBD,^13^ the PD-causing *LRRK2* mutations and the DLB-associated *APOE* ε4 allele were not associated with RBD.^15,17^ The H1/H2 *MAPT* haplotypes, associated with PD and other neurodegenerative disorders, were not associated with RBD either.^16^ More recently, it was demonstrated that the PD-associated *TMEM175* coding variant p.M393T was associated with RBD and potentially affects the activity of the lysosomal enzyme glucocerebrosidase (encoded by *GBA*) (Krohn et al., under revision).^52^ Therefore, it is possible that some genetic variants are relevant for all types of synucleinopathy, with and without RBD, while other variants are specifically relevant for RBD-associated synucleinopathy. Currently, the division between PD and DLB is somewhat arbitrary, determined by a cut-off of one year between the onset of parkinsonism and the onset of dementia.^53^ In late stages of their disease, patients with RBD who converted to PD first and later on developed dementia, and RBD patients who first presented with dementia and later with parkinsonism, can eventually have an undistinguishable clinical presentation. It is possible that in the future this division will be based on genetic background and/or the molecular mechanism involved, such as “*GBA*-associated synucleinopathy”. This would be especially true if treatments that target the underlying genetic cause would be available.

There are several limitations to the current study. One limitation is sample size, as the PD meta-analyses are much larger than the iRBD cohort studied here, which makes it possible that smaller effect size in the *SNCA* locus were not detected in the iRBD cohort. However, with the world’s largest isolated RBD genetic cohort with more than 1000 cases, this study is comparable to previous DLB studies^21,54^ and is sufficiently powered to detect strong associations between *SNCA* and RBD risk. A second limitation lies in the analysis of RBD conversion, which is based on a small number of iRBD patients who had converted to overt synucleinopathies, therefore underpowered to reach corrected statistical significance. Furthermore, the time from onset or diagnosis to conversion is likely an inaccurate estimate of disease duration. Onset of RBD is based on patients’ report, who can be unaware of the actual time when RBD initially presented. Diagnosis time is also not necessarily a good indicator for disease duration, as patients can contact physicians many years after disease onset, depending on whether or not the RBD symptoms disturb the patients or their spouses. Therefore, the results presented here on rate of conversion should be taken as preliminary and with caution. Future studies on larger datasets with adjustments for other clinical variables will enable a more accurate study of the effect of genetics on rate of RBD conversion.

Further studies of RBD genetics are of great importance. As clinical trials in PD have repeatedly failed, it is possible that performing studies on prodromal patients such as iRBD would increase the chances of success in clinical trials since the neurons carry less damage and may therefore be more responsive to treatment. Understanding the underlying genetics of iRBD will enable genetic stratification of patients and may potentially help identify individuals at risk for iRBD and overt synucleinopathies at an earlier stage. Furthermore, genetics can provide targets for drug development (e.g. *GBA* and *LRRK2* in PD) and drive molecular and cellular studies to understand the underlying mechanisms of RBD and synucleinopathies.

## Acknowledgements

We thank the patients and control subjects for their participation in this study. This work was financially supported by Parkinson’s Society Canada, the Michael J. Fox Foundation, the Canadian Consortium on Neurodegeneration in Aging (CCNA), the Canadian Glycomics Network (GlycoNet) and the Canada First Research Excellence Fund (CFREF), awarded to McGill University for the Healthy Brains for Healthy Lives (HBHL) program. JFG holds a Canada Research Chair in Cognitive Decline in Pathological Aging. GAR holds a Canada Research Chair in Genetics of the Nervous System and the Wilder Penfield Chair in Neurosciences. WO is Hertie Senior Research Professor, supported by the Charitable Hertie Foundation, Frankfurt/Main, Germany. EAF holds a Canada Research Chair (Tier 1) in Parkinson Disease. ZGO is supported by the Fonds de recherche du Québec - Santé (FRQS) Chercheurs-boursiers award, and is a Parkinson’s Disease Canada New Investigator awardee. The access to part of the participants for this research has been made possible thanks to the Quebec Parkinson’s Network (http://rpq-qpn.ca/en/). We thank Daniel Rochefort, Helene Catoire and Vessela Zaharieva for their assistance. Mayo Clinic is supported in part by the Mangurian Foundation Lewy Body Dementia Program, The Little Family Foundation, an American Parkinson Disease Association (APDA) Mayo Clinic Information and Referral Center, an APDA Center for Advanced Research and the Mayo Clinic Lewy Body Dementia Association (LBDA) Research Center of Excellence.

## Author contributions

*Conception and design of the study:* LK, ZG-O

*Acquisition and analysis of data:* LK, RYJW, KH, JAR, SBL, CB, AA, IA, MTMH, YD, BH, MT, KAB, AS, EH, CCM, AB, GP, EA, LF-S, PY, AH, VCC, BM, FS-D, CT, KS, DK, MF, MP, FD, VM, WO, MT, GLG, MV, J-FG, MAN, ABS, 23M, AD, JYM, PC, OAR, BFB, ND, EAF, RBP, LP, GAR, ZG-O

*Drafting a significant portion of the manuscript or figures:* LK, ZG-O

