## Supplementary for "Fine-mapping of *SNCA* in REM sleep behavior disorder and overt synucleinopathies"

**Table S1. Molecular Inversion Probes designed to sequence *SNCA* and its regulatory regions.**

| Chr | Probe Start | Probe End | Sequence |
| --- | --- | --- | --- |
| 4 | 90645206 | 90645224 | cactagtgcagagattctgaaCTTCAGCTTCCCGATATCCGACGGTAGTGTttttaggagctctttgtaa |
| 4 | 90645448 | 90645465 | gagatgcaaacatgtttcctcaCTTCAGCTTCCCGATATCCGACGGTAGTGTcatatgactccacggtcg |
| 4 | 90645425 | 90645442 | agtgaatatgagacaagcttccCTTCAGCTTCCCGATATCCGACGGTAGTGTactctgaactgttttggt |
| 4 | 90645636 | 90645651 | gatcaatccagtcctaggtttattCTTCAGCTTCCCGATATCCGACGGTAGTGTctttatatatcttaat |
| 4 | 90645614 | 90645631 | ggtcacaactttcctaatctcaCTTCAGCTTCCCGATATCCGACGGTAGTGTagtgtttggtcccaaata |
| 4 | 90645854 | 90645870 | gaatttaaggatttatgtggataCTTCAGCTTCCCGATATCCGACGGTAGTGTttttaaatctacctaaa |
| 4 | 90645790 | 90645807 | acaaatcctttccttgcttaaaCTTCAGCTTCCCGATATCCGACGGTAGTGTgtctagttctgtcctcta |
| 4 | 90646014 | 90646033 | aggaaggttagaaagtggcgCTTCAGCTTCCCGATATCCGACGGTAGTGTtaattgaagagagactacta |
| 4 | 90645997 | 90646012 | atgaatacatataaactgctagcaCTTCAGCTTCCCGATATCCGACGGTAGTGTtgcagcttagcactct |
| 4 | 90646211 | 90646230 | ggtgaggaaggaaggaggaaCTTCAGCTTCCCGATATCCGACGGTAGTGTctctttttaattattctggg |
| 4 | 90646194 | 90646209 | aagagaaaaacctttcctaaggaaCTTCAGCTTCCCGATATCCGACGGTAGTGTtttagaaatgactatg |
| 4 | 90646433 | 90646449 | cagggaagatctattaactccccCTTCAGCTTCCCGATATCCGACGGTAGTGTtcgttggaactaccaga |
| 4 | 90646410 | 90646428 | aaaagtgaggaatgctgagttCTTCAGCTTCCCGATATCCGACGGTAGTGTgaattgatctcctttaagg |
| 4 | 90646643 | 90646662 | catcagaaattctctctctcCTTCAGCTTCCCGATATCCGACGGTAGTGTgtagaagatgattttgacta |
| 4 | 90646610 | 90646627 | gcattcacaccaatatcagacaCTTCAGCTTCCCGATATCCGACGGTAGTGTgctaatgtgtcttatggc |
| 4 | 90646807 | 90646826 | aaaaaaaagtgggttcccggCTTCAGCTTCCCGATATCCGACGGTAGTGTtgagccttttattaacatat |
| 4 | 90646786 | 90646805 | aatagaccactctacaatagCTTCAGCTTCCCGATATCCGACGGTAGTGTtttcgagacaaaaataacaa |
| 4 | 90647027 | 90647044 | cccttcaatcctgtcaatgtttCTTCAGCTTCCCGATATCCGACGGTAGTGTctcggaattccctgaagc |
| 4 | 90647007 | 90647023 | cagttcttaattcatgttgcttaCTTCAGCTTCCCGATATCCGACGGTAGTGTaacacacttctggcagt |
| 4 | 90647247 | 90647264 | cacaaaggacaaaaatataaagCTTCAGCTTCCCGATATCCGACGGTAGTGTcacttgtgtttgtatata |
| 4 | 90647181 | 90647199 | ggtgcatagtttcatgctcacCTTCAGCTTCCCGATATCCGACGGTAGTGTagtgagatgggataaaaat |
| 4 | 90647419 | 90647436 | atactaaatatgaaattttaccCTTCAGCTTCCCGATATCCGACGGTAGTGTtttgctatcatatattat |
| 4 | 90647395 | 90647414 | catcgtagattgaagccacaCTTCAGCTTCCCGATATCCGACGGTAGTGTcattaaaagacacctaaaaa |
| 4 | 90647623 | 90647640 | acacctaagtgactaccacttaCTTCAGCTTCCCGATATCCGACGGTAGTGTtgaagtatctgtacctgc |
| 4 | 90647604 | 90647621 | aacatctgtcagcagatctcaaCTTCAGCTTCCCGATATCCGACGGTAGTGTaccgaaatgctgagtggg |
| 4 | 90647837 | 90647854 | acaagtgctcagttccaatgtgCTTCAGCTTCCCGATATCCGACGGTAGTGTatactaacagtgtgtgct |
| 4 | 90650282 | 90650300 | gggctccttcttcattctaatCTTCAGCTTCCCGATATCCGACGGTAGTGTgaaatgtgacaatgacagg |
| 4 | 90650524 | 90650539 | acaggaaggaattctggaagatatCTTCAGCTTCCCGATATCCGACGGTAGTGTcgtggccaacatccct |
| 4 | 90743356 | 90743375 | gtcaccactgctcctccaacCTTCAGCTTCCCGATATCCGACGGTAGTGTcaccagcttataaatgtaac |
| 4 | 90743599 | 90743617 | ggtgtgacagcagtagcccagCTTCAGCTTCCCGATATCCGACGGTAGTGTgctagtggaagtggaatga |
| 4 | 90749231 | 90749248 | gtctcaagctagccttaaattgCTTCAGCTTCCCGATATCCGACGGTAGTGTgggccacactaatcacta |
| 4 | 90749366 | 90749383 | gtatctagtgattagtgtggccCTTCAGCTTCCCGATATCCGACGGTAGTGTgcaatttaaggctagctt |
| 4 | 90756658 | 90756675 | cctttgaaagtcctttcatgaaCTTCAGCTTCCCGATATCCGACGGTAGTGTtcatgaacaagcaccaaa |
| 4 | 90756899 | 90756917 | caaggagggagttgtggctgcCTTCAGCTTCCCGATATCCGACGGTAGTGTctcatgatttttcagtgtt |
| 4 | 90757846 | 90757862 | ataatatttaataggaaagaagaCTTCAGCTTCCCGATATCCGACGGTAGTGTgttctccaggatttcca |
| 4 | 90758045 | 90758061 | ggggaggagtcggagttgtggagCTTCAGCTTCCCGATATCCGACGGTAGTGTggcctcctctggggaca |
| 4 | 90757999 | 90758016 | actcccagttctccgctcacgaCTTCAGCTTCCCGATATCCGACGGTAGTGTggaaaaggagcgcacagg |
| 4 | 90758240 | 90758255 | gccattcgacgacaggttagcgggCTTCAGCTTCCCGATATCCGACGGTAGTGTaagtgaggtgcgtgcg |
| 4 | 90758200 | 90758215 | gtcctcctccttctccttctcctcCTTCAGCTTCCCGATATCCGACGGTAGTGTacgctctcggaggggc |
| 4 | 90758422 | 90758437 | gacgaccagaaggggcccaagagaCTTCAGCTTCCCGATATCCGACGGTAGTGTcgaggagcacgctgca |
| 4 | 90759361 | 90759380 | gcctggattcggaagattagCTTCAGCTTCCCGATATCCGACGGTAGTGTcgaacttcaagacaattttt |
| 4 | 90759495 | 90759512 | gcaaaaattgtcttgaagttcgCTTCAGCTTCCCGATATCCGACGGTAGTGTctaatcttccgaatccag |

**Table S2. All variants in the *SNCA* locus analyzed in the GWAS data.**

| **Variant Name** | **Position** | **Allele Effect / Reference** | **Freq. RBD** | **Freq. PD** | **Freq. Controls** |
| --- | --- | --- | --- | --- | --- |
| rs9995167 | 4:90126190 | A / G | 0.15 | 0.15 | 0.14 |
| rs679672 | 4:90126331 | G / A | 0.13 | 0.10 | 0.10 |
| rs583908 | 4:90126713 | T / C | 0.16 | 0.15 | 0.15 |
| rs7674810 | 4:90126898 | C / T | 0.09 | 0.10 | 0.09 |
| rs7696471 | 4:90126909 | T / C | 0.09 | 0.09 | 0.09 |
| rs28735728 | 4:90126959 | C / G | 0.24 | 0.27 | 0.26 |
| rs2037038 | 4:90127328 | G / A | 0.15 | 0.13 | 0.14 |
| rs2043173 | 4:90128492 | C / T | 0.14 | 0.16 | 0.15 |
| rs35825078 | 4:90128682 | C / T | 0.08 | 0.10 | 0.09 |
| rs7663833 | 4:90129219 | T / C | 0.07 | 0.08 | 0.08 |
| rs13140206 | 4:90129890 | C / A | 0.18 | 0.18 | 0.18 |
| rs7675366 | 4:90130491 | G / A | 0.08 | 0.07 | 0.07 |
| rs28442140 | 4:90130753 | T / C | 0.07 | 0.07 | 0.08 |
| rs526210 | 4:90131593 | G / A | 0.47 | 0.47 | 0.47 |
| rs661314 | 4:90132470 | T / C | 0.21 | 0.21 | 0.22 |
| rs557239 | 4:90132664 | C / T | 0.47 | 0.47 | 0.47 |
| rs579195 | 4:90132768 | T / A | 0.47 | 0.47 | 0.47 |
| rs13111385 | 4:90133032 | A / G | 0.09 | 0.09 | 0.09 |
| rs11097217 | 4:90133140 | A / G | 0.15 | 0.15 | 0.15 |
| rs473534 | 4:90133333 | G / A | 0.21 | 0.21 | 0.22 |
| rs686984 | 4:90133612 | G / T | 0.21 | 0.21 | 0.22 |
| rs499411 | 4:90133795 | C / T | 0.47 | 0.48 | 0.47 |
| rs530321 | 4:90134843 | C / T | 0.47 | 0.47 | 0.47 |
| rs612130 | 4:90134953 | C / T | 0.24 | 0.24 | 0.25 |
| rs17015181 | 4:90135036 | G / A | 0.09 | 0.09 | 0.09 |
| rs538005 | 4:90135739 | T / C | 0.33 | 0.33 | 0.34 |
| rs483996 | 4:90135877 | T / C | 0.21 | 0.21 | 0.22 |
| rs6857517 | 4:90135929 | A / T | 0.15 | 0.16 | 0.15 |
| rs74646822 | 4:90136033 | T / C | 0.09 | 0.09 | 0.09 |
| rs17015186 | 4:90136264 | A / G | 0.09 | 0.09 | 0.08 |
| rs17015187 | 4:90136611 | A / C | 0.09 | 0.09 | 0.09 |
| rs642924 | 4:90137256 | C / G | 0.47 | 0.48 | 0.48 |
| rs518156 | 4:90137336 | A / C | 0.24 | 0.24 | 0.25 |
| rs656063 | 4:90137884 | C / A | 0.47 | 0.48 | 0.48 |
| rs17015190 | 4:90138063 | G / A | 0.09 | 0.09 | 0.09 |
| rs576721 | 4:90139073 | G / T | 0.21 | 0.21 | 0.22 |
| rs17772979 | 4:90139147 | G / A | 0.07 | 0.07 | 0.07 |
| rs673366 | 4:90139465 | T / C | 0.47 | 0.47 | 0.48 |
| rs35535962 | 4:90139557 | C / T | 0.09 | 0.09 | 0.09 |
| rs7654139 | 4:90139939 | G / A | 0.32 | 0.31 | 0.30 |
| rs482407 | 4:90141954 | G / A | 0.47 | 0.47 | 0.48 |
| rs17824229 | 4:90142328 | C / G | 0.25 | 0.25 | 0.26 |
| rs10031664 | 4:90142629 | G / T | 0.47 | 0.47 | 0.47 |
| rs72653519 | 4:90143091 | G / A | 0.21 | 0.21 | 0.22 |
| rs72653521 | 4:90143124 | A / C | 0.21 | 0.21 | 0.22 |
| rs4693222 | 4:90143990 | G / A | 0.47 | 0.47 | 0.47 |
| rs7681105 | 4:90144021 | A / G | 0.20 | 0.21 | 0.22 |
| rs6821762 | 4:90144783 | T / A | 0.38 | 0.39 | 0.39 |
| rs13108370 | 4:90145054 | C / G | 0.38 | 0.38 | 0.39 |
| rs13136064 | 4:90145147 | G / T | 0.18 | 0.18 | 0.18 |
| rs34456900 | 4:90145245 | T / C | 0.19 | 0.18 | 0.18 |
| rs6841611 | 4:90146131 | C / T | 0.19 | 0.18 | 0.18 |
| rs9685609 | 4:90146512 | C / T | 0.38 | 0.38 | 0.39 |
| rs9684290 | 4:90146519 | A / G | 0.38 | 0.39 | 0.39 |
| rs17015209 | 4:90146609 | T / A | 0.19 | 0.18 | 0.18 |
| rs9307064 | 4:90146649 | G / A | 0.38 | 0.38 | 0.39 |
| rs17015211 | 4:90147776 | C / T | 0.19 | 0.18 | 0.18 |
| rs1864672 | 4:90149566 | A / G | 0.41 | 0.40 | 0.39 |
| rs17773361 | 4:90149589 | C / A | 0.20 | 0.21 | 0.21 |
| rs4693223 | 4:90150552 | A / G | 0.41 | 0.41 | 0.39 |
| rs1978674 | 4:90151072 | T / G | 0.19 | 0.18 | 0.18 |
| rs17773433 | 4:90151860 | A / T | 0.20 | 0.20 | 0.21 |
| rs10023728 | 4:90152353 | G / A | 0.18 | 0.18 | 0.18 |
| rs1978673 | 4:90152565 | T / C | 0.19 | 0.18 | 0.18 |
| rs1978672 | 4:90152606 | A / G | 0.18 | 0.17 | 0.17 |
| rs10026936 | 4:90153347 | G / A | 0.16 | 0.16 | 0.15 |
| rs6820834 | 4:90153440 | C / T | 0.38 | 0.38 | 0.39 |
| rs6821212 | 4:90153466 | G / A | 0.38 | 0.38 | 0.39 |
| rs4525936 | 4:90153892 | T / C | 0.36 | 0.36 | 0.36 |
| rs746880 | 4:90155040 | T / G | 0.18 | 0.17 | 0.17 |
| rs1431555 | 4:90155732 | C / T | 0.43 | 0.42 | 0.43 |
| rs6532140 | 4:90156461 | C / T | 0.43 | 0.42 | 0.43 |
| rs6853781 | 4:90156489 | T / C | 0.19 | 0.19 | 0.20 |
| rs6532141 | 4:90156629 | G / A | 0.39 | 0.40 | 0.40 |
| rs12500719 | 4:90157051 | A / G | 0.43 | 0.42 | 0.43 |
| rs34465796 | 4:90157854 | T / C | 0.39 | 0.38 | 0.39 |
| rs17773692 | 4:90160401 | T / C | 0.39 | 0.38 | 0.39 |
| rs34115916 | 4:90160486 | T / C | 0.39 | 0.38 | 0.39 |
| rs12504973 | 4:90160697 | A / G | 0.39 | 0.38 | 0.39 |
| rs7660707 | 4:90161863 | C / T | 0.39 | 0.38 | 0.39 |
| rs7682863 | 4:90161995 | A / C | 0.39 | 0.38 | 0.39 |
| rs17015234 | 4:90162268 | A / C | 0.39 | 0.38 | 0.39 |
| rs13110739 | 4:90162842 | A / G | 0.05 | 0.06 | 0.06 |
| rs4396961 | 4:90163617 | A / G | 0.39 | 0.38 | 0.39 |
| rs6532144 | 4:90164136 | C / T | 0.43 | 0.42 | 0.43 |
| rs756004 | 4:90164712 | G / C | 0.39 | 0.39 | 0.39 |
| rs2005264 | 4:90164964 | C / T | 0.39 | 0.38 | 0.39 |
| rs872615 | 4:90165500 | C / T | 0.39 | 0.38 | 0.39 |
| rs872614 | 4:90165642 | C / A | 0.39 | 0.38 | 0.39 |
| rs872613 | 4:90165651 | A / T | 0.39 | 0.38 | 0.39 |
| rs872612 | 4:90165690 | A / G | 0.39 | 0.38 | 0.39 |
| rs3819204 | 4:90165904 | A / T | 0.39 | 0.38 | 0.39 |
| rs891676 | 4:90165952 | C / T | 0.40 | 0.38 | 0.39 |
| rs891675 | 4:90166045 | T / C | 0.40 | 0.38 | 0.39 |
| rs891674 | 4:90166159 | A / G | 0.40 | 0.38 | 0.39 |
| rs17015264 | 4:90166383 | C / A | 0.40 | 0.38 | 0.39 |
| rs6814696 | 4:90166439 | C / T | 0.40 | 0.38 | 0.39 |
| rs731926 | 4:90166679 | G / A | 0.39 | 0.38 | 0.39 |
| rs6532145 | 4:90166773 | C / T | 0.40 | 0.38 | 0.39 |
| rs6532146 | 4:90166778 | A / C | 0.49 | 0.47 | 0.48 |
| rs6532147 | 4:90166920 | T / C | 0.40 | 0.38 | 0.39 |
| rs6532148 | 4:90166921 | G / A | 0.40 | 0.38 | 0.39 |
| rs6838748 | 4:90166957 | A / G | 0.39 | 0.38 | 0.39 |
| rs1835678 | 4:90167253 | C / T | 0.39 | 0.38 | 0.39 |
| rs1431553 | 4:90167294 | C / T | 0.40 | 0.38 | 0.39 |
| rs1431552 | 4:90167374 | T / G | 0.49 | 0.47 | 0.48 |
| rs1431551 | 4:90167376 | T / A | 0.49 | 0.47 | 0.48 |
| rs10516830 | 4:90167445 | G / T | 0.09 | 0.08 | 0.09 |
| rs9790352 | 4:90167489 | G / A | 0.40 | 0.38 | 0.39 |
| rs9790623 | 4:90167506 | C / G | 0.40 | 0.38 | 0.39 |
| rs7686730 | 4:90167542 | T / C | 0.40 | 0.38 | 0.39 |
| rs9790754 | 4:90167597 | G / T | 0.40 | 0.38 | 0.39 |
| rs36120117 | 4:90167633 | T / C | 0.09 | 0.09 | 0.10 |
| rs6844815 | 4:90167781 | G / A | 0.40 | 0.38 | 0.39 |
| rs6828034 | 4:90168336 | C / T | 0.39 | 0.38 | 0.39 |
| rs6850964 | 4:90168340 | T / C | 0.39 | 0.38 | 0.39 |
| rs1431550 | 4:90168415 | T / A | 0.49 | 0.47 | 0.48 |
| rs1431549 | 4:90168566 | A / C | 0.09 | 0.08 | 0.09 |
| rs1431548 | 4:90168572 | T / C | 0.47 | 0.46 | 0.48 |
| rs7672015 | 4:90168825 | C / T | 0.43 | 0.42 | 0.44 |
| rs13115988 | 4:90168882 | A / G | 0.28 | 0.28 | 0.28 |
| rs7653897 | 4:90169925 | A / G | 0.46 | 0.49 | 0.46 |
| rs28622301 | 4:90170118 | G / A | 0.09 | 0.08 | 0.08 |
| rs2298757 | 4:90171478 | A / G | 0.47 | 0.49 | 0.46 |
| rs2298756 | 4:90171503 | C / T | 0.26 | 0.27 | 0.28 |
| rs12500796 | 4:90172474 | T / C | 0.47 | 0.49 | 0.46 |
| rs4270553 | 4:90172722 | A / T | 0.47 | 0.49 | 0.46 |
| rs13126986 | 4:90173451 | T / C | 0.47 | 0.49 | 0.46 |
| rs13106955 | 4:90174346 | G / C | 0.47 | 0.49 | 0.46 |
| rs17825868 | 4:90174968 | T / C | 0.47 | 0.49 | 0.46 |
| rs1125861 | 4:90175113 | A / G | 0.47 | 0.49 | 0.46 |
| rs1367787 | 4:90176942 | T / C | 0.47 | 0.49 | 0.46 |
| rs28432746 | 4:90176943 | A / G | 0.17 | 0.15 | 0.16 |
| rs13142984 | 4:90178067 | C / G | 0.47 | 0.50 | 0.46 |
| rs13118995 | 4:90178131 | T / C | 0.47 | 0.49 | 0.46 |
| rs13149373 | 4:90178671 | C / T | 0.47 | 0.50 | 0.47 |
| rs10011508 | 4:90178849 | A / T | 0.47 | 0.49 | 0.46 |
| rs6532149 | 4:90180437 | A / C | 0.47 | 0.49 | 0.46 |
| rs7690986 | 4:90180489 | G / A | 0.47 | 0.49 | 0.46 |
| rs6849469 | 4:90180934 | A / C | 0.46 | 0.49 | 0.46 |
| rs13141296 | 4:90184334 | G / A | 0.47 | 0.49 | 0.47 |
| rs10470938 | 4:90184428 | G / T | 0.47 | 0.49 | 0.46 |
| rs13140990 | 4:90184457 | C / A | 0.46 | 0.42 | 0.45 |
| rs13117636 | 4:90184707 | C / T | 0.37 | 0.35 | 0.36 |
| rs72653549 | 4:90185078 | A / G | 0.08 | 0.07 | 0.08 |
| rs9307065 | 4:90185448 | C / A | 0.32 | 0.31 | 0.31 |
| rs6825306 | 4:90186236 | C / T | 0.15 | 0.16 | 0.16 |
| rs1125129 | 4:90187746 | C / G | 0.06 | 0.06 | 0.06 |
| rs58290011 | 4:90187929 | G / A | 0.06 | 0.06 | 0.06 |
| rs6532151 | 4:90188353 | C / T | 0.14 | 0.12 | 0.13 |
| rs56688135 | 4:90189122 | A / G | 0.06 | 0.06 | 0.06 |
| rs6824975 | 4:90189313 | T / A | 0.14 | 0.14 | 0.13 |
| rs12331680 | 4:90189860 | C / T | 0.19 | 0.19 | 0.19 |
| rs2116324 | 4:90190815 | C / T | 0.14 | 0.15 | 0.14 |
| rs72653553 | 4:90191792 | A / G | 0.06 | 0.06 | 0.06 |
| rs9991188 | 4:90192063 | C / A | 0.26 | 0.26 | 0.26 |
| rs9991189 | 4:90192064 | G / A | 0.19 | 0.19 | 0.19 |
| rs9993760 | 4:90192423 | G / A | 0.19 | 0.19 | 0.18 |
| rs4425339 | 4:90192559 | G / A | 0.19 | 0.19 | 0.19 |
| rs6830724 | 4:90193727 | C / T | 0.14 | 0.15 | 0.14 |
| rs6854292 | 4:90194160 | G / A | 0.14 | 0.15 | 0.14 |
| rs4538447 | 4:90194685 | C / A | 0.05 | 0.06 | 0.06 |
| rs4399946 | 4:90194997 | T / C | 0.06 | 0.06 | 0.06 |
| rs4475113 | 4:90196192 | G / T | 0.06 | 0.06 | 0.06 |
| rs63674162 | 4:90196552 | C / T | 0.43 | 0.42 | 0.43 |
| rs1560489 | 4:90198463 | C / T | 0.22 | 0.21 | 0.21 |
| rs17015323 | 4:90205166 | T / C | 0.23 | 0.21 | 0.22 |
| rs12640416 | 4:90205253 | C / A | 0.39 | 0.36 | 0.36 |
| rs56351347 | 4:90205612 | C / T | 0.20 | 0.19 | 0.19 |
| rs1835679 | 4:90206392 | G / A | 0.37 | 0.34 | 0.34 |
| rs112440335 | 4:90206433 | A / G | 0.20 | 0.18 | 0.19 |
| rs62304747 | 4:90206621 | A / G | 0.06 | 0.06 | 0.06 |
| rs10014765 | 4:90207422 | C / A | 0.16 | 0.15 | 0.15 |
| rs17015332 | 4:90207602 | C / T | 0.20 | 0.19 | 0.19 |
| rs11722128 | 4:90207810 | A / G | 0.37 | 0.34 | 0.34 |
| rs11730448 | 4:90207849 | G / A | 0.17 | 0.15 | 0.16 |
| rs13106910 | 4:90207956 | G / A | 0.17 | 0.15 | 0.16 |
| rs4693990 | 4:90208148 | G / A | 0.37 | 0.34 | 0.34 |
| rs75764336 | 4:90208169 | G / A | 0.10 | 0.11 | 0.11 |
| rs4693991 | 4:90208438 | A / G | 0.17 | 0.15 | 0.15 |
| rs13140788 | 4:90208758 | T / A | 0.17 | 0.15 | 0.15 |
| rs1367789 | 4:90209159 | C / T | 0.17 | 0.15 | 0.16 |
| rs937719 | 4:90209723 | A / G | 0.37 | 0.34 | 0.34 |
| rs10428466 | 4:90211343 | G / C | 0.20 | 0.19 | 0.19 |
| rs76520868 | 4:90211409 | T / C | 0.07 | 0.07 | 0.06 |
| rs2005104 | 4:90211779 | T / C | 0.37 | 0.34 | 0.34 |
| rs1079070 | 4:90211918 | T / C | 0.37 | 0.34 | 0.34 |
| rs1079069 | 4:90212053 | C / T | 0.37 | 0.34 | 0.34 |
| rs55920273 | 4:90212600 | C / T | 0.20 | 0.19 | 0.19 |
| rs1431546 | 4:90213766 | G / C | 0.17 | 0.15 | 0.16 |
| rs1019875 | 4:90214108 | T / C | 0.17 | 0.15 | 0.15 |
| rs754750 | 4:90214617 | C / G | 0.46 | 0.49 | 0.47 |
| rs7687888 | 4:90216747 | T / G | 0.33 | 0.32 | 0.33 |
| rs7670752 | 4:90218019 | G / T | 0.33 | 0.33 | 0.34 |
| rs7671122 | 4:90218221 | A / G | 0.20 | 0.18 | 0.18 |
| rs7671160 | 4:90218294 | A / G | 0.20 | 0.18 | 0.18 |
| rs1346946 | 4:90218311 | T / C | 0.33 | 0.33 | 0.34 |
| rs12498405 | 4:90219493 | T / G | 0.46 | 0.48 | 0.47 |
| rs1816367 | 4:90219653 | G / A | 0.20 | 0.18 | 0.18 |
| rs1431545 | 4:90220659 | C / T | 0.33 | 0.33 | 0.34 |
| rs56266320 | 4:90220910 | T / G | 0.20 | 0.19 | 0.18 |
| rs1394342 | 4:90221525 | C / T | 0.33 | 0.33 | 0.34 |
| rs1821102 | 4:90222456 | C / A | 0.33 | 0.33 | 0.33 |
| rs1367786 | 4:90222481 | G / T | 0.46 | 0.48 | 0.47 |
| rs6839114 | 4:90223103 | C / T | 0.46 | 0.48 | 0.47 |
| rs10516832 | 4:90223149 | T / C | 0.20 | 0.18 | 0.18 |
| rs2174325 | 4:90223207 | C / A | 0.20 | 0.18 | 0.18 |
| rs1814762 | 4:90223345 | C / T | 0.33 | 0.33 | 0.34 |
| rs2174326 | 4:90223611 | G / A | 0.46 | 0.48 | 0.47 |
| rs6826449 | 4:90224925 | A / C | 0.23 | 0.22 | 0.22 |
| rs1560488 | 4:90225835 | T / C | 0.23 | 0.22 | 0.22 |
| rs7653939 | 4:90227469 | A / G | 0.46 | 0.48 | 0.47 |
| rs1036111 | 4:90227976 | C / T | 0.47 | 0.49 | 0.48 |
| rs72653573 | 4:90228233 | C / T | 0.06 | 0.06 | 0.06 |
| rs17015404 | 4:90228238 | T / C | 0.33 | 0.29 | 0.29 |
| rs1560486 | 4:90228612 | G / C | 0.20 | 0.22 | 0.22 |
| rs2116327 | 4:90228697 | A / G | 0.33 | 0.29 | 0.29 |
| rs28378092 | 4:90228807 | T / C | 0.47 | 0.49 | 0.48 |
| rs919616 | 4:90229385 | A / G | 0.19 | 0.20 | 0.21 |
| rs919615 | 4:90229400 | A / G | 0.47 | 0.49 | 0.48 |
| rs919614 | 4:90229607 | G / C | 0.19 | 0.20 | 0.21 |
| rs6830590 | 4:90230495 | T / C | 0.47 | 0.49 | 0.48 |
| rs28523829 | 4:90230717 | A / G | 0.33 | 0.30 | 0.30 |
| rs72653575 | 4:90231102 | A / G | 0.20 | 0.18 | 0.18 |
| rs1431560 | 4:90231344 | C / G | 0.41 | 0.44 | 0.42 |
| rs62306575 | 4:90231536 | A / G | 0.07 | 0.06 | 0.06 |
| rs72874818 | 4:90231898 | G / T | 0.06 | 0.05 | 0.06 |
| rs71633375 | 4:90233236 | A / G | 0.07 | 0.08 | 0.08 |
| rs4693992 | 4:90233839 | A / G | 0.17 | 0.17 | 0.17 |
| rs34332773 | 4:90234041 | A / G | 0.10 | 0.09 | 0.10 |
| rs35874402 | 4:90234114 | C / T | 0.12 | 0.10 | 0.11 |
| rs4693993 | 4:90234948 | A / G | 0.22 | 0.23 | 0.23 |
| rs1431559 | 4:90235488 | A / G | 0.19 | 0.21 | 0.20 |
| rs12508378 | 4:90235778 | T / C | 0.42 | 0.45 | 0.44 |
| rs112417608 | 4:90235979 | A / G | 0.06 | 0.06 | 0.07 |
| rs62306577 | 4:90236017 | A / C | 0.06 | 0.06 | 0.07 |
| rs10034276 | 4:90237281 | G / A | 0.12 | 0.13 | 0.12 |
| rs10015406 | 4:90238073 | A / G | 0.21 | 0.22 | 0.22 |
| rs28688334 | 4:90238092 | A / G | 0.12 | 0.13 | 0.12 |
| rs28714536 | 4:90238733 | T / C | 0.49 | 0.49 | 0.49 |
| rs11097218 | 4:90239038 | G / A | 0.32 | 0.34 | 0.32 |
| rs139871031 | 4:90239329 | A / G | 0.09 | 0.08 | 0.09 |
| rs13104039 | 4:90241029 | G / T | 0.07 | 0.07 | 0.08 |
| rs80349569 | 4:90241463 | T / A | 0.09 | 0.08 | 0.09 |
| rs7664724 | 4:90241483 | G / A | 0.21 | 0.22 | 0.21 |
| rs17015418 | 4:90241518 | A / G | 0.10 | 0.08 | 0.09 |
| rs10027938 | 4:90242059 | A / G | 0.16 | 0.16 | 0.17 |
| rs13141463 | 4:90242334 | A / C | 0.16 | 0.16 | 0.17 |
| rs13141880 | 4:90242532 | G / C | 0.06 | 0.06 | 0.08 |
| rs17015420 | 4:90242945 | G / A | 0.10 | 0.09 | 0.09 |
| rs28661065 | 4:90243330 | C / T | 0.10 | 0.09 | 0.09 |
| rs10001189 | 4:90244026 | T / C | 0.21 | 0.22 | 0.21 |
| rs4106153 | 4:90244476 | C / A | 0.21 | 0.22 | 0.21 |
| rs62306578 | 4:90245451 | A / G | 0.06 | 0.06 | 0.08 |
| rs17776097 | 4:90247163 | T / G | 0.06 | 0.06 | 0.07 |
| rs17832698 | 4:90248527 | T / C | 0.06 | 0.06 | 0.07 |
| rs2202369 | 4:90249028 | T / A | 0.09 | 0.09 | 0.09 |
| rs35779808 | 4:90249737 | A / G | 0.07 | 0.07 | 0.08 |
| rs28721334 | 4:90250200 | G / A | 0.09 | 0.09 | 0.09 |
| rs28599627 | 4:90250484 | T / C | 0.09 | 0.09 | 0.09 |
| rs28560034 | 4:90250583 | A / G | 0.09 | 0.09 | 0.09 |
| rs28626395 | 4:90250596 | T / G | 0.09 | 0.09 | 0.09 |
| rs76752714 | 4:90251117 | C / T | 0.09 | 0.09 | 0.09 |
| rs62306580 | 4:90251267 | T / G | 0.06 | 0.07 | 0.07 |
| rs34577169 | 4:90251314 | C / T | 0.06 | 0.07 | 0.07 |
| rs6812359 | 4:90252703 | G / A | 0.09 | 0.09 | 0.09 |
| rs34920535 | 4:90253013 | A / G | 0.16 | 0.16 | 0.16 |
| rs79317284 | 4:90253185 | C / T | 0.09 | 0.09 | 0.09 |
| rs11935294 | 4:90253613 | A / G | 0.20 | 0.20 | 0.21 |
| rs62306582 | 4:90254102 | A / T | 0.06 | 0.07 | 0.07 |
| rs11735870 | 4:90255150 | A / G | 0.42 | 0.42 | 0.42 |
| rs1463908 | 4:90255167 | G / A | 0.31 | 0.32 | 0.30 |
| rs73847640 | 4:90255687 | C / T | 0.06 | 0.07 | 0.08 |
| rs1875863 | 4:90256211 | C / T | 0.42 | 0.42 | 0.42 |
| rs2869995 | 4:90256463 | G / T | 0.31 | 0.32 | 0.30 |
| rs10002808 | 4:90258402 | G / C | 0.21 | 0.23 | 0.22 |
| rs1504489 | 4:90258588 | T / G | 0.42 | 0.42 | 0.42 |
| rs13135157 | 4:90262643 | T / A | 0.31 | 0.32 | 0.30 |
| rs13108105 | 4:90262909 | A / G | 0.31 | 0.32 | 0.30 |
| rs13108493 | 4:90263021 | C / T | 0.36 | 0.36 | 0.35 |
| rs62306583 | 4:90263770 | A / G | 0.06 | 0.07 | 0.07 |
| rs6532162 | 4:90263893 | A / G | 0.31 | 0.32 | 0.30 |
| rs2869996 | 4:90264361 | C / T | 0.31 | 0.31 | 0.30 |
| rs12710850 | 4:90264609 | G / T | 0.31 | 0.32 | 0.30 |
| rs13150374 | 4:90266958 | C / T | 0.47 | 0.48 | 0.48 |
| rs1394343 | 4:90267231 | A / G | 0.41 | 0.42 | 0.42 |
| rs6842203 | 4:90267380 | G / A | 0.31 | 0.32 | 0.30 |
| rs13133877 | 4:90268041 | T / G | 0.31 | 0.32 | 0.30 |
| rs13119214 | 4:90269335 | A / G | 0.31 | 0.32 | 0.30 |
| rs13119908 | 4:90269546 | A / C | 0.06 | 0.06 | 0.07 |
| rs74354915 | 4:90270452 | C / A | 0.06 | 0.06 | 0.07 |
| rs11721752 | 4:90270991 | T / C | 0.30 | 0.31 | 0.30 |
| rs13134158 | 4:90271319 | G / C | 0.06 | 0.06 | 0.07 |
| rs13134747 | 4:90271442 | T / G | 0.06 | 0.06 | 0.07 |
| rs28475836 | 4:90271485 | T / C | 0.42 | 0.42 | 0.42 |
| rs17015458 | 4:90271894 | A / G | 0.08 | 0.08 | 0.07 |
| rs10516834 | 4:90271978 | T / C | 0.09 | 0.08 | 0.08 |
| rs10516833 | 4:90272120 | G / A | 0.31 | 0.32 | 0.30 |
| rs1394338 | 4:90273016 | C / T | 0.30 | 0.31 | 0.30 |
| rs1394339 | 4:90273098 | A / C | 0.48 | 0.48 | 0.48 |
| rs17776587 | 4:90275862 | G / A | 0.41 | 0.42 | 0.42 |
| rs12511565 | 4:90277638 | G / A | 0.41 | 0.42 | 0.42 |
| rs13140982 | 4:90278279 | A / G | 0.30 | 0.31 | 0.29 |
| rs17833214 | 4:90278805 | T / C | 0.05 | 0.06 | 0.07 |
| rs13101936 | 4:90279299 | C / A | 0.25 | 0.26 | 0.26 |
| rs10856874 | 4:90279758 | C / T | 0.21 | 0.22 | 0.21 |
| rs17015492 | 4:90280428 | A / G | 0.37 | 0.37 | 0.37 |
| rs6846715 | 4:90280688 | A / G | 0.21 | 0.21 | 0.20 |
| rs4693994 | 4:90280761 | A / G | 0.21 | 0.22 | 0.21 |
| rs4693995 | 4:90280906 | T / C | 0.21 | 0.21 | 0.20 |
| rs71609559 | 4:90281464 | T / C | 0.07 | 0.06 | 0.07 |
| rs35299795 | 4:90282186 | A / G | 0.07 | 0.06 | 0.07 |
| rs13125130 | 4:90282599 | G / A | 0.37 | 0.37 | 0.37 |
| rs17838994 | 4:90282656 | A / G | 0.07 | 0.06 | 0.07 |
| rs7680595 | 4:90282853 | G / T | 0.07 | 0.06 | 0.07 |
| rs7655932 | 4:90282910 | A / G | 0.21 | 0.21 | 0.20 |
| rs6818633 | 4:90283325 | G / A | 0.23 | 0.23 | 0.23 |
| rs13435597 | 4:90283497 | G / A | 0.09 | 0.09 | 0.08 |
| rs34685585 | 4:90283792 | G / A | 0.07 | 0.06 | 0.07 |
| rs10004082 | 4:90283806 | G / A | 0.28 | 0.28 | 0.28 |
| rs10004107 | 4:90284002 | C / T | 0.28 | 0.28 | 0.29 |
| rs36085188 | 4:90284332 | C / T | 0.07 | 0.06 | 0.08 |
| rs2904271 | 4:90284516 | C / A | 0.37 | 0.37 | 0.36 |
| rs6821767 | 4:90285544 | G / T | 0.32 | 0.32 | 0.32 |
| rs72653599 | 4:90286155 | A / C | 0.09 | 0.08 | 0.08 |
| rs11931992 | 4:90286319 | A / C | 0.30 | 0.30 | 0.29 |
| rs7693500 | 4:90286489 | A / G | 0.32 | 0.32 | 0.31 |
| rs72653600 | 4:90287725 | C / G | 0.09 | 0.09 | 0.08 |
| rs11943738 | 4:90288553 | G / A | 0.32 | 0.32 | 0.31 |
| rs6843814 | 4:90289197 | C / T | 0.20 | 0.21 | 0.20 |
| rs17015513 | 4:90289949 | T / A | 0.32 | 0.32 | 0.31 |
| rs28506120 | 4:90290364 | C / T | 0.32 | 0.32 | 0.31 |
| rs17777075 | 4:90291497 | T / C | 0.09 | 0.09 | 0.08 |
| rs3913682 | 4:90291730 | G / A | 0.32 | 0.32 | 0.31 |
| rs7687375 | 4:90293076 | T / C | 0.30 | 0.30 | 0.29 |
| rs2134893 | 4:90293458 | T / C | 0.32 | 0.32 | 0.31 |
| rs74398089 | 4:90293825 | C / T | 0.15 | 0.17 | 0.15 |
| rs6851534 | 4:90293893 | G / A | 0.32 | 0.32 | 0.31 |
| rs6852128 | 4:90294113 | C / A | 0.32 | 0.32 | 0.31 |
| rs2904272 | 4:90294484 | G / A | 0.32 | 0.32 | 0.31 |
| rs28653188 | 4:90294500 | A / T | 0.09 | 0.09 | 0.08 |
| rs2869997 | 4:90294503 | C / T | 0.30 | 0.30 | 0.29 |
| rs28543987 | 4:90295299 | C / T | 0.32 | 0.32 | 0.31 |
| rs1504484 | 4:90295681 | A / G | 0.20 | 0.21 | 0.20 |
| rs7677610 | 4:90296443 | T / C | 0.20 | 0.21 | 0.20 |
| rs28714620 | 4:90297075 | G / C | 0.09 | 0.08 | 0.08 |
| rs28416071 | 4:90297362 | C / T | 0.09 | 0.08 | 0.08 |
| rs1504483 | 4:90297633 | G / A | 0.12 | 0.11 | 0.11 |
| rs6843061 | 4:90297854 | A / G | 0.20 | 0.21 | 0.20 |
| rs6843085 | 4:90297887 | C / G | 0.32 | 0.32 | 0.31 |
| rs1504482 | 4:90298387 | T / C | 0.32 | 0.32 | 0.31 |
| rs6855556 | 4:90299462 | T / C | 0.20 | 0.21 | 0.20 |
| rs12331887 | 4:90301859 | A / G | 0.09 | 0.09 | 0.08 |
| rs28416334 | 4:90303741 | A / G | 0.09 | 0.09 | 0.08 |
| rs7677756 | 4:90303856 | G / A | 0.23 | 0.23 | 0.23 |
| rs72655807 | 4:90304023 | C / A | 0.09 | 0.09 | 0.08 |
| rs7662160 | 4:90304608 | G / T | 0.24 | 0.24 | 0.24 |
| rs9992323 | 4:90304918 | C / G | 0.24 | 0.24 | 0.24 |
| rs4499668 | 4:90305406 | G / A | 0.32 | 0.33 | 0.32 |
| rs10007482 | 4:90305628 | T / C | 0.32 | 0.33 | 0.32 |
| rs10461166 | 4:90305975 | A / G | 0.32 | 0.33 | 0.32 |
| rs6827974 | 4:90306233 | G / A | 0.24 | 0.24 | 0.24 |
| rs6832961 | 4:90306309 | G / A | 0.21 | 0.22 | 0.21 |
| rs1394340 | 4:90307154 | T / C | 0.24 | 0.24 | 0.24 |
| rs1827644 | 4:90307459 | C / A | 0.24 | 0.24 | 0.24 |
| rs9994884 | 4:90309082 | G / A | 0.24 | 0.24 | 0.24 |
| rs12650758 | 4:90309312 | C / T | 0.21 | 0.22 | 0.21 |
| rs17015542 | 4:90309879 | C / A | 0.21 | 0.22 | 0.21 |
| rs1105138 | 4:90310238 | C / T | 0.32 | 0.33 | 0.32 |
| rs1105137 | 4:90310346 | C / T | 0.24 | 0.24 | 0.24 |
| rs1105136 | 4:90310629 | A / G | 0.21 | 0.22 | 0.21 |
| rs1910679 | 4:90310693 | A / G | 0.21 | 0.22 | 0.21 |
| rs10001309 | 4:90311105 | G / A | 0.21 | 0.22 | 0.21 |
| rs4305484 | 4:90313373 | G / A | 0.21 | 0.22 | 0.21 |
| rs185064860 | 4:90316060 | G / C | 0.09 | 0.08 | 0.07 |
| rs113535357 | 4:90316097 | A / C | 0.09 | 0.08 | 0.07 |
| rs10026193 | 4:90317246 | T / C | 0.24 | 0.24 | 0.24 |
| rs12332003 | 4:90318322 | G / C | 0.21 | 0.22 | 0.21 |
| rs77051637 | 4:90322664 | A / C | 0.09 | 0.09 | 0.08 |
| rs141886430 | 4:90322730 | T / C | 0.09 | 0.09 | 0.08 |
| rs72655811 | 4:90323888 | T / G | 0.09 | 0.09 | 0.08 |
| rs72655813 | 4:90325122 | C / T | 0.10 | 0.12 | 0.11 |
| rs72655815 | 4:90325868 | A / G | 0.09 | 0.09 | 0.08 |
| rs72655817 | 4:90326683 | A / G | 0.09 | 0.09 | 0.08 |
| rs72655818 | 4:90327069 | A / G | 0.09 | 0.09 | 0.08 |
| rs6834282 | 4:90327801 | T / A | 0.10 | 0.13 | 0.12 |
| rs56362344 | 4:90328595 | C / A | 0.09 | 0.09 | 0.08 |
| rs56329129 | 4:90328864 | G / A | 0.09 | 0.09 | 0.08 |
| rs72655820 | 4:90331517 | A / G | 0.10 | 0.10 | 0.09 |
| rs10012175 | 4:90332273 | T / C | 0.10 | 0.10 | 0.09 |
| rs10018371 | 4:90334129 | A / C | 0.09 | 0.09 | 0.08 |
| rs72655822 | 4:90334443 | G / T | 0.09 | 0.08 | 0.08 |
| rs10029711 | 4:90335696 | G / A | 0.10 | 0.10 | 0.09 |
| rs9307068 | 4:90336018 | G / A | 0.09 | 0.09 | 0.08 |
| rs10032499 | 4:90336304 | G / A | 0.09 | 0.09 | 0.08 |
| rs772480 | 4:90336965 | G / A | 0.10 | 0.13 | 0.12 |
| rs17777756 | 4:90337651 | C / T | 0.10 | 0.11 | 0.11 |
| rs17777845 | 4:90343298 | A / G | 0.09 | 0.09 | 0.08 |
| rs6840276 | 4:90344613 | A / G | 0.08 | 0.07 | 0.08 |
| rs17777909 | 4:90345503 | G / A | 0.09 | 0.09 | 0.08 |
| rs28752002 | 4:90346897 | G / A | 0.10 | 0.10 | 0.09 |
| rs35510201 | 4:90348139 | A / G | 0.10 | 0.10 | 0.09 |
| rs35563601 | 4:90350064 | T / C | 0.18 | 0.20 | 0.19 |
| rs28539869 | 4:90353105 | C / T | 0.11 | 0.10 | 0.09 |
| rs1995518 | 4:90355200 | C / T | 0.09 | 0.08 | 0.08 |
| rs12646144 | 4:90355727 | A / C | 0.48 | 0.50 | 0.50 |
| rs10027198 | 4:90359410 | C / A | 0.10 | 0.09 | 0.09 |
| rs10027199 | 4:90359411 | G / A | 0.10 | 0.09 | 0.09 |
| rs772477 | 4:90360199 | G / C | 0.36 | 0.38 | 0.36 |
| rs772478 | 4:90361318 | G / C | 0.29 | 0.33 | 0.30 |
| rs772479 | 4:90361822 | G / T | 0.29 | 0.33 | 0.30 |
| rs772481 | 4:90363246 | C / A | 0.32 | 0.37 | 0.33 |
| rs12511414 | 4:90363314 | C / T | 0.06 | 0.05 | 0.05 |
| rs772482 | 4:90363750 | C / A | 0.39 | 0.42 | 0.39 |
| rs76072948 | 4:90364101 | C / T | 0.06 | 0.05 | 0.05 |
| rs62306598 | 4:90364557 | A / C | 0.08 | 0.09 | 0.06 |
| rs772483 | 4:90365843 | A / C | 0.29 | 0.33 | 0.30 |
| rs772484 | 4:90366020 | T / C | 0.29 | 0.33 | 0.30 |
| rs4693230 | 4:90366237 | G / A | 0.06 | 0.05 | 0.05 |
| rs772485 | 4:90366387 | A / G | 0.36 | 0.38 | 0.36 |
| rs1081769 | 4:90366673 | G / A | 0.45 | 0.47 | 0.45 |
| rs772486 | 4:90366721 | G / A | 0.36 | 0.38 | 0.36 |
| rs772487 | 4:90367051 | A / T | 0.36 | 0.38 | 0.36 |
| rs75883804 | 4:90367525 | G / A | 0.06 | 0.05 | 0.05 |
| rs61625513 | 4:90367593 | A / T | 0.06 | 0.05 | 0.05 |
| rs76942491 | 4:90368407 | A / T | 0.06 | 0.05 | 0.05 |
| rs41326145 | 4:90368433 | A / G | 0.10 | 0.09 | 0.10 |
| rs772488 | 4:90368601 | T / C | 0.45 | 0.47 | 0.45 |
| rs772489 | 4:90369343 | A / C | 0.36 | 0.38 | 0.36 |
| rs1504485 | 4:90370139 | T / G | 0.36 | 0.38 | 0.36 |
| rs1504486 | 4:90370169 | A / G | 0.36 | 0.38 | 0.36 |
| rs1504487 | 4:90370229 | C / T | 0.36 | 0.38 | 0.36 |
| rs1566853 | 4:90370602 | A / G | 0.06 | 0.05 | 0.05 |
| rs772490 | 4:90371719 | G / A | 0.36 | 0.38 | 0.36 |
| rs12643268 | 4:90371779 | G / A | 0.06 | 0.05 | 0.05 |
| rs17778445 | 4:90371934 | A / G | 0.09 | 0.08 | 0.10 |
| rs2643795 | 4:90372003 | A / G | 0.29 | 0.33 | 0.30 |
| rs2655908 | 4:90372007 | T / C | 0.29 | 0.33 | 0.30 |
| rs772491 | 4:90372125 | G / C | 0.45 | 0.47 | 0.45 |
| rs772492 | 4:90372566 | T / C | 0.29 | 0.33 | 0.30 |
| rs2132286 | 4:90372775 | T / G | 0.13 | 0.16 | 0.15 |
| rs77583026 | 4:90372783 | T / G | 0.06 | 0.05 | 0.05 |
| rs772493 | 4:90372784 | C / G | 0.34 | 0.37 | 0.34 |
| rs772494 | 4:90373071 | T / C | 0.45 | 0.47 | 0.45 |
| rs772495 | 4:90373130 | A / G | 0.45 | 0.47 | 0.45 |
| rs772496 | 4:90373435 | A / G | 0.36 | 0.38 | 0.36 |
| rs772497 | 4:90373560 | G / A | 0.29 | 0.33 | 0.30 |
| rs72655840 | 4:90373815 | C / T | 0.09 | 0.08 | 0.08 |
| rs28691145 | 4:90375052 | C / T | 0.09 | 0.08 | 0.08 |
| rs772498 | 4:90375440 | C / T | 0.36 | 0.38 | 0.36 |
| rs772499 | 4:90375457 | G / A | 0.29 | 0.33 | 0.30 |
| rs772500 | 4:90375514 | C / T | 0.36 | 0.38 | 0.36 |
| rs772501 | 4:90375951 | T / C | 0.36 | 0.38 | 0.36 |
| rs772502 | 4:90376702 | C / A | 0.36 | 0.38 | 0.36 |
| rs772503 | 4:90377577 | T / C | 0.36 | 0.38 | 0.36 |
| rs2201539 | 4:90377749 | G / T | 0.13 | 0.15 | 0.14 |
| rs772504 | 4:90377991 | G / T | 0.29 | 0.33 | 0.30 |
| rs10428294 | 4:90381514 | T / C | 0.08 | 0.07 | 0.07 |
| rs12505424 | 4:90383179 | G / T | 0.32 | 0.37 | 0.35 |
| rs55997152 | 4:90383629 | C / T | 0.32 | 0.37 | 0.35 |
| rs6816800 | 4:90383688 | G / C | 0.32 | 0.37 | 0.35 |
| rs7691267 | 4:90384672 | C / T | 0.32 | 0.37 | 0.35 |
| rs7667191 | 4:90384738 | A / G | 0.32 | 0.37 | 0.35 |
| rs34666606 | 4:90385295 | G / A | 0.07 | 0.05 | 0.06 |
| rs12646051 | 4:90385491 | G / A | 0.07 | 0.05 | 0.06 |
| rs2172895 | 4:90385574 | T / G | 0.48 | 0.49 | 0.48 |
| rs6849255 | 4:90386496 | G / C | 0.16 | 0.16 | 0.16 |
| rs2220910 | 4:90387509 | G / A | 0.48 | 0.50 | 0.48 |
| rs1948520 | 4:90387594 | G / A | 0.14 | 0.12 | 0.12 |
| rs976595 | 4:90388994 | G / A | 0.35 | 0.38 | 0.37 |
| rs976594 | 4:90389181 | T / C | 0.48 | 0.50 | 0.48 |
| rs976593 | 4:90389458 | A / G | 0.05 | 0.06 | 0.06 |
| rs1495553 | 4:90389965 | C / T | 0.48 | 0.50 | 0.48 |
| rs6823126 | 4:90390282 | A / G | 0.35 | 0.38 | 0.37 |
| rs28836900 | 4:90390517 | C / G | 0.14 | 0.12 | 0.11 |
| rs6829149 | 4:90390887 | A / G | 0.48 | 0.50 | 0.48 |
| rs6811937 | 4:90391302 | C / T | 0.35 | 0.38 | 0.37 |
| rs62305079 | 4:90392000 | G / A | 0.35 | 0.38 | 0.37 |
| rs9992358 | 4:90393086 | A / G | 0.29 | 0.32 | 0.33 |
| rs17778743 | 4:90393645 | G / A | 0.48 | 0.50 | 0.48 |
| rs55782613 | 4:90394198 | A / C | 0.14 | 0.12 | 0.11 |
| rs55736949 | 4:90394202 | A / G | 0.14 | 0.12 | 0.11 |
| rs72878810 | 4:90394404 | G / C | 0.06 | 0.05 | 0.05 |
| rs9998962 | 4:90395266 | G / A | 0.48 | 0.50 | 0.48 |
| rs10033107 | 4:90395276 | A / T | 0.48 | 0.50 | 0.48 |
| rs2870010 | 4:90395312 | T / C | 0.07 | 0.04 | 0.06 |
| rs12644111 | 4:90395499 | T / C | 0.07 | 0.06 | 0.06 |
| rs28655612 | 4:90396429 | A / G | 0.07 | 0.07 | 0.06 |
| rs17015625 | 4:90396655 | A / G | 0.07 | 0.06 | 0.07 |
| rs10013527 | 4:90397339 | T / G | 0.14 | 0.13 | 0.13 |
| rs28500640 | 4:90398267 | A / G | 0.14 | 0.13 | 0.13 |
| rs28728387 | 4:90398313 | G / A | 0.08 | 0.07 | 0.06 |
| rs28396386 | 4:90398529 | T / C | 0.15 | 0.13 | 0.13 |
| rs17015631 | 4:90398856 | T / G | 0.14 | 0.13 | 0.13 |
| rs62305080 | 4:90399234 | G / C | 0.12 | 0.09 | 0.10 |
| rs28396663 | 4:90399248 | T / G | 0.15 | 0.13 | 0.13 |
| rs11723929 | 4:90399712 | G / A | 0.07 | 0.06 | 0.07 |
| rs11736586 | 4:90399763 | C / T | 0.15 | 0.13 | 0.13 |
| rs11732242 | 4:90399764 | C / G | 0.15 | 0.13 | 0.13 |
| rs28815018 | 4:90400218 | T / C | 0.07 | 0.07 | 0.06 |
| rs28875504 | 4:90402314 | T / C | 0.08 | 0.07 | 0.06 |
| rs6826153 | 4:90402934 | G / A | 0.14 | 0.13 | 0.13 |
| rs2132281 | 4:90404263 | G / A | 0.14 | 0.13 | 0.13 |
| rs2172894 | 4:90404465 | T / C | 0.15 | 0.13 | 0.13 |
| rs17835460 | 4:90404917 | A / C | 0.07 | 0.07 | 0.06 |
| rs10006627 | 4:90405700 | T / C | 0.07 | 0.06 | 0.07 |
| rs62305081 | 4:90405899 | C / T | 0.08 | 0.07 | 0.07 |
| rs62305082 | 4:90406028 | A / T | 0.12 | 0.09 | 0.10 |
| rs28610923 | 4:90406533 | C / T | 0.08 | 0.07 | 0.06 |
| rs17015642 | 4:90407017 | T / C | 0.08 | 0.07 | 0.06 |
| rs17015650 | 4:90407772 | C / T | 0.08 | 0.07 | 0.06 |
| rs11729694 | 4:90407860 | C / G | 0.07 | 0.06 | 0.07 |
| rs17015658 | 4:90408036 | C / A | 0.08 | 0.07 | 0.06 |
| rs72655848 | 4:90408670 | G / A | 0.08 | 0.07 | 0.06 |
| rs28662801 | 4:90408690 | A / C | 0.08 | 0.07 | 0.06 |
| rs58262842 | 4:90408722 | A / G | 0.07 | 0.06 | 0.07 |
| rs28572341 | 4:90408729 | C / T | 0.08 | 0.07 | 0.06 |
| rs17015665 | 4:90408959 | A / G | 0.08 | 0.07 | 0.06 |
| rs17779297 | 4:90409136 | G / C | 0.17 | 0.19 | 0.20 |
| rs6852601 | 4:90409844 | G / A | 0.08 | 0.07 | 0.06 |
| rs28379423 | 4:90409931 | T / C | 0.07 | 0.07 | 0.06 |
| rs6852971 | 4:90409946 | T / C | 0.08 | 0.07 | 0.06 |
| rs6830052 | 4:90409979 | C / T | 0.15 | 0.13 | 0.13 |
| rs17015675 | 4:90410293 | T / A | 0.08 | 0.07 | 0.06 |
| rs17015679 | 4:90410367 | A / C | 0.08 | 0.07 | 0.06 |
| rs28432001 | 4:90410542 | T / A | 0.21 | 0.16 | 0.17 |
| rs12648284 | 4:90410551 | A / G | 0.06 | 0.06 | 0.06 |
| rs12648334 | 4:90410754 | A / G | 0.07 | 0.06 | 0.07 |
| rs115539802 | 4:90411254 | T / C | 0.08 | 0.07 | 0.06 |
| rs28627588 | 4:90412279 | A / G | 0.08 | 0.07 | 0.06 |
| rs143372373 | 4:90412386 | A / G | 0.05 | 0.07 | 0.06 |
| rs112744012 | 4:90412435 | G / A | 0.10 | 0.09 | 0.10 |
| rs3796669 | 4:90412734 | C / T | 0.07 | 0.07 | 0.07 |
| rs3796670 | 4:90412880 | A / G | 0.15 | 0.13 | 0.13 |
| rs62305085 | 4:90414415 | G / A | 0.07 | 0.07 | 0.07 |
| rs28463872 | 4:90414445 | A / G | 0.15 | 0.13 | 0.13 |
| rs62305086 | 4:90414462 | G / A | 0.07 | 0.07 | 0.07 |
| rs11934629 | 4:90414925 | C / T | 0.07 | 0.07 | 0.07 |
| rs11729660 | 4:90415267 | G / A | 0.07 | 0.07 | 0.07 |
| rs72655852 | 4:90415385 | T / G | 0.34 | 0.39 | 0.37 |
| rs10025166 | 4:90417200 | T / C | 0.07 | 0.07 | 0.07 |
| rs17174961 | 4:90418192 | A / G | 0.34 | 0.39 | 0.37 |
| rs1908557 | 4:90421353 | C / T | 0.21 | 0.19 | 0.17 |
| rs10017959 | 4:90421644 | C / A | 0.21 | 0.19 | 0.17 |
| rs1908556 | 4:90421700 | G / A | 0.14 | 0.15 | 0.16 |
| rs10008041 | 4:90422209 | C / T | 0.14 | 0.12 | 0.10 |
| rs11732740 | 4:90423029 | G / A | 0.21 | 0.19 | 0.17 |
| rs17779630 | 4:90423385 | G / A | 0.08 | 0.06 | 0.06 |
| rs17015692 | 4:90424103 | T / C | 0.07 | 0.08 | 0.07 |
| rs17015698 | 4:90425675 | T / C | 0.07 | 0.06 | 0.07 |
| rs13131907 | 4:90425944 | C / T | 0.45 | 0.45 | 0.44 |
| rs62305089 | 4:90426185 | T / C | 0.07 | 0.06 | 0.07 |
| rs6824131 | 4:90426307 | T / C | 0.14 | 0.16 | 0.17 |
| rs17015705 | 4:90426646 | G / A | 0.07 | 0.06 | 0.07 |
| rs903601 | 4:90427876 | A / T | 0.07 | 0.07 | 0.07 |
| rs13119598 | 4:90428054 | A / G | 0.45 | 0.46 | 0.44 |
| rs17175345 | 4:90428534 | T / A | 0.06 | 0.06 | 0.06 |
| rs1495552 | 4:90429040 | T / G | 0.47 | 0.48 | 0.46 |
| rs1495551 | 4:90429082 | A / G | 0.47 | 0.48 | 0.46 |
| rs17175401 | 4:90429676 | G / T | 0.08 | 0.06 | 0.06 |
| rs11734628 | 4:90429792 | G / A | 0.07 | 0.06 | 0.07 |
| rs1874345 | 4:90430568 | C / T | 0.28 | 0.28 | 0.28 |
| rs903600 | 4:90430695 | T / C | 0.28 | 0.28 | 0.28 |
| rs903598 | 4:90430770 | T / C | 0.25 | 0.24 | 0.25 |
| rs10032446 | 4:90431371 | A / G | 0.28 | 0.28 | 0.28 |
| rs13146919 | 4:90431418 | T / C | 0.35 | 0.38 | 0.36 |
| rs72655858 | 4:90431579 | G / A | 0.06 | 0.07 | 0.07 |
| rs12504969 | 4:90431635 | C / T | 0.28 | 0.28 | 0.28 |
| rs10032889 | 4:90431888 | A / C | 0.28 | 0.28 | 0.28 |
| rs2870009 | 4:90431966 | A / G | 0.28 | 0.28 | 0.28 |
| rs17175612 | 4:90432177 | T / C | 0.06 | 0.05 | 0.06 |
| rs903597 | 4:90432229 | A / G | 0.28 | 0.28 | 0.28 |
| rs903596 | 4:90432417 | T / G | 0.28 | 0.28 | 0.28 |
| rs903595 | 4:90432462 | T / C | 0.28 | 0.28 | 0.28 |
| rs2008787 | 4:90433049 | T / C | 0.08 | 0.06 | 0.06 |
| rs6823702 | 4:90433359 | T / C | 0.25 | 0.24 | 0.25 |
| rs17175738 | 4:90433679 | C / A | 0.08 | 0.06 | 0.06 |
| rs6854472 | 4:90434672 | T / G | 0.40 | 0.41 | 0.39 |
| rs729686 | 4:90435011 | G / A | 0.42 | 0.40 | 0.40 |
| rs729685 | 4:90435274 | C / T | 0.32 | 0.32 | 0.33 |
| rs924032 | 4:90435483 | G / A | 0.32 | 0.32 | 0.33 |
| rs924033 | 4:90435553 | G / T | 0.08 | 0.07 | 0.06 |
| rs7667895 | 4:90435863 | G / A | 0.27 | 0.26 | 0.28 |
| rs924034 | 4:90435865 | A / G | 0.33 | 0.33 | 0.33 |
| rs11942435 | 4:90437149 | T / C | 0.32 | 0.32 | 0.33 |
| rs1603251 | 4:90438365 | G / A | 0.32 | 0.32 | 0.33 |
| rs28814315 | 4:90438536 | C / G | 0.40 | 0.41 | 0.38 |
| rs4694007 | 4:90439656 | A / G | 0.33 | 0.33 | 0.33 |
| rs17015738 | 4:90439694 | T / C | 0.32 | 0.34 | 0.32 |
| rs11731934 | 4:90439763 | G / T | 0.32 | 0.32 | 0.33 |
| rs17015739 | 4:90439809 | G / A | 0.40 | 0.41 | 0.38 |
| rs11723738 | 4:90440083 | A / T | 0.33 | 0.32 | 0.33 |
| rs57946672 | 4:90440510 | A / G | 0.32 | 0.34 | 0.32 |
| rs17015743 | 4:90440916 | A / C | 0.40 | 0.41 | 0.38 |
| rs71609571 | 4:90440951 | T / C | 0.05 | 0.05 | 0.05 |
| rs7679614 | 4:90441114 | C / T | 0.32 | 0.34 | 0.32 |
| rs7654775 | 4:90441186 | G / A | 0.32 | 0.34 | 0.32 |
| rs990695 | 4:90441242 | A / G | 0.21 | 0.19 | 0.20 |
| rs7660434 | 4:90441444 | T / G | 0.32 | 0.34 | 0.32 |
| rs7660467 | 4:90441487 | A / G | 0.32 | 0.34 | 0.32 |
| rs11097225 | 4:90441824 | G / T | 0.32 | 0.32 | 0.33 |
| rs9790733 | 4:90441904 | A / G | 0.32 | 0.34 | 0.32 |
| rs2172893 | 4:90442287 | A / C | 0.08 | 0.09 | 0.09 |
| rs28655128 | 4:90443169 | C / T | 0.32 | 0.34 | 0.32 |
| rs72880655 | 4:90445466 | G / C | 0.32 | 0.34 | 0.32 |
| rs72880657 | 4:90445743 | C / T | 0.32 | 0.34 | 0.32 |
| rs72880660 | 4:90445966 | G / T | 0.32 | 0.34 | 0.32 |
| rs10021688 | 4:90446005 | T / C | 0.08 | 0.07 | 0.06 |
| rs55810953 | 4:90446268 | C / T | 0.32 | 0.34 | 0.32 |
| rs4694009 | 4:90446331 | T / C | 0.06 | 0.07 | 0.07 |
| rs10001744 | 4:90446427 | G / A | 0.08 | 0.07 | 0.06 |
| rs114699154 | 4:90446671 | A / G | 0.09 | 0.09 | 0.10 |
| rs61534811 | 4:90446994 | T / C | 0.32 | 0.34 | 0.32 |
| rs79657661 | 4:90447361 | G / A | 0.21 | 0.19 | 0.20 |
| rs9790381 | 4:90448105 | C / T | 0.32 | 0.34 | 0.32 |
| rs9790532 | 4:90448379 | A / G | 0.33 | 0.33 | 0.33 |
| rs10008137 | 4:90448417 | C / A | 0.27 | 0.26 | 0.28 |
| rs9790800 | 4:90448514 | A / C | 0.32 | 0.34 | 0.32 |
| rs9790803 | 4:90448639 | A / G | 0.32 | 0.34 | 0.32 |
| rs11946217 | 4:90448771 | A / G | 0.32 | 0.34 | 0.32 |
| rs11736388 | 4:90449070 | A / G | 0.35 | 0.33 | 0.34 |
| rs11937181 | 4:90449155 | G / A | 0.32 | 0.34 | 0.32 |
| rs56958658 | 4:90449607 | A / G | 0.32 | 0.34 | 0.32 |
| rs13104727 | 4:90450382 | T / A | 0.27 | 0.26 | 0.28 |
| rs4694010 | 4:90450579 | C / T | 0.06 | 0.07 | 0.07 |
| rs56402113 | 4:90450652 | T / C | 0.32 | 0.34 | 0.32 |
| rs9992832 | 4:90451094 | A / G | 0.08 | 0.07 | 0.06 |
| rs2055174 | 4:90451656 | T / G | 0.08 | 0.08 | 0.09 |
| rs6850584 | 4:90452160 | T / C | 0.25 | 0.24 | 0.26 |
| rs13114147 | 4:90452280 | C / A | 0.05 | 0.05 | 0.05 |
| rs67262058 | 4:90453241 | G / A | 0.31 | 0.29 | 0.33 |
| rs35175911 | 4:90453601 | C / T | 0.16 | 0.15 | 0.17 |
| rs13128501 | 4:90453866 | C / G | 0.31 | 0.29 | 0.33 |
| rs10002374 | 4:90454284 | T / C | 0.38 | 0.36 | 0.39 |
| rs4694011 | 4:90454945 | C / T | 0.07 | 0.09 | 0.09 |
| rs4694012 | 4:90455143 | A / G | 0.23 | 0.23 | 0.23 |
| rs72655869 | 4:90455344 | G / A | 0.06 | 0.06 | 0.06 |
| rs1495554 | 4:90455699 | T / G | 0.38 | 0.40 | 0.38 |
| rs6532175 | 4:90456406 | C / T | 0.38 | 0.40 | 0.38 |
| rs1603253 | 4:90456589 | C / T | 0.07 | 0.07 | 0.06 |
| rs1603254 | 4:90456596 | T / C | 0.07 | 0.07 | 0.06 |
| rs1390283 | 4:90457235 | T / C | 0.39 | 0.37 | 0.40 |
| rs10516838 | 4:90458232 | A / G | 0.07 | 0.07 | 0.06 |
| rs1495556 | 4:90458283 | T / C | 0.32 | 0.30 | 0.34 |
| rs17015756 | 4:90458351 | G / A | 0.07 | 0.07 | 0.06 |
| rs1845698 | 4:90459225 | A / C | 0.23 | 0.23 | 0.23 |
| rs17176388 | 4:90460511 | T / C | 0.33 | 0.36 | 0.33 |
| rs4147250 | 4:90461313 | C / T | 0.38 | 0.40 | 0.38 |
| rs721136 | 4:90461780 | G / C | 0.34 | 0.36 | 0.33 |
| rs13148501 | 4:90462203 | C / T | 0.23 | 0.23 | 0.23 |
| rs12505194 | 4:90462267 | A / G | 0.32 | 0.30 | 0.34 |
| rs17015768 | 4:90462379 | C / A | 0.34 | 0.36 | 0.33 |
| rs28402865 | 4:90462833 | C / T | 0.38 | 0.40 | 0.38 |
| rs28898346 | 4:90463643 | G / A | 0.34 | 0.36 | 0.33 |
| rs72655874 | 4:90463836 | T / C | 0.33 | 0.36 | 0.33 |
| rs72655875 | 4:90464423 | T / C | 0.33 | 0.36 | 0.33 |
| rs12511270 | 4:90465953 | G / A | 0.39 | 0.37 | 0.39 |
| rs12507716 | 4:90466011 | G / T | 0.39 | 0.37 | 0.39 |
| rs7668310 | 4:90466166 | C / A | 0.34 | 0.37 | 0.33 |
| rs7669026 | 4:90466289 | C / G | 0.38 | 0.40 | 0.38 |
| rs10034469 | 4:90467329 | C / A | 0.34 | 0.37 | 0.33 |
| rs10012532 | 4:90467471 | T / C | 0.38 | 0.40 | 0.38 |
| rs7681582 | 4:90468198 | T / A | 0.39 | 0.37 | 0.40 |
| rs4694013 | 4:90468824 | C / G | 0.23 | 0.23 | 0.22 |
| rs11097226 | 4:90469131 | G / A | 0.39 | 0.37 | 0.40 |
| rs12512901 | 4:90470497 | C / T | 0.07 | 0.07 | 0.06 |
| rs1973409 | 4:90470530 | C / T | 0.23 | 0.23 | 0.22 |
| rs13122379 | 4:90470550 | G / A | 0.32 | 0.30 | 0.34 |
| rs35643643 | 4:90470553 | A / G | 0.32 | 0.30 | 0.34 |
| rs10034176 | 4:90471033 | T / C | 0.23 | 0.23 | 0.22 |
| rs2870004 | 4:90471245 | T / A | 0.23 | 0.23 | 0.23 |
| rs1552970 | 4:90473225 | A / G | 0.43 | 0.41 | 0.44 |
| rs1124015 | 4:90473234 | C / G | 0.43 | 0.41 | 0.44 |
| rs1552971 | 4:90473282 | T / G | 0.44 | 0.41 | 0.44 |
| rs1552972 | 4:90473540 | G / A | 0.44 | 0.41 | 0.44 |
| rs1552973 | 4:90473624 | A / G | 0.44 | 0.41 | 0.44 |
| rs1552974 | 4:90473728 | T / C | 0.39 | 0.37 | 0.40 |
| rs6857779 | 4:90474291 | C / T | 0.32 | 0.30 | 0.34 |
| rs4694014 | 4:90475519 | A / G | 0.07 | 0.06 | 0.06 |
| rs9307072 | 4:90477005 | G / A | 0.33 | 0.36 | 0.33 |
| rs6841552 | 4:90477090 | T / C | 0.23 | 0.23 | 0.23 |
| rs28413036 | 4:90477360 | G / A | 0.34 | 0.37 | 0.33 |
| rs115533832 | 4:90477494 | C / G | 0.07 | 0.06 | 0.06 |
| rs62305118 | 4:90477541 | A / G | 0.08 | 0.07 | 0.07 |
| rs28449583 | 4:90477662 | T / A | 0.33 | 0.36 | 0.33 |
| rs12500488 | 4:90478334 | C / T | 0.07 | 0.06 | 0.06 |
| rs12500582 | 4:90478608 | C / T | 0.07 | 0.06 | 0.06 |
| rs2869998 | 4:90479087 | C / A | 0.23 | 0.23 | 0.23 |
| rs1603252 | 4:90479517 | A / G | 0.33 | 0.36 | 0.33 |
| rs6813115 | 4:90479725 | G / T | 0.35 | 0.37 | 0.34 |
| rs6836059 | 4:90479949 | A / G | 0.23 | 0.23 | 0.23 |
| rs1542810 | 4:90480224 | C / T | 0.23 | 0.23 | 0.23 |
| rs7685009 | 4:90480831 | G / A | 0.33 | 0.36 | 0.33 |
| rs17015809 | 4:90481590 | C / G | 0.34 | 0.37 | 0.33 |
| rs4693233 | 4:90481621 | G / C | 0.42 | 0.40 | 0.43 |
| rs1390279 | 4:90482677 | T / C | 0.23 | 0.23 | 0.23 |
| rs989057 | 4:90482945 | G / C | 0.39 | 0.36 | 0.39 |
| rs62305119 | 4:90483780 | T / C | 0.11 | 0.09 | 0.09 |
| rs1826739 | 4:90484613 | T / G | 0.23 | 0.23 | 0.23 |
| rs72655887 | 4:90485484 | G / A | 0.33 | 0.36 | 0.33 |
| rs2172890 | 4:90485959 | C / T | 0.23 | 0.23 | 0.23 |
| rs28800226 | 4:90486463 | C / T | 0.33 | 0.36 | 0.33 |
| rs6847090 | 4:90486507 | C / T | 0.23 | 0.23 | 0.23 |
| rs112179934 | 4:90488104 | A / C | 0.33 | 0.36 | 0.33 |
| rs72655890 | 4:90488379 | T / C | 0.33 | 0.36 | 0.33 |
| rs56259843 | 4:90488681 | C / T | 0.33 | 0.36 | 0.33 |
| rs6839240 | 4:90489179 | T / C | 0.23 | 0.23 | 0.23 |
| rs74498181 | 4:90489280 | A / G | 0.06 | 0.06 | 0.06 |
| rs6819340 | 4:90489459 | A / G | 0.42 | 0.40 | 0.43 |
| rs6818842 | 4:90489481 | A / G | 0.23 | 0.23 | 0.23 |
| rs76797874 | 4:90489660 | G / T | 0.06 | 0.06 | 0.06 |
| rs2870005 | 4:90491767 | A / T | 0.23 | 0.23 | 0.23 |
| rs1156115 | 4:90492766 | C / T | 0.23 | 0.23 | 0.23 |
| rs72655891 | 4:90493028 | G / A | 0.05 | 0.06 | 0.05 |
| rs4694015 | 4:90493172 | G / A | 0.23 | 0.23 | 0.23 |
| rs66539107 | 4:90493437 | G / A | 0.33 | 0.36 | 0.33 |
| rs59458976 | 4:90493729 | G / A | 0.06 | 0.06 | 0.06 |
| rs28676330 | 4:90494004 | C / T | 0.33 | 0.36 | 0.33 |
| rs1495557 | 4:90494272 | A / G | 0.34 | 0.37 | 0.34 |
| rs2870006 | 4:90495946 | G / A | 0.42 | 0.40 | 0.43 |
| rs10024210 | 4:90497449 | C / T | 0.33 | 0.36 | 0.33 |
| rs10012805 | 4:90497642 | C / G | 0.33 | 0.36 | 0.33 |
| rs2036081 | 4:90498133 | T / C | 0.23 | 0.23 | 0.23 |
| rs7693262 | 4:90499392 | G / T | 0.23 | 0.23 | 0.23 |
| rs10018847 | 4:90499545 | T / C | 0.33 | 0.36 | 0.33 |
| rs6532176 | 4:90500400 | C / G | 0.36 | 0.38 | 0.36 |
| rs2904280 | 4:90500460 | T / C | 0.13 | 0.12 | 0.13 |
| rs2870007 | 4:90500952 | C / T | 0.10 | 0.11 | 0.11 |
| rs4502649 | 4:90500977 | T / A | 0.12 | 0.12 | 0.11 |
| rs72655897 | 4:90501506 | G / A | 0.12 | 0.11 | 0.11 |
| rs74288556 | 4:90501532 | T / C | 0.12 | 0.11 | 0.11 |
| rs9985581 | 4:90503064 | C / T | 0.18 | 0.16 | 0.16 |
| rs79483792 | 4:90503661 | C / T | 0.07 | 0.07 | 0.06 |
| rs17015858 | 4:90505568 | T / C | 0.12 | 0.11 | 0.11 |
| rs1390282 | 4:90506095 | A / G | 0.18 | 0.16 | 0.16 |
| rs9994582 | 4:90506129 | A / T | 0.18 | 0.16 | 0.16 |
| rs12643604 | 4:90506168 | G / A | 0.06 | NA | 0.05 |
| rs10007471 | 4:90506334 | G / A | 0.18 | 0.16 | 0.16 |
| rs10516839 | 4:90508340 | G / T | 0.19 | 0.24 | 0.22 |
| rs6842271 | 4:90509045 | T / C | 0.18 | 0.16 | 0.16 |
| rs12507346 | 4:90509880 | G / A | 0.18 | 0.16 | 0.16 |
| rs12108375 | 4:90510214 | A / T | 0.18 | 0.16 | 0.16 |
| rs76759470 | 4:90511396 | T / C | 0.06 | 0.06 | 0.06 |
| rs1495549 | 4:90511673 | G / A | 0.18 | 0.16 | 0.16 |
| rs1390281 | 4:90512099 | A / G | 0.14 | 0.12 | 0.12 |
| rs115269995 | 4:90512990 | A / G | 0.06 | 0.06 | 0.06 |
| rs7694753 | 4:90513676 | G / T | 0.23 | 0.21 | 0.23 |
| rs79517939 | 4:90513701 | A / G | 0.05 | 0.07 | 0.06 |
| rs1603250 | 4:90513906 | T / C | 0.18 | 0.16 | 0.16 |
| rs11097227 | 4:90516308 | C / T | 0.18 | 0.16 | 0.16 |
| rs1473533 | 4:90516679 | A / G | 0.06 | 0.06 | 0.06 |
| rs1473532 | 4:90516766 | A / T | 0.06 | 0.06 | 0.06 |
| rs11097228 | 4:90517413 | G / A | 0.18 | 0.16 | 0.16 |
| rs1390280 | 4:90519452 | A / G | 0.18 | 0.16 | 0.16 |
| rs114061474 | 4:90521460 | T / C | 0.06 | 0.07 | 0.06 |
| rs9993835 | 4:90521832 | C / A | 0.18 | 0.16 | 0.16 |
| rs6852411 | 4:90522480 | A / C | 0.18 | 0.16 | 0.16 |
| rs9992729 | 4:90524680 | C / T | 0.18 | 0.16 | 0.16 |
| rs9993107 | 4:90525023 | C / T | 0.18 | 0.16 | 0.16 |
| rs1121173 | 4:90525808 | C / T | 0.18 | 0.16 | 0.16 |
| rs76529608 | 4:90526433 | T / A | 0.07 | 0.07 | 0.06 |
| rs6850389 | 4:90527344 | A / G | 0.07 | 0.06 | 0.06 |
| rs6849913 | 4:90527394 | G / A | 0.07 | 0.06 | 0.06 |
| rs77322437 | 4:90527432 | T / C | 0.07 | 0.07 | 0.06 |
| rs6850616 | 4:90527474 | A / G | 0.07 | 0.06 | 0.06 |
| rs6827771 | 4:90527647 | A / T | 0.18 | 0.16 | 0.16 |
| rs73830663 | 4:90527762 | A / G | 0.07 | 0.07 | 0.06 |
| rs17015895 | 4:90527778 | C / T | 0.07 | 0.07 | 0.06 |
| rs73830682 | 4:90528218 | G / T | 0.07 | 0.07 | 0.06 |
| rs17015896 | 4:90529920 | G / C | 0.07 | 0.07 | 0.06 |
| rs17015898 | 4:90529991 | T / C | 0.19 | 0.16 | 0.16 |
| rs73830685 | 4:90530736 | A / G | 0.07 | 0.07 | 0.06 |
| rs111852332 | 4:90531263 | A / G | 0.07 | 0.07 | 0.06 |
| rs6855067 | 4:90531550 | C / T | 0.19 | 0.16 | 0.16 |
| rs116431061 | 4:90531586 | C / G | 0.07 | 0.07 | 0.06 |
| rs112988132 | 4:90531829 | G / C | 0.07 | 0.07 | 0.06 |
| rs139466426 | 4:90532129 | G / A | 0.07 | 0.07 | 0.06 |
| rs75017686 | 4:90532879 | A / G | 0.08 | 0.06 | 0.06 |
| rs11737634 | 4:90533049 | T / C | 0.07 | 0.07 | 0.06 |
| rs11722151 | 4:90533454 | A / G | 0.07 | 0.07 | 0.06 |
| rs77211998 | 4:90533529 | C / A | 0.07 | 0.07 | 0.06 |
| rs75878722 | 4:90534463 | G / A | 0.07 | 0.07 | 0.06 |
| rs75152640 | 4:90534895 | A / G | 0.07 | 0.07 | 0.06 |
| rs1036084 | 4:90535487 | G / T | 0.19 | 0.16 | 0.16 |
| rs73830688 | 4:90535601 | T / C | 0.07 | 0.07 | 0.06 |
| rs73830689 | 4:90536733 | C / T | 0.07 | 0.07 | 0.06 |
| rs79371144 | 4:90537581 | A / T | 0.07 | 0.07 | 0.06 |
| rs73830690 | 4:90537790 | T / C | 0.07 | 0.07 | 0.06 |
| rs7683607 | 4:90538643 | T / C | 0.19 | 0.17 | 0.16 |
| rs73830692 | 4:90539151 | A / G | 0.07 | 0.07 | 0.06 |
| rs17015906 | 4:90541469 | C / T | 0.07 | 0.07 | 0.06 |
| rs17015911 | 4:90541803 | C / A | 0.07 | 0.07 | 0.06 |
| rs61648356 | 4:90542174 | A / C | 0.08 | 0.06 | 0.06 |
| rs79640052 | 4:90542194 | T / C | 0.08 | 0.06 | 0.06 |
| rs58310666 | 4:90543092 | T / C | 0.08 | 0.06 | 0.06 |
| rs2163392 | 4:90543684 | G / A | 0.07 | 0.07 | 0.06 |
| rs74636730 | 4:90543882 | G / C | 0.07 | 0.07 | 0.06 |
| rs17015916 | 4:90544349 | G / T | 0.07 | 0.07 | 0.06 |
| rs111873607 | 4:90544629 | G / C | 0.06 | 0.07 | 0.06 |
| rs11728943 | 4:90545039 | T / G | 0.06 | 0.07 | 0.06 |
| rs11737155 | 4:90545062 | G / A | 0.06 | 0.07 | 0.06 |
| rs11728999 | 4:90545192 | A / G | 0.06 | 0.07 | 0.06 |
| rs13125108 | 4:90545213 | G / T | 0.49 | 0.43 | 0.46 |
| rs11728970 | 4:90545239 | T / C | 0.06 | 0.07 | 0.06 |
| rs11723723 | 4:90545298 | C / G | 0.06 | 0.07 | 0.06 |
| rs11723725 | 4:90545315 | A / G | 0.06 | 0.07 | 0.06 |
| rs75754323 | 4:90545492 | A / T | 0.06 | 0.07 | 0.06 |
| rs79208813 | 4:90545495 | T / C | 0.06 | 0.07 | 0.06 |
| rs10031817 | 4:90549446 | C / A | 0.14 | 0.12 | 0.12 |
| rs10027842 | 4:90551543 | C / T | 0.25 | 0.21 | 0.21 |
| rs1430961 | 4:90552920 | G / A | 0.09 | 0.09 | 0.08 |
| rs1430960 | 4:90553180 | A / G | 0.09 | 0.09 | 0.08 |
| rs12498987 | 4:90554975 | A / T | 0.08 | 0.05 | 0.06 |
| rs12513009 | 4:90555520 | G / A | 0.07 | 0.05 | 0.06 |
| rs10012262 | 4:90555837 | T / C | 0.17 | 0.15 | 0.15 |
| rs11933399 | 4:90559006 | G / A | 0.09 | 0.09 | 0.08 |
| rs9992254 | 4:90560206 | C / T | 0.07 | 0.05 | 0.06 |
| rs16996194 | 4:90560529 | A / C | 0.09 | 0.09 | 0.08 |
| rs10011947 | 4:90562801 | G / A | 0.22 | 0.18 | 0.19 |
| rs17015961 | 4:90563398 | G / C | 0.08 | 0.05 | 0.06 |
| rs76142252 | 4:90564400 | C / T | 0.09 | 0.09 | 0.08 |
| rs17015968 | 4:90565207 | A / G | 0.07 | 0.05 | 0.06 |
| rs11932021 | 4:90565218 | T / C | 0.09 | 0.09 | 0.08 |
| rs11944530 | 4:90565615 | A / C | 0.09 | 0.09 | 0.08 |
| rs57856896 | 4:90566339 | T / C | 0.08 | 0.05 | 0.06 |
| rs13140923 | 4:90566469 | A / G | 0.21 | 0.19 | 0.21 |
| rs28498646 | 4:90566912 | A / G | 0.07 | 0.05 | 0.06 |
| rs77431271 | 4:90567048 | A / G | 0.09 | 0.09 | 0.08 |
| rs189924866 | 4:90569158 | G / A | 0.17 | 0.15 | 0.15 |
| rs146632109 | 4:90569619 | C / T | 0.21 | 0.19 | 0.18 |
| rs72657705 | 4:90572277 | T / A | 0.19 | 0.26 | 0.23 |
| rs3733449 | 4:90572322 | T / C | 0.21 | 0.19 | 0.18 |
| rs78553492 | 4:90573045 | A / C | 0.09 | 0.09 | 0.08 |
| rs28517275 | 4:90573173 | C / T | 0.07 | 0.05 | 0.06 |
| rs356231 | 4:90573396 | G / A | 0.04 | 0.05 | 0.06 |
| rs3846288 | 4:90574806 | A / C | 0.07 | 0.05 | 0.06 |
| rs11933687 | 4:90576512 | T / C | 0.09 | 0.09 | 0.08 |
| rs28673968 | 4:90576890 | C / T | 0.21 | 0.19 | 0.18 |
| rs72657707 | 4:90578511 | A / C | 0.20 | 0.26 | 0.23 |
| rs7674594 | 4:90579508 | A / G | 0.11 | 0.09 | 0.10 |
| rs7674641 | 4:90579586 | A / G | 0.11 | 0.09 | 0.10 |
| rs28545331 | 4:90581017 | A / T | 0.21 | 0.19 | 0.18 |
| rs62305131 | 4:90581986 | G / A | 0.08 | 0.07 | 0.07 |
| rs78786660 | 4:90583076 | C / T | 0.09 | 0.09 | 0.08 |
| rs62305132 | 4:90585132 | G / A | 0.08 | 0.07 | 0.07 |
| rs10018310 | 4:90587691 | G / C | 0.11 | 0.09 | 0.09 |
| rs17015973 | 4:90589362 | G / A | 0.08 | 0.07 | 0.07 |
| rs9986016 | 4:90590406 | C / G | 0.11 | 0.09 | 0.09 |
| rs9994859 | 4:90590704 | C / T | 0.11 | 0.09 | 0.09 |
| rs60443785 | 4:90591180 | C / G | 0.08 | 0.07 | 0.07 |
| rs60043746 | 4:90591239 | G / A | 0.08 | 0.07 | 0.07 |
| rs60639576 | 4:90591283 | G / A | 0.08 | 0.07 | 0.07 |
| rs58123417 | 4:90591376 | G / C | 0.08 | 0.07 | 0.07 |
| rs59130061 | 4:90591397 | C / T | 0.08 | 0.07 | 0.07 |
| rs56098452 | 4:90591552 | C / T | 0.08 | 0.07 | 0.07 |
| rs9985612 | 4:90591889 | C / T | 0.20 | 0.18 | 0.17 |
| rs9986065 | 4:90592039 | T / C | 0.11 | 0.09 | 0.09 |
| rs61631234 | 4:90592262 | A / C | 0.08 | 0.07 | 0.07 |
| rs58992864 | 4:90592311 | G / A | 0.08 | 0.07 | 0.07 |
| rs59580572 | 4:90592345 | C / T | 0.08 | 0.07 | 0.07 |
| rs57360006 | 4:90592448 | G / A | 0.08 | 0.07 | 0.07 |
| rs61528967 | 4:90592455 | T / C | 0.08 | 0.07 | 0.07 |
| rs62305143 | 4:90592544 | A / C | 0.08 | 0.07 | 0.07 |
| rs61253650 | 4:90592551 | C / T | 0.08 | 0.07 | 0.07 |
| rs6835887 | 4:90592695 | C / A | 0.20 | 0.18 | 0.17 |
| rs56389733 | 4:90592856 | A / G | 0.08 | 0.07 | 0.07 |
| rs58951429 | 4:90592857 | T / C | 0.08 | 0.07 | 0.07 |
| rs60954064 | 4:90592962 | G / A | 0.11 | 0.09 | 0.09 |
| rs57864032 | 4:90593212 | C / T | 0.08 | 0.07 | 0.07 |
| rs56970479 | 4:90593251 | T / A | 0.08 | 0.07 | 0.07 |
| rs62305148 | 4:90593447 | C / A | 0.08 | 0.07 | 0.07 |
| rs62305149 | 4:90593479 | A / C | 0.08 | 0.07 | 0.07 |
| rs62305150 | 4:90593545 | G / A | 0.08 | 0.07 | 0.07 |
| rs62305151 | 4:90593708 | G / T | 0.08 | 0.07 | 0.07 |
| rs62305152 | 4:90593804 | G / A | 0.08 | 0.07 | 0.07 |
| rs62305153 | 4:90593895 | T / C | 0.08 | 0.07 | 0.07 |
| rs62305154 | 4:90593991 | T / C | 0.08 | 0.07 | 0.07 |
| rs62305155 | 4:90594084 | T / C | 0.08 | 0.07 | 0.07 |
| rs113525047 | 4:90594387 | G / A | 0.08 | 0.07 | 0.07 |
| rs112657084 | 4:90594444 | C / G | 0.08 | 0.07 | 0.07 |
| rs9685020 | 4:90594466 | G / A | 0.08 | 0.07 | 0.07 |
| rs144938752 | 4:90594725 | G / C | 0.08 | 0.07 | 0.07 |
| rs112249490 | 4:90594929 | C / T | 0.08 | 0.07 | 0.07 |
| rs113895591 | 4:90594942 | A / G | 0.08 | 0.07 | 0.07 |
| rs28871018 | 4:90594987 | A / G | 0.21 | 0.18 | 0.19 |
| rs113764237 | 4:90594996 | C / T | 0.08 | 0.07 | 0.07 |
| rs62305156 | 4:90595607 | T / C | 0.08 | 0.07 | 0.07 |
| rs62305157 | 4:90595639 | A / G | 0.08 | 0.07 | 0.07 |
| rs62305158 | 4:90595817 | G / A | 0.08 | 0.07 | 0.07 |
| rs28881549 | 4:90595940 | C / T | 0.20 | 0.18 | 0.17 |
| rs72884405 | 4:90596153 | T / G | 0.08 | 0.07 | 0.07 |
| rs4122859 | 4:90596176 | C / T | 0.08 | 0.07 | 0.07 |
| rs4122858 | 4:90596515 | C / A | 0.08 | 0.07 | 0.07 |
| rs4449381 | 4:90596628 | G / A | 0.08 | 0.07 | 0.07 |
| rs2882783 | 4:90596890 | T / A | 0.08 | 0.07 | 0.07 |
| rs356230 | 4:90596923 | C / G | 0.21 | 0.27 | 0.24 |
| rs2351861 | 4:90597049 | C / G | 0.08 | 0.07 | 0.07 |
| rs7437342 | 4:90597132 | G / A | 0.08 | 0.07 | 0.07 |
| rs112150218 | 4:90597600 | C / T | 0.08 | 0.07 | 0.07 |
| rs11724963 | 4:90597736 | A / T | 0.11 | 0.09 | 0.09 |
| rs62306262 | 4:90598165 | T / A | 0.08 | 0.07 | 0.07 |
| rs62306263 | 4:90598175 | G / A | 0.08 | 0.07 | 0.07 |
| rs7682447 | 4:90598311 | C / A | 0.08 | 0.07 | 0.07 |
| rs7683289 | 4:90598763 | C / A | 0.11 | 0.09 | 0.09 |
| rs112033215 | 4:90599069 | T / C | 0.08 | 0.07 | 0.07 |
| rs10213062 | 4:90600620 | C / T | 0.11 | 0.09 | 0.09 |
| rs3857046 | 4:90600771 | A / G | 0.08 | 0.06 | 0.07 |
| rs7697331 | 4:90600999 | A / G | 0.10 | 0.09 | 0.08 |
| rs11942911 | 4:90601310 | G / A | 0.10 | 0.09 | 0.08 |
| rs7656703 | 4:90601889 | G / A | 0.08 | 0.06 | 0.07 |
| rs13108925 | 4:90602795 | C / T | 0.21 | 0.20 | 0.21 |
| rs17015982 | 4:90603044 | G / A | 0.08 | 0.06 | 0.07 |
| rs17015985 | 4:90603210 | A / G | 0.08 | 0.06 | 0.07 |
| rs12644119 | 4:90603419 | A / C | 0.12 | 0.12 | 0.10 |
| rs17015990 | 4:90604322 | T / C | 0.08 | 0.06 | 0.07 |
| rs17015996 | 4:90604619 | T / A | 0.20 | 0.18 | 0.17 |
| rs6532182 | 4:90605949 | T / A | 0.15 | 0.14 | 0.13 |
| rs6532183 | 4:90606054 | G / A | 0.08 | 0.07 | 0.07 |
| rs3857047 | 4:90606518 | G / T | 0.13 | 0.12 | 0.11 |
| rs356229 | 4:90606597 | C / T | 0.33 | 0.41 | 0.37 |
| rs10029694 | 4:90607077 | C / G | 0.13 | 0.11 | 0.11 |
| rs356228 | 4:90607126 | C / G | 0.41 | 0.48 | 0.44 |
| rs3857048 | 4:90608959 | T / C | 0.13 | 0.11 | 0.11 |
| rs3910106 | 4:90610135 | T / A | 0.15 | 0.13 | 0.13 |
| rs112180327 | 4:90616430 | T / A | 0.03 | NA | 0.05 |
| rs17286290 | 4:90618264 | G / T | 0.08 | 0.07 | 0.07 |
| rs3906628 | 4:90618526 | A / G | 0.13 | 0.11 | 0.11 |
| rs77918812 | 4:90622129 | A / T | 0.08 | 0.07 | 0.07 |
| rs11734067 | 4:90624853 | A / G | 0.08 | 0.07 | 0.07 |
| rs3857050 | 4:90625419 | C / T | 0.13 | 0.11 | 0.11 |
| rs6812245 | 4:90625714 | A / T | 0.08 | 0.07 | 0.08 |
| rs356183 | 4:90626098 | G / C | 0.40 | 0.48 | 0.43 |
| rs356182 | 4:90626111 | A / G | 0.70 | 1.00 | 0.66 |
| rs356181 | 4:90626139 | G / A | 0.41 | 0.50 | 0.44 |
| rs356180 | 4:90628127 | A / G | 0.25 | 0.34 | 0.30 |
| rs62306277 | 4:90628267 | T / C | 0.06 | 0.06 | 0.05 |
| rs356179 | 4:90628439 | C / G | 0.28 | 0.36 | 0.33 |
| rs356178 | 4:90629465 | G / C | 0.35 | 0.43 | 0.39 |
| rs356177 | 4:90630675 | C / A | 0.24 | 0.33 | 0.29 |
| rs356176 | 4:90630801 | G / C | 0.28 | 0.36 | 0.33 |
| rs356175 | 4:90630814 | C / T | 0.28 | 0.36 | 0.33 |
| rs356174 | 4:90630901 | G / T | 0.28 | 0.36 | 0.33 |
| rs356173 | 4:90631716 | G / A | 0.28 | 0.36 | 0.33 |
| rs356172 | 4:90631986 | C / T | 0.28 | 0.36 | 0.33 |
| rs356171 | 4:90632023 | C / T | 0.28 | 0.36 | 0.33 |
| rs356170 | 4:90632276 | G / C | 0.25 | 0.33 | 0.30 |
| rs356169 | 4:90632768 | G / T | 0.28 | 0.36 | 0.33 |
| rs62306279 | 4:90633693 | C / T | 0.06 | 0.06 | 0.05 |
| rs2572322 | 4:90634546 | G / C | 0.25 | 0.33 | 0.30 |
| rs2572323 | 4:90634552 | A / G | 0.25 | 0.33 | 0.30 |
| rs6825421 | 4:90634806 | T / A | 0.25 | 0.33 | 0.30 |
| rs6848708 | 4:90634828 | G / T | 0.25 | 0.33 | 0.30 |
| rs181489 | 4:90635020 | T / C | 0.25 | 0.33 | 0.30 |
| rs181490 | 4:90635033 | A / G | 0.25 | 0.33 | 0.30 |
| rs356206 | 4:90635049 | T / C | 0.25 | 0.33 | 0.30 |
| rs73831461 | 4:90635338 | C / G | 0.07 | 0.08 | 0.06 |
| rs356207 | 4:90635407 | C / T | 0.25 | 0.33 | 0.30 |
| rs356208 | 4:90635453 | A / T | 0.25 | 0.33 | 0.30 |
| rs62306280 | 4:90635738 | T / C | 0.06 | 0.06 | 0.05 |
| rs356210 | 4:90636193 | T / C | 0.25 | 0.33 | 0.30 |
| rs356211 | 4:90636418 | C / T | 0.42 | 0.50 | 0.45 |
| rs3933594 | 4:90636419 | A / G | 0.12 | 0.11 | 0.10 |
| rs356213 | 4:90636489 | G / T | 0.25 | 0.33 | 0.30 |
| rs356214 | 4:90636541 | C / T | 0.25 | 0.33 | 0.30 |
| rs356215 | 4:90636561 | G / A | 0.36 | 0.43 | 0.39 |
| rs4437213 | 4:90636629 | T / A | 0.25 | 0.33 | 0.30 |
| rs5019538 | 4:90636630 | G / A | 0.25 | 0.33 | 0.30 |
| rs356218 | 4:90637010 | A / G | 0.28 | 0.36 | 0.33 |
| rs9999231 | 4:90637425 | C / T | 0.11 | 0.10 | 0.09 |
| rs356219 | 4:90637601 | G / A | 0.32 | 0.41 | 0.36 |
| rs7681312 | 4:90638527 | G / C | 0.08 | 0.08 | 0.07 |
| rs7681815 | 4:90638614 | C / G | 0.08 | 0.08 | 0.07 |
| rs60838312 | 4:90638896 | G / A | 0.06 | 0.06 | 0.05 |
| rs11931074 | 4:90639515 | T / G | 0.08 | 0.08 | 0.07 |
| rs10516844 | 4:90639847 | C / T | 0.08 | 0.08 | 0.07 |
| rs62306283 | 4:90641048 | A / G | 0.06 | 0.06 | 0.05 |
| rs356220 | 4:90641340 | T / C | 0.33 | 0.41 | 0.36 |
| rs61032876 | 4:90641403 | T / C | 0.08 | 0.08 | 0.07 |
| rs7655792 | 4:90642025 | T / G | 0.08 | 0.08 | 0.07 |
| rs356221 | 4:90642464 | A / T | 0.42 | 0.50 | 0.44 |
| rs62306284 | 4:90642531 | G / A | 0.09 | 0.08 | 0.07 |
| rs6818319 | 4:90642868 | G / A | 0.08 | 0.08 | 0.07 |
| rs10003708 | 4:90643051 | A / C | 0.08 | 0.08 | 0.07 |
| rs356222 | 4:90643123 | C / T | 0.24 | 0.32 | 0.29 |
| rs168552 | 4:90643144 | C / T | 0.24 | 0.33 | 0.29 |
| rs11945223 | 4:90643325 | G / A | 0.08 | 0.08 | 0.07 |
| rs356224 | 4:90643623 | A / G | 0.24 | 0.33 | 0.29 |
| rs3857051 | 4:90643697 | G / A | 0.08 | 0.08 | 0.07 |
| rs356225 | 4:90643757 | C / G | 0.42 | 0.50 | 0.44 |
| rs62306285 | 4:90643807 | C / T | 0.06 | 0.06 | 0.05 |
| rs3857052 | 4:90643857 | A / G | 0.08 | 0.08 | 0.07 |
| rs7436973 | 4:90643921 | C / G | 0.08 | 0.08 | 0.07 |
| rs17016071 | 4:90644281 | G / A | 0.08 | 0.08 | 0.07 |
| rs8180209 | 4:90644454 | G / A | 0.08 | 0.08 | 0.07 |
| rs8180214 | 4:90644508 | A / G | 0.08 | 0.08 | 0.07 |
| rs7675105 | 4:90644960 | T / C | 0.08 | 0.06 | 0.06 |
| rs7675290 | 4:90645003 | T / C | 0.08 | 0.08 | 0.07 |
| rs1045722 | 4:90645671 | A / T | 0.08 | 0.08 | 0.07 |
| rs3857053 | 4:90645674 | T / C | 0.08 | 0.08 | 0.07 |
| rs356166 | 4:90649290 | C / G | 0.24 | 0.33 | 0.29 |
| rs35761327 | 4:90650910 | G / C | 0.06 | 0.06 | 0.05 |
| rs17180453 | 4:90653134 | T / C | 0.05 | 0.08 | 0.07 |
| rs10033209 | 4:90653704 | G / T | 0.08 | 0.08 | 0.07 |
| rs3775422 | 4:90654664 | A / G | 0.08 | 0.08 | 0.07 |
| rs7684318 | 4:90655003 | C / T | 0.07 | 0.08 | 0.06 |
| rs62306303 | 4:90655484 | T / C | 0.06 | 0.06 | 0.05 |
| rs356205 | 4:90657186 | C / T | 0.24 | 0.33 | 0.29 |
| rs3775423 | 4:90657491 | T / C | 0.08 | 0.08 | 0.07 |
| rs62306304 | 4:90659929 | C / G | 0.06 | 0.06 | 0.05 |
| rs6820267 | 4:90660422 | A / G | 0.06 | 0.06 | 0.05 |
| rs3857055 | 4:90660740 | A / G | 0.06 | 0.06 | 0.05 |
| rs142059155 | 4:90661018 | A / G | 0.04 | 0.05 | 0.06 |
| rs62306305 | 4:90661612 | A / G | 0.06 | 0.06 | 0.05 |
| rs62306306 | 4:90662853 | C / G | 0.06 | 0.06 | 0.05 |
| rs356204 | 4:90663542 | T / C | 0.42 | 0.50 | 0.45 |
| rs7661330 | 4:90663670 | G / T | 0.08 | 0.08 | 0.07 |
| rs3822086 | 4:90664794 | T / C | 0.08 | 0.08 | 0.07 |
| rs356203 | 4:90666041 | C / T | 0.33 | 0.41 | 0.36 |
| rs356202 | 4:90666291 | G / A | 0.24 | 0.33 | 0.30 |
| rs3857057 | 4:90668019 | G / A | 0.07 | 0.08 | 0.06 |
| rs356201 | 4:90668419 | C / A | 0.29 | 0.36 | 0.33 |
| rs356200 | 4:90668614 | T / C | 0.42 | 0.50 | 0.45 |
| rs4088093 | 4:90671287 | G / A | 0.08 | 0.08 | 0.07 |
| rs189596 | 4:90671336 | G / A | 0.42 | 0.50 | 0.45 |
| rs4088094 | 4:90671487 | A / G | 0.07 | 0.08 | 0.06 |
| rs28412513 | 4:90671549 | G / A | 0.07 | 0.08 | 0.06 |
| rs2736991 | 4:90671670 | A / T | 0.24 | 0.33 | 0.29 |
| rs3857058 | 4:90672078 | G / A | 0.07 | 0.08 | 0.06 |
| rs190165673 | 4:90672457 | T / A | 0.06 | 0.06 | 0.05 |
| rs356167 | 4:90673770 | A / G | 0.22 | 0.31 | 0.27 |
| rs28393675 | 4:90673841 | G / A | 0.07 | 0.08 | 0.06 |
| rs35236343 | 4:90674230 | C / G | 0.06 | 0.06 | 0.05 |
| rs356168 | 4:90674431 | G / A | 0.42 | 0.50 | 0.45 |
| rs3756054 | 4:90674451 | C / T | 0.08 | 0.08 | 0.07 |
| rs3857059 | 4:90675238 | G / A | 0.08 | 0.08 | 0.07 |
| rs11097231 | 4:90677148 | T / C | 0.24 | 0.33 | 0.30 |
| rs28613708 | 4:90677243 | A / G | 0.07 | 0.08 | 0.06 |
| rs2736990 | 4:90678541 | G / A | 0.42 | 0.50 | 0.45 |
| rs2572324 | 4:90678798 | G / A | 0.24 | 0.33 | 0.29 |
| rs112645122 | 4:90681236 | C / A | 0.04 | 0.05 | 0.06 |
| rs356199 | 4:90682327 | G / A | 0.23 | 0.30 | 0.27 |
| rs6826785 | 4:90682474 | C / T | 0.08 | 0.09 | 0.07 |
| rs356198 | 4:90682504 | T / C | 0.17 | 0.19 | 0.22 |
| rs3910105 | 4:90682571 | G / A | 0.52 | 0.42 | 0.44 |
| rs356197 | 4:90682750 | A / G | 0.17 | 0.19 | 0.22 |
| rs356196 | 4:90682803 | T / A | 0.17 | 0.19 | 0.22 |
| rs356195 | 4:90683168 | T / C | 0.23 | 0.30 | 0.27 |
| rs6834765 | 4:90683990 | C / T | 0.07 | 0.08 | 0.06 |
| rs34806123 | 4:90684122 | G / A | 0.07 | 0.08 | 0.06 |
| rs35495602 | 4:90684123 | G / A | 0.07 | 0.08 | 0.06 |
| rs10516845 | 4:90684278 | G / A | 0.07 | 0.08 | 0.06 |
| rs168550 | 4:90685632 | C / G | 0.16 | 0.17 | 0.21 |
| rs3857061 | 4:90686742 | G / A | 0.08 | 0.08 | 0.07 |
| rs7690873 | 4:90687862 | A / G | 0.07 | 0.08 | 0.06 |
| rs356192 | 4:90687927 | G / A | 0.23 | 0.30 | 0.27 |
| rs356191 | 4:90688120 | A / G | 0.20 | 0.21 | 0.24 |
| rs60031383 | 4:90690071 | A / C | 0.07 | 0.08 | 0.06 |
| rs11944331 | 4:90690329 | T / C | 0.08 | 0.09 | 0.07 |
| rs11931062 | 4:90690704 | C / T | 0.07 | 0.08 | 0.06 |
| rs3910104 | 4:90690748 | T / C | 0.44 | 0.35 | 0.36 |
| rs11935469 | 4:90690832 | G / A | 0.07 | 0.08 | 0.06 |
| rs3899608 | 4:90690931 | G / A | 0.07 | 0.08 | 0.06 |
| rs356189 | 4:90691132 | T / C | 0.23 | 0.30 | 0.27 |
| rs356188 | 4:90691537 | C / T | 0.20 | 0.21 | 0.24 |
| rs356187 | 4:90692468 | T / C | 0.23 | 0.30 | 0.27 |
| rs356164 | 4:90693476 | G / C | 0.12 | 0.14 | 0.16 |
| rs3775433 | 4:90694587 | G / T | 0.07 | 0.08 | 0.06 |
| rs10025915 | 4:90695508 | A / G | 0.07 | 0.08 | 0.06 |
| rs58054215 | 4:90696036 | C / T | 0.07 | 0.08 | 0.06 |
| rs356163 | 4:90696827 | A / C | 0.23 | 0.30 | 0.27 |
| rs356162 | 4:90697157 | C / T | 0.20 | 0.21 | 0.24 |
| rs7698672 | 4:90697466 | C / T | 0.07 | 0.08 | 0.06 |
| rs184810 | 4:90697979 | C / T | 0.20 | 0.21 | 0.24 |
| rs4031753 | 4:90700161 | G / C | 0.08 | 0.08 | 0.07 |
| rs28781297 | 4:90701442 | G / A | 0.08 | 0.08 | 0.06 |
| rs7655611 | 4:90702146 | A / G | 0.08 | 0.08 | 0.06 |
| rs3775434 | 4:90702781 | G / A | 0.13 | 0.13 | 0.11 |
| rs356184 | 4:90703233 | A / G | 0.23 | 0.30 | 0.27 |
| rs10018362 | 4:90703753 | C / T | 0.11 | 0.12 | 0.10 |
| rs3822089 | 4:90704011 | A / G | 0.13 | 0.13 | 0.11 |
| rs356185 | 4:90704617 | A / G | 0.23 | 0.30 | 0.27 |
| rs3822090 | 4:90704876 | T / C | 0.13 | 0.13 | 0.11 |
| rs356186 | 4:90705364 | A / G | 0.18 | 0.19 | 0.23 |
| rs2737033 | 4:90707947 | C / T | 0.23 | 0.30 | 0.27 |
| rs2737032 | 4:90709169 | G / A | 0.12 | 0.14 | 0.16 |
| rs3775439 | 4:90709741 | A / G | 0.13 | 0.13 | 0.11 |
| rs2737030 | 4:90710099 | C / T | 0.20 | 0.21 | 0.24 |
| rs2737029 | 4:90711770 | C / T | 0.36 | 0.44 | 0.38 |
| rs2572320 | 4:90712205 | C / A | 0.18 | 0.19 | 0.23 |
| rs10014396 | 4:90712629 | C / T | 0.11 | 0.12 | 0.10 |
| rs12502363 | 4:90713064 | A / G | 0.45 | 0.36 | 0.37 |
| rs17016168 | 4:90713259 | A / G | 0.06 | 0.07 | 0.06 |
| rs11097234 | 4:90713330 | G / C | 0.45 | 0.36 | 0.37 |
| rs7665887 | 4:90713674 | G / A | 0.06 | 0.07 | 0.06 |
| rs2583962 | 4:90713747 | T / C | 0.20 | 0.21 | 0.24 |
| rs6829514 | 4:90714635 | G / A | 0.06 | 0.07 | 0.06 |
| rs3775443 | 4:90715674 | G / T | 0.06 | 0.07 | 0.06 |
| rs3906824 | 4:90716139 | G / C | 0.06 | 0.07 | 0.06 |
| rs3889917 | 4:90716852 | C / T | 0.06 | 0.07 | 0.06 |
| rs3889916 | 4:90716952 | T / C | 0.06 | 0.07 | 0.06 |
| rs2737028 | 4:90717016 | A / G | 0.20 | 0.21 | 0.24 |
| rs34634504 | 4:90717245 | A / G | 0.06 | 0.07 | 0.06 |
| rs2572318 | 4:90718719 | C / T | 0.20 | 0.21 | 0.24 |
| rs62306323 | 4:90718995 | T / C | 0.14 | 0.10 | 0.12 |
| rs56270968 | 4:90719020 | T / G | 0.06 | 0.07 | 0.06 |
| rs2737025 | 4:90719192 | A / G | 0.20 | 0.21 | 0.24 |
| rs6532190 | 4:90719196 | T / A | 0.07 | 0.07 | 0.05 |
| rs17016183 | 4:90719460 | C / T | 0.06 | 0.07 | 0.06 |
| rs2583958 | 4:90720365 | C / T | 0.22 | 0.30 | 0.27 |
| rs3756055 | 4:90720957 | G / A | 0.06 | 0.07 | 0.06 |
| rs2737024 | 4:90721560 | G / A | 0.22 | 0.30 | 0.27 |
| rs2583959 | 4:90721637 | G / C | 0.22 | 0.30 | 0.27 |
| rs17016188 | 4:90721880 | C / T | 0.06 | 0.07 | 0.05 |
| rs7684892 | 4:90721997 | A / G | 0.06 | 0.07 | 0.06 |
| rs7684637 | 4:90722145 | C / A | 0.06 | 0.07 | 0.06 |
| rs7689942 | 4:90722400 | T / C | 0.06 | 0.07 | 0.06 |
| rs2619373 | 4:90722433 | A / G | 0.22 | 0.30 | 0.27 |
| rs17016190 | 4:90722832 | G / A | 0.06 | 0.07 | 0.06 |
| rs6848726 | 4:90722871 | T / C | 0.45 | 0.36 | 0.38 |
| rs2583960 | 4:90724869 | G / A | 0.26 | 0.27 | 0.31 |
| rs17016193 | 4:90726022 | G / T | 0.06 | 0.07 | 0.05 |
| rs34164595 | 4:90726089 | A / C | 0.06 | 0.07 | 0.05 |
| rs2583963 | 4:90727088 | C / T | 0.20 | 0.21 | 0.24 |
| rs2583964 | 4:90727218 | A / G | 0.20 | 0.21 | 0.24 |
| rs3775446 | 4:90727923 | T / G | 0.06 | 0.07 | 0.06 |
| rs3775447 | 4:90728423 | A / G | 0.06 | 0.07 | 0.06 |
| rs2197120 | 4:90729602 | A / G | 0.20 | 0.21 | 0.24 |
| rs2619368 | 4:90729747 | T / G | 0.20 | 0.21 | 0.24 |
| rs3796664 | 4:90730687 | A / C | 0.45 | 0.36 | 0.38 |
| rs59398886 | 4:90731685 | A / G | 0.06 | 0.07 | 0.06 |
| rs61027512 | 4:90731736 | G / C | 0.06 | 0.07 | 0.06 |
| rs34410265 | 4:90733379 | T / G | 0.06 | 0.07 | 0.06 |
| rs3775448 | 4:90734105 | G / C | 0.06 | 0.07 | 0.06 |
| rs3756056 | 4:90734333 | C / T | 0.06 | 0.07 | 0.06 |
| rs3756057 | 4:90734426 | C / T | 0.06 | 0.07 | 0.06 |
| rs894278 | 4:90734535 | G / T | 0.06 | 0.07 | 0.05 |
| rs2119787 | 4:90734728 | G / A | 0.45 | 0.36 | 0.38 |
| rs748849 | 4:90734961 | G / A | 0.20 | 0.21 | 0.24 |
| rs1837890 | 4:90736006 | A / C | 0.26 | 0.27 | 0.31 |
| rs1442143 | 4:90736024 | A / T | 0.26 | 0.27 | 0.31 |
| rs1442144 | 4:90736113 | T / C | 0.20 | 0.21 | 0.24 |
| rs1837891 | 4:90736131 | C / G | 0.20 | 0.21 | 0.24 |
| rs3822095 | 4:90736517 | C / T | 0.45 | 0.36 | 0.38 |
| rs2619370 | 4:90736585 | T / C | 0.22 | 0.30 | 0.27 |
| rs2619371 | 4:90736678 | G / A | 0.22 | 0.30 | 0.27 |
| rs2619372 | 4:90736727 | A / G | 0.26 | 0.27 | 0.31 |
| rs1442145 | 4:90736923 | G / A | 0.26 | 0.27 | 0.31 |
| rs974711 | 4:90737327 | A / G | 0.52 | 0.42 | 0.42 |
| rs17016235 | 4:90737879 | C / G | 0.06 | 0.07 | 0.05 |
| rs77164054 | 4:90738148 | A / T | 0.03 | 0.04 | 0.05 |
| rs10155475 | 4:90738158 | C / G | 0.45 | 0.36 | 0.38 |
| rs3775458 | 4:90739248 | C / T | 0.06 | 0.07 | 0.06 |
| rs972880 | 4:90739255 | A / G | 0.26 | 0.27 | 0.31 |
| rs2737023 | 4:90739505 | C / T | 0.22 | 0.30 | 0.27 |
| rs1812923 | 4:90739539 | A / C | 0.52 | 0.42 | 0.42 |
| rs2737022 | 4:90739662 | C / A | 0.22 | 0.30 | 0.27 |
| rs2737021 | 4:90739992 | T / A | 0.20 | 0.21 | 0.24 |
| rs2583965 | 4:90740103 | T / G | 0.22 | 0.30 | 0.27 |
| rs2737020 | 4:90740878 | G / A | 0.26 | 0.27 | 0.31 |
| rs2583966 | 4:90741519 | A / G | 0.22 | 0.30 | 0.27 |
| rs2619341 | 4:90741773 | A / G | 0.20 | 0.21 | 0.24 |
| rs3775461 | 4:90741803 | G / A | 0.06 | 0.07 | 0.06 |
| rs17016251 | 4:90742639 | C / G | 0.51 | 0.42 | 0.44 |
| rs35897584 | 4:90742692 | G / A | 0.06 | 0.07 | 0.05 |
| rs2298728 | 4:90742815 | A / G | 0.06 | 0.07 | 0.05 |
| rs2298727 | 4:90742861 | G / T | 0.06 | 0.07 | 0.05 |
| rs10005233 | 4:90743331 | C / T | 0.42 | 0.51 | 0.51 |
| rs7356228 | 4:90743855 | G / C | 0.45 | 0.36 | 0.38 |
| rs2583967 | 4:90744100 | T / C | 0.22 | 0.30 | 0.27 |
| rs2619342 | 4:90744170 | T / A | 0.22 | 0.30 | 0.27 |
| rs34238791 | 4:90744216 | A / G | 0.07 | 0.07 | 0.05 |
| rs2619343 | 4:90744221 | T / C | 0.22 | 0.30 | 0.27 |
| rs2619344 | 4:90744248 | T / C | 0.23 | 0.31 | 0.28 |
| rs2619345 | 4:90744270 | G / A | 0.22 | 0.30 | 0.27 |
| rs17016255 | 4:90744328 | A / G | 0.06 | 0.07 | 0.06 |
| rs2737019 | 4:90744903 | T / G | 0.22 | 0.30 | 0.27 |
| rs6830166 | 4:90744993 | T / C | 0.13 | 0.13 | 0.11 |
| rs2737018 | 4:90745091 | C / T | 0.22 | 0.30 | 0.26 |
| rs4293762 | 4:90745096 | G / A | 0.42 | 1.00 | 0.51 |
| rs6830383 | 4:90745113 | C / A | 0.07 | 0.07 | 0.05 |
| rs2737017 | 4:90745238 | T / C | 0.22 | 0.30 | 0.26 |
| rs2737016 | 4:90745242 | C / T | 0.22 | 0.30 | 0.26 |
| rs2737015 | 4:90745283 | C / T | 0.22 | 0.30 | 0.26 |
| rs3113355 | 4:90745345 | A / T | 0.20 | 0.21 | 0.25 |
| rs2583968 | 4:90745395 | C / G | 0.20 | 0.21 | 0.25 |
| rs2737014 | 4:90745456 | C / A | 0.22 | 0.30 | 0.26 |
| rs2737013 | 4:90745503 | T / C | 0.22 | 0.30 | 0.26 |
| rs2737012 | 4:90745707 | A / G | 0.22 | 0.30 | 0.26 |
| rs2619347 | 4:90745770 | G / A | 0.20 | 0.21 | 0.25 |
| rs6532191 | 4:90745930 | C / T | 0.42 | 1.00 | 0.51 |
| rs2583969 | 4:90746133 | C / T | 0.22 | 0.30 | 0.26 |
| rs2737011 | 4:90746219 | T / C | 0.19 | 0.20 | 0.23 |
| rs2619348 | 4:90746321 | G / C | 0.22 | 0.30 | 0.26 |
| rs7356297 | 4:90746329 | C / T | 0.45 | 0.35 | 0.37 |
| rs2737010 | 4:90746466 | G / A | 0.20 | 0.21 | 0.25 |
| rs2583970 | 4:90746610 | A / G | 0.22 | 0.30 | 0.26 |
| rs2619349 | 4:90746646 | G / T | 0.20 | 0.21 | 0.25 |
| rs11097238 | 4:90746709 | T / C | 0.45 | 0.35 | 0.37 |
| rs2737009 | 4:90746836 | G / A | 0.22 | 0.30 | 0.26 |
| rs2619350 | 4:90747004 | G / A | 0.22 | 0.30 | 0.26 |
| rs2737008 | 4:90747183 | A / G | 0.22 | 0.30 | 0.26 |
| rs2583971 | 4:90747207 | T / C | 0.22 | 0.30 | 0.26 |
| rs2583972 | 4:90747411 | T / G | 0.20 | 0.21 | 0.25 |
| rs2583973 | 4:90747499 | T / C | 0.22 | 0.30 | 0.26 |
| rs2619351 | 4:90747704 | G / A | 0.22 | 0.30 | 0.26 |
| rs2619352 | 4:90747709 | T / C | 0.22 | 0.30 | 0.26 |
| rs1811442 | 4:90747751 | C / T | 0.20 | 0.21 | 0.25 |
| rs1811443 | 4:90747883 | G / C | 0.20 | 0.21 | 0.25 |
| rs2619353 | 4:90747975 | C / G | 0.22 | 0.30 | 0.26 |
| rs920624 | 4:90748195 | T / A | 0.42 | 1.00 | 0.51 |
| rs2619354 | 4:90748284 | T / C | 0.22 | 0.30 | 0.26 |
| rs2583974 | 4:90748297 | C / T | 0.22 | 0.30 | 0.26 |
| rs2619355 | 4:90748374 | A / C | 0.20 | 0.21 | 0.25 |
| rs2583975 | 4:90748488 | T / C | 0.19 | 0.20 | 0.23 |
| rs1442146 | 4:90748646 | G / C | 0.22 | 0.30 | 0.26 |
| rs3796665 | 4:90748710 | A / G | 0.45 | 0.35 | 0.38 |
| rs1442147 | 4:90748738 | T / C | 0.22 | 0.30 | 0.26 |
| rs1442148 | 4:90748846 | T / A | 0.20 | 0.21 | 0.25 |
| rs1442149 | 4:90749132 | T / A | 0.22 | 0.30 | 0.26 |
| rs2737006 | 4:90749686 | G / A | 0.22 | 0.30 | 0.26 |
| rs34699553 | 4:90749966 | T / C | 0.06 | 0.07 | 0.06 |
| rs6828271 | 4:90750025 | C / T | 0.42 | 1.00 | 0.51 |
| rs33965306 | 4:90750043 | G / A | 0.22 | 0.30 | 0.26 |
| rs13142587 | 4:90750169 | T / G | 0.22 | 0.30 | 0.26 |
| rs2583976 | 4:90750173 | C / T | 0.22 | 0.30 | 0.26 |
| rs2583977 | 4:90750225 | T / C | 0.22 | 0.30 | 0.26 |
| rs2583978 | 4:90750326 | A / C | 0.20 | 0.21 | 0.25 |
| rs2583979 | 4:90750588 | A / T | 0.20 | 0.21 | 0.25 |
| rs2737005 | 4:90750590 | G / A | 0.22 | 0.30 | 0.26 |
| rs2737004 | 4:90750722 | G / A | 0.22 | 0.30 | 0.26 |
| rs2619356 | 4:90750844 | C / T | 0.20 | 0.21 | 0.25 |
| rs2583980 | 4:90751109 | G / T | 0.22 | 0.30 | 0.26 |
| rs986609 | 4:90751415 | G / C | 0.22 | 0.30 | 0.26 |
| rs986610 | 4:90751655 | C / T | 0.22 | 0.30 | 0.26 |
| rs2619357 | 4:90752205 | T / C | 0.12 | 0.13 | 0.16 |
| rs2583981 | 4:90752582 | T / A | 0.22 | 0.30 | 0.26 |
| rs60642683 | 4:90752646 | A / C | 0.06 | 0.07 | 0.06 |
| rs2619358 | 4:90752670 | C / A | 0.22 | 0.30 | 0.26 |
| rs2737003 | 4:90753068 | G / A | 0.22 | 0.30 | 0.26 |
| rs2737002 | 4:90753180 | A / G | 0.19 | 0.20 | 0.24 |
| rs2583982 | 4:90753244 | C / T | 0.22 | 0.30 | 0.26 |
| rs2737001 | 4:90753477 | A / G | 0.22 | 0.30 | 0.26 |
| rs2583983 | 4:90753960 | T / C | 0.22 | 0.30 | 0.26 |
| rs2737000 | 4:90754022 | A / C | 0.22 | 0.30 | 0.26 |
| rs2736999 | 4:90754111 | G / A | 0.22 | 0.30 | 0.26 |
| rs2583984 | 4:90754153 | C / T | 0.22 | 0.30 | 0.26 |
| rs1471483 | 4:90754292 | T / C | 0.42 | 1.00 | 0.50 |
| rs1471484 | 4:90754313 | C / T | 0.19 | 0.20 | 0.24 |
| rs2736998 | 4:90754336 | A / C | 0.22 | 0.30 | 0.26 |
| rs2736997 | 4:90754557 | A / G | 0.22 | 0.30 | 0.26 |
| rs2736996 | 4:90754691 | G / A | 0.22 | 0.30 | 0.26 |
| rs990085 | 4:90754771 | A / G | 0.19 | 0.20 | 0.24 |
| rs2619359 | 4:90754828 | C / A | 0.22 | 0.30 | 0.26 |
| rs990086 | 4:90755090 | T / A | 0.22 | 0.30 | 0.26 |
| rs990087 | 4:90755177 | C / A | 0.22 | 0.30 | 0.26 |
| rs990088 | 4:90755225 | C / T | 0.22 | 0.30 | 0.26 |
| rs1372516 | 4:90755730 | A / G | 0.22 | 0.30 | 0.26 |
| rs1372517 | 4:90755909 | G / A | 0.42 | 1.00 | 0.50 |
| rs2583985 | 4:90755939 | G / A | 0.22 | 0.30 | 0.26 |
| rs2035268 | 4:90756077 | G / T | 0.06 | 0.07 | 0.06 |
| rs2619360 | 4:90756191 | T / C | 0.22 | 0.30 | 0.26 |
| rs2028535 | 4:90756421 | C / G | 0.22 | 0.30 | 0.26 |
| rs7681440 | 4:90756550 | C / G | 0.42 | 1.00 | 0.50 |
| rs3756059 | 4:90757272 | A / G | 0.57 | 0.49 | 0.48 |
| rs1372518 | 4:90757294 | A / C | 0.21 | 0.21 | 0.25 |
| rs1372519 | 4:90757309 | A / G | 0.21 | 0.21 | 0.25 |
| rs3756063 | 4:90757394 | C / G | 0.57 | 0.49 | 0.48 |
| rs1372520 | 4:90757505 | T / C | 0.21 | 0.21 | 0.25 |
| rs2619361 | 4:90757735 | A / C | 0.22 | 0.30 | 0.26 |
| rs2245801 | 4:90757840 | T / C | 0.20 | 0.20 | 0.24 |
| rs2619362 | 4:90757845 | T / C | 0.22 | 0.30 | 0.26 |
| rs2301135 | 4:90758389 | C / G | 0.57 | 0.49 | 0.48 |
| rs2301134 | 4:90758945 | G / A | 0.57 | 0.49 | 0.48 |
| rs2619363 | 4:90759047 | T / G | 0.22 | 0.30 | 0.26 |
| rs3806789 | 4:90759556 | T / C | 0.57 | 0.49 | 0.48 |
| rs2619364 | 4:90759887 | G / A | 0.22 | 0.29 | 0.26 |
| rs2583987 | 4:90760221 | C / A | 0.22 | 0.29 | 0.26 |
| rs2583988 | 4:90760828 | T / C | 0.22 | 0.29 | 0.26 |
| rs894280 | 4:90760883 | T / C | 0.57 | 0.49 | 0.48 |
| rs17016274 | 4:90761357 | A / T | 0.06 | 0.07 | 0.05 |
| rs1023777 | 4:90761412 | C / T | 0.44 | 0.36 | 0.37 |
| rs983361 | 4:90761944 | T / G | 0.21 | 0.22 | 0.26 |
| rs2736995 | 4:90762197 | C / A | 0.22 | 0.29 | 0.26 |
| rs10009760 | 4:90762622 | T / A | 0.44 | 0.36 | 0.37 |
| rs2619366 | 4:90763260 | G / A | 0.22 | 0.29 | 0.26 |
| rs7680557 | 4:90763360 | A / C | 0.57 | 0.49 | 0.48 |
| rs7680761 | 4:90763510 | T / A | 0.57 | 0.49 | 0.48 |
| rs7681154 | 4:90763703 | C / A | 0.57 | 0.49 | 0.48 |
| rs6532192 | 4:90764131 | A / G | 0.57 | 0.49 | 0.48 |
| rs6532193 | 4:90764188 | A / G | 0.57 | 0.49 | 0.48 |
| rs6822088 | 4:90764310 | C / T | 0.12 | 0.13 | 0.11 |
| rs7687945 | 4:90764699 | T / C | 0.57 | 0.49 | 0.48 |
| rs1822217 | 4:90765052 | T / C | 0.56 | 0.48 | 0.47 |
| rs2870028 | 4:90765213 | T / C | 0.44 | 0.35 | 0.37 |
| rs1372521 | 4:90765223 | A / G | 0.56 | 0.48 | 0.47 |
| rs1372522 | 4:90765280 | G / A | 0.56 | 0.48 | 0.47 |
| rs6816736 | 4:90765746 | A / G | 0.56 | 0.48 | 0.47 |
| rs6816633 | 4:90765944 | T / A | 0.06 | 0.07 | 0.05 |
| rs6817001 | 4:90766019 | T / C | 0.56 | 0.48 | 0.47 |
| rs6817026 | 4:90766069 | T / C | 0.56 | 0.48 | 0.47 |
| rs6843084 | 4:90766230 | C / T | 0.56 | 0.48 | 0.47 |
| rs10028246 | 4:90766704 | A / G | 0.56 | 0.48 | 0.47 |
| rs58864428 | 4:90766986 | C / T | 0.56 | 0.48 | 0.47 |
| rs28507349 | 4:90767063 | A / G | 0.56 | 0.48 | 0.47 |
| rs2619367 | 4:90767095 | C / A | 0.22 | 0.30 | 0.26 |
| rs28415623 | 4:90767110 | T / C | 0.56 | 0.48 | 0.47 |
| rs28734152 | 4:90767359 | T / C | 0.56 | 0.48 | 0.47 |
| rs10030935 | 4:90767433 | T / C | 0.44 | 0.35 | 0.37 |
| rs115630491 | 4:90767504 | C / T | 0.06 | 0.08 | 0.07 |
| rs7672979 | 4:90767873 | G / A | 0.56 | 0.47 | 0.47 |
| rs35424815 | 4:90767877 | T / C | 0.56 | 0.47 | 0.47 |
| rs7697570 | 4:90767880 | G / T | 0.44 | 0.35 | 0.37 |
| rs7678651 | 4:90768371 | A / C | 0.44 | 0.35 | 0.37 |
| rs765517 | 4:90768889 | A / T | 0.44 | 0.35 | 0.37 |
| rs2583989 | 4:90769422 | G / A | 0.22 | 0.30 | 0.26 |
| rs10004413 | 4:90769479 | C / T | 0.56 | 0.48 | 0.47 |
| rs28403500 | 4:90769729 | G / T | 0.56 | 0.48 | 0.47 |
| rs75278754 | 4:90769859 | C / G | 0.05 | 0.06 | 0.05 |
| rs28777703 | 4:90769924 | T / C | 0.56 | 0.48 | 0.47 |
| rs12331318 | 4:90770255 | C / T | 0.56 | 0.48 | 0.47 |
| rs17016279 | 4:90770408 | G / A | 0.05 | 0.06 | 0.05 |
| rs7671549 | 4:90771147 | A / T | 0.56 | 0.48 | 0.47 |
| rs7698219 | 4:90771351 | A / C | 0.56 | 0.48 | 0.47 |
| rs7676702 | 4:90771397 | C / T | 0.05 | 0.06 | 0.05 |
| rs2583990 | 4:90771499 | A / G | 0.22 | 0.23 | 0.27 |
| rs62304971 | 4:90771626 | T / C | 0.05 | 0.06 | 0.05 |
| rs62304972 | 4:90771987 | G / T | 0.05 | 0.06 | 0.05 |
| rs62304973 | 4:90771992 | T / G | 0.05 | 0.06 | 0.05 |
| rs2870029 | 4:90772057 | T / G | 0.56 | 0.48 | 0.47 |
| rs6815691 | 4:90772489 | A / G | 0.44 | 0.35 | 0.37 |
| rs6841476 | 4:90772697 | C / T | 0.44 | 0.35 | 0.37 |
| rs61579967 | 4:90772725 | G / A | 0.06 | 0.07 | 0.05 |
| rs2736993 | 4:90772851 | C / A | 0.22 | 0.30 | 0.26 |
| rs6816469 | 4:90772885 | C / G | 0.56 | 0.47 | 0.47 |
| rs75017256 | 4:90772929 | T / G | 0.05 | 0.06 | 0.05 |
| rs12643708 | 4:90773770 | A / C | 0.44 | 0.35 | 0.37 |
| rs12649704 | 4:90773800 | T / A | 0.44 | 0.35 | 0.37 |
| rs62304974 | 4:90774131 | G / A | 0.05 | 0.06 | 0.05 |
| rs28750169 | 4:90774303 | G / A | 0.56 | 0.48 | 0.47 |
| rs59480711 | 4:90774461 | T / A | 0.05 | 0.06 | 0.05 |
| rs75226172 | 4:90774492 | A / G | 0.06 | 0.07 | 0.05 |
| rs28567580 | 4:90774615 | T / A | 0.56 | 0.48 | 0.47 |
| rs1551980 | 4:90774724 | A / G | 0.22 | 0.30 | 0.26 |
| rs17192189 | 4:90774928 | T / G | 0.56 | 0.48 | 0.47 |
| rs10516848 | 4:90775212 | G / A | 0.44 | 0.35 | 0.37 |
| rs1442151 | 4:90775491 | A / T | 0.56 | 0.48 | 0.47 |
| rs1442152 | 4:90775649 | C / A | 0.22 | 0.23 | 0.27 |
| rs1442153 | 4:90775719 | G / T | 0.56 | 0.48 | 0.47 |
| rs1442154 | 4:90775761 | G / C | 0.56 | 0.48 | 0.47 |
| rs1372523 | 4:90775923 | A / G | 0.56 | 0.48 | 0.47 |
| rs1372524 | 4:90776045 | G / A | 0.56 | 0.48 | 0.47 |
| rs1372525 | 4:90776165 | A / G | 0.56 | 0.48 | 0.47 |
| rs2736989 | 4:90776257 | T / C | 0.22 | 0.23 | 0.27 |
| rs2736988 | 4:90776345 | T / C | 0.22 | 0.30 | 0.26 |
| rs2737035 | 4:90777213 | T / C | 0.22 | 0.30 | 0.26 |
| rs2737034 | 4:90777464 | G / C | 0.22 | 0.30 | 0.26 |
| rs2619339 | 4:90777468 | C / G | 0.22 | 0.30 | 0.26 |
| rs60193684 | 4:90777774 | T / C | 0.56 | 0.48 | 0.47 |
| rs55732478 | 4:90777837 | C / G | 0.56 | 0.48 | 0.47 |
| rs11725564 | 4:90778040 | A / G | 0.44 | 0.36 | 0.37 |
| rs11733704 | 4:90778044 | G / A | 0.56 | 0.48 | 0.47 |
| rs9884400 | 4:90778833 | A / G | 0.44 | 0.35 | 0.37 |
| rs2619340 | 4:90779518 | C / G | 0.22 | 0.30 | 0.26 |
| rs74402838 | 4:90779710 | A / G | 0.06 | 0.07 | 0.05 |
| rs2583961 | 4:90779758 | A / G | 0.22 | 0.23 | 0.27 |
| rs2737026 | 4:90779823 | A / G | 0.22 | 0.23 | 0.27 |
| rs62315169 | 4:90780148 | G / T | 0.05 | 0.06 | 0.05 |
| rs6532194 | 4:90780902 | T / C | 0.09 | 0.10 | 0.09 |
| rs10025824 | 4:90781538 | C / T | 0.44 | 0.35 | 0.37 |
| rs6841352 | 4:90781655 | T / G | 0.52 | 0.44 | 0.44 |
| rs62315170 | 4:90783392 | C / T | 0.06 | 0.06 | 0.06 |
| rs1871399 | 4:90783545 | G / C | 0.44 | 0.35 | 0.37 |
| rs2736994 | 4:90784528 | A / G | 0.20 | 0.20 | 0.24 |
| rs7656954 | 4:90784670 | G / A | 0.42 | 1.00 | 0.50 |
| rs6831847 | 4:90785341 | G / A | 0.42 | 1.00 | 0.50 |
| rs17016299 | 4:90786665 | A / C | 0.08 | 0.09 | 0.07 |
| rs28796128 | 4:90787905 | T / C | 0.50 | 0.41 | 0.42 |
| rs72657784 | 4:90788943 | A / G | 0.05 | 0.08 | 0.06 |
| rs10516849 | 4:90789539 | G / A | 0.08 | 0.09 | 0.07 |
| rs1372513 | 4:90790237 | G / A | 0.42 | 1.00 | 0.50 |
| rs17016305 | 4:90791381 | G / A | 0.50 | 0.41 | 0.42 |
| rs7668531 | 4:90791819 | G / T | 0.42 | 1.00 | 0.50 |
| rs7693616 | 4:90792230 | G / T | 0.50 | 0.41 | 0.42 |
| rs59559246 | 4:90792765 | G / A | 0.08 | 0.09 | 0.07 |
| rs73831827 | 4:90794942 | A / G | 0.08 | 0.09 | 0.07 |
| rs11727074 | 4:90795281 | C / T | 0.08 | 0.09 | 0.07 |
| rs11721779 | 4:90795356 | C / T | 0.08 | 0.09 | 0.07 |
| rs11097239 | 4:90795804 | A / C | 0.41 | 0.35 | 0.35 |
| rs6848483 | 4:90796613 | A / T | 0.08 | 0.09 | 0.07 |
| rs17016316 | 4:90797235 | G / A | 0.05 | 0.05 | 0.05 |
| rs6532197 | 4:90797301 | G / A | 0.08 | 0.09 | 0.07 |
| rs6532198 | 4:90798870 | C / T | 0.42 | 1.00 | 0.50 |
| rs59559279 | 4:90799072 | T / C | 0.08 | 0.09 | 0.07 |
| rs79438231 | 4:90800258 | T / A | 0.08 | 0.09 | 0.07 |
| rs1899389 | 4:90802998 | G / A | 0.42 | 1.00 | 0.50 |
| rs6848022 | 4:90803635 | C / T | 0.08 | 0.09 | 0.07 |
| rs11941682 | 4:90804316 | G / T | 0.50 | 0.41 | 0.42 |
| rs6839608 | 4:90805493 | T / C | 0.42 | 1.00 | 0.50 |
| rs6814221 | 4:90805501 | T / C | 0.50 | 0.41 | 0.42 |
| rs6814481 | 4:90805660 | T / C | 0.50 | 0.41 | 0.43 |
| rs2165614 | 4:90805724 | C / A | 0.22 | 0.30 | 0.26 |
| rs73831836 | 4:90806717 | G / T | 0.08 | 0.09 | 0.07 |
| rs6848181 | 4:90807252 | T / C | 0.42 | 1.00 | 0.50 |
| rs6828095 | 4:90807512 | A / G | 0.08 | 0.09 | 0.07 |
| rs67876224 | 4:90808918 | C / T | 0.22 | 0.30 | 0.26 |
| rs77823616 | 4:90811007 | A / C | 0.06 | 0.08 | 0.06 |
| rs1372511 | 4:90811049 | G / C | 0.08 | 0.09 | 0.07 |
| rs7356205 | 4:90813521 | C / A | 0.50 | 0.41 | 0.42 |
| rs1442141 | 4:90815688 | C / A | 0.06 | 0.05 | 0.06 |
| rs1442140 | 4:90816023 | A / G | 0.42 | 1.00 | 0.50 |
| rs1442138 | 4:90816294 | G / A | 0.06 | 0.08 | 0.06 |
| rs2289515 | 4:90816790 | A / T | 0.50 | 0.41 | 0.42 |
| rs1372510 | 4:90817259 | A / G | 0.50 | 0.41 | 0.42 |
| rs1372509 | 4:90817311 | G / A | 0.50 | 0.41 | 0.42 |
| rs6812192 | 4:90817848 | G / A | 0.56 | 0.48 | 0.48 |
| rs59718120 | 4:90817981 | C / T | 0.06 | 0.05 | 0.06 |
| rs6812593 | 4:90817991 | T / C | 0.50 | 0.41 | 0.42 |
| rs10001067 | 4:90818216 | G / C | 0.56 | 0.48 | 0.48 |
| rs1904350 | 4:90818514 | C / T | 0.50 | 0.41 | 0.42 |
| rs1442137 | 4:90818857 | C / T | 0.06 | 0.05 | 0.06 |
| rs1442136 | 4:90819026 | C / T | 0.50 | 0.41 | 0.42 |
| rs1442135 | 4:90819093 | T / C | 0.42 | 1.00 | 0.50 |
| rs1442134 | 4:90819154 | C / T | 0.56 | 0.48 | 0.48 |
| rs1442133 | 4:90819195 | T / C | 0.50 | 0.41 | 0.42 |
| rs4522832 | 4:90819391 | C / A | 0.50 | 0.41 | 0.42 |
| rs6824979 | 4:90819407 | G / C | 0.50 | 0.41 | 0.42 |
| rs11941110 | 4:90819569 | C / T | 0.50 | 0.41 | 0.42 |
| rs1372508 | 4:90819786 | G / C | 0.50 | 0.41 | 0.42 |
| rs763443 | 4:90819961 | T / C | 0.42 | 1.00 | 0.50 |
| rs72657796 | 4:90820808 | T / C | 0.15 | 0.20 | 0.18 |
| rs72657797 | 4:90820809 | T / C | 0.15 | 0.20 | 0.18 |
| rs1442131 | 4:90821381 | G / A | 0.09 | 0.09 | 0.08 |
| rs72657800 | 4:90822051 | C / T | 0.09 | 0.09 | 0.08 |
| rs56146644 | 4:90824430 | T / C | 0.09 | 0.09 | 0.08 |
| rs72659403 | 4:90825002 | G / A | 0.09 | 0.09 | 0.08 |
| rs3775465 | 4:90827466 | C / A | 0.08 | 0.08 | 0.07 |
| rs10433955 | 4:90830783 | G / A | 0.09 | 0.09 | 0.08 |
| rs7691487 | 4:90832001 | T / C | 0.40 | 0.47 | 0.47 |
| rs7666981 | 4:90832044 | G / C | 0.09 | 0.09 | 0.07 |
| rs3775467 | 4:90832608 | A / C | 0.08 | 0.07 | 0.06 |
| rs9307077 | 4:90833372 | A / G | 0.40 | 0.47 | 0.47 |
| rs10011790 | 4:90833421 | G / A | 0.42 | 0.35 | 0.35 |
| rs1372507 | 4:90835701 | T / C | 0.40 | 0.47 | 0.47 |
| rs2119785 | 4:90836519 | C / A | 0.52 | 0.44 | 0.46 |
| rs10516850 | 4:90837127 | G / A | 0.09 | 0.09 | 0.07 |
| rs3775473 | 4:90837641 | T / G | 0.09 | 0.09 | 0.07 |
| rs10516851 | 4:90837944 | T / C | 0.09 | 0.09 | 0.07 |
| rs12644375 | 4:90837980 | G / A | 0.09 | 0.09 | 0.07 |
| rs3775474 | 4:90838158 | A / G | 0.09 | 0.09 | 0.07 |
| rs3775475 | 4:90838643 | T / C | 0.51 | 0.44 | 0.46 |
| rs3775477 | 4:90839022 | C / T | 0.09 | 0.09 | 0.07 |
| rs73831856 | 4:90839249 | T / C | 0.09 | 0.09 | 0.07 |
| rs12641781 | 4:90841787 | A / G | 0.09 | 0.09 | 0.07 |
| rs3775478 | 4:90842840 | G / A | 0.09 | 0.09 | 0.07 |
| rs3775479 | 4:90842971 | C / A | 0.51 | 0.45 | 0.46 |
| rs2011543 | 4:90843509 | A / G | 0.40 | 0.47 | 0.47 |
| rs10516852 | 4:90843650 | A / T | 0.09 | 0.09 | 0.07 |
| rs12643823 | 4:90845731 | C / T | 0.09 | 0.09 | 0.07 |
| rs74739978 | 4:90847454 | T / C | 0.09 | 0.08 | 0.07 |
| rs6812172 | 4:90847859 | T / C | 0.06 | 0.06 | 0.06 |
| rs72659413 | 4:90848113 | A / C | 0.08 | 0.07 | 0.06 |
| rs115690633 | 4:90848275 | G / T | 0.05 | 0.07 | 0.05 |
| rs58930449 | 4:90848643 | G / A | 0.08 | 0.07 | 0.06 |
| rs1442129 | 4:90849446 | A / G | 0.41 | 0.48 | 0.48 |
| rs61611959 | 4:90851676 | T / C | 0.09 | 0.09 | 0.08 |
| rs1038140 | 4:90855932 | C / T | 0.44 | 0.37 | 0.38 |
| rs67212195 | 4:90859711 | C / G | 0.08 | 0.07 | 0.06 |
| rs59345481 | 4:90860123 | G / T | 0.10 | 0.09 | 0.08 |
| rs57175349 | 4:90860516 | T / A | 0.08 | 0.07 | 0.07 |
| rs10030931 | 4:90862372 | A / G | 0.43 | 0.37 | 0.38 |
| rs6532205 | 4:90862675 | T / C | 0.53 | 0.46 | 0.46 |
| rs17016396 | 4:90863170 | C / T | 0.08 | 0.07 | 0.06 |
| rs6837483 | 4:90863959 | T / G | 0.08 | 0.07 | 0.06 |
| rs10516853 | 4:90866819 | T / C | 0.09 | 0.09 | 0.08 |
| rs60909103 | 4:90868837 | A / T | 0.09 | 0.09 | 0.08 |
| rs16996201 | 4:90868849 | C / T | 0.07 | 0.07 | 0.06 |
| rs3775482 | 4:90870340 | T / G | 0.37 | 0.43 | 0.41 |
| rs3775483 | 4:90870359 | G / A | 0.37 | 0.43 | 0.42 |
| rs6854896 | 4:90871561 | T / C | 0.37 | 0.33 | 0.33 |
| rs1479429 | 4:90871662 | T / C | 0.37 | 0.33 | 0.33 |
| rs62312664 | 4:90872163 | G / T | 0.25 | 0.24 | 0.24 |
| rs3822098 | 4:90873421 | C / T | 0.42 | 0.38 | 0.39 |
| rs1046994 | 4:90874879 | T / C | 0.37 | 0.42 | 0.42 |
| rs13109927 | 4:90875910 | A / G | 0.35 | 0.41 | 0.39 |
| rs1479427 | 4:90876239 | T / G | 0.37 | 0.33 | 0.33 |
| rs920805 | 4:90876849 | A / G | 0.43 | 0.38 | 0.39 |
| rs13124183 | 4:90877707 | G / A | 0.37 | 0.33 | 0.33 |
| rs6812004 | 4:90877962 | A / G | 0.35 | 0.41 | 0.40 |
| rs6837771 | 4:90877994 | T / A | 0.37 | 0.42 | 0.42 |
| rs1443799 | 4:90878522 | T / C | 0.43 | 0.38 | 0.39 |
| rs17016405 | 4:90878642 | C / T | 0.25 | 0.24 | 0.24 |
| rs7662354 | 4:90879192 | G / A | 0.37 | 0.33 | 0.33 |
| rs7667606 | 4:90879406 | G / A | 0.45 | 0.40 | 0.41 |
| rs1318558 | 4:90879586 | T / C | 0.43 | 0.38 | 0.39 |
| rs1318557 | 4:90879613 | T / C | 0.45 | 0.40 | 0.41 |
| rs12233759 | 4:90880403 | A / G | 0.20 | 0.19 | 0.19 |
| rs56073501 | 4:90881300 | T / C | 0.37 | 0.43 | 0.42 |
| rs62312682 | 4:90881301 | A / G | 0.42 | 0.38 | 0.39 |
| rs3899491 | 4:90881534 | A / G | 0.37 | 0.43 | 0.42 |
| rs1117840 | 4:90881875 | T / C | 0.43 | 0.38 | 0.39 |
| rs61259703 | 4:90881876 | C / G | 0.19 | 0.18 | 0.18 |
| rs1117841 | 4:90882029 | T / A | 0.43 | 0.38 | 0.39 |
| rs11939062 | 4:90882154 | A / C | 0.43 | 0.38 | 0.39 |
| rs11939159 | 4:90882245 | A / G | 0.43 | 0.38 | 0.39 |
| rs11931134 | 4:90882997 | A / G | 0.35 | 0.41 | 0.40 |
| rs11931138 | 4:90883047 | A / C | 0.35 | 0.41 | 0.40 |
| rs11941275 | 4:90883322 | T / C | 0.43 | 0.38 | 0.39 |
| rs1479430 | 4:90883969 | A / G | 0.43 | 0.38 | 0.39 |
| rs6830621 | 4:90884090 | C / T | 0.20 | 0.18 | 0.19 |
| rs11933698 | 4:90884738 | G / A | 0.43 | 0.38 | 0.39 |
| rs13143093 | 4:90885427 | G / A | 0.43 | 0.38 | 0.39 |
| rs58615073 | 4:90886187 | A / G | 0.19 | 0.18 | 0.18 |
| rs112652354 | 4:90886724 | T / G | 0.43 | 0.38 | 0.39 |
| rs142149186 | 4:90887086 | A / G | 0.35 | 0.41 | 0.40 |
| rs35817029 | 4:90887269 | A / T | 0.43 | 0.38 | 0.39 |
| rs34997679 | 4:90887538 | T / C | 0.43 | 0.38 | 0.39 |
| rs112527817 | 4:90890618 | A / G | 0.22 | 0.20 | 0.21 |
| rs142858238 | 4:90890691 | A / G | 0.08 | 0.06 | 0.06 |
| rs6816577 | 4:90890830 | G / A | 0.35 | 0.41 | 0.40 |
| rs62312690 | 4:90891181 | A / G | 0.43 | 0.38 | 0.39 |
| rs12643741 | 4:90891914 | T / A | 0.42 | 0.38 | 0.39 |
| rs13107488 | 4:90895017 | G / A | 0.35 | 0.41 | 0.40 |
| rs13126339 | 4:90895532 | T / C | 0.43 | 0.39 | 0.39 |
| rs2120371 | 4:90896649 | C / A | 0.35 | 0.41 | 0.40 |
| rs7688832 | 4:90896809 | C / T | 0.22 | 0.20 | 0.21 |
| rs13134650 | 4:90896983 | A / C | 0.43 | 0.38 | 0.39 |
| rs13135048 | 4:90896984 | A / G | 0.43 | 0.38 | 0.39 |
| rs13118035 | 4:90898111 | C / T | 0.43 | 0.38 | 0.39 |
| rs992611 | 4:90899053 | C / A | 0.35 | 0.41 | 0.40 |
| rs992612 | 4:90899387 | G / C | 0.43 | 0.39 | 0.39 |
| rs6848497 | 4:90900933 | G / A | 0.22 | 0.20 | 0.21 |
| rs11097240 | 4:90902499 | A / G | 0.35 | 0.41 | 0.40 |
| rs4429701 | 4:90905651 | A / G | 0.42 | 0.38 | 0.39 |
| rs4431189 | 4:90905658 | C / T | 0.46 | 0.41 | 0.42 |
| rs12639645 | 4:90906161 | C / T | 0.42 | 0.38 | 0.39 |
| rs6532209 | 4:90907501 | G / C | 0.35 | 0.42 | 0.41 |
| rs74786971 | 4:90909176 | C / G | 0.07 | 0.08 | 0.06 |
| rs6532210 | 4:90910930 | G / A | 0.38 | 0.44 | 0.43 |
| rs7689553 | 4:90910974 | G / A | 0.35 | 0.42 | 0.41 |
| rs755053 | 4:90912225 | C / T | 0.38 | 0.43 | 0.42 |
| rs13152356 | 4:90913081 | T / C | 0.35 | 0.42 | 0.40 |
| rs35763285 | 4:90913226 | T / C | 0.35 | 0.41 | 0.40 |
| rs13131903 | 4:90913552 | A / G | 0.43 | 0.39 | 0.40 |
| rs6845108 | 4:90914004 | C / T | 0.19 | 0.18 | 0.18 |
| rs2120375 | 4:90914122 | C / A | 0.43 | 0.39 | 0.40 |
| rs993331 | 4:90914149 | A / G | 0.35 | 0.41 | 0.40 |
| rs962542 | 4:90914973 | G / A | 0.35 | 0.41 | 0.40 |
| rs7686547 | 4:90915669 | A / G | 0.35 | 0.41 | 0.40 |
| rs10516854 | 4:90916437 | C / T | 0.34 | 0.40 | 0.39 |
| rs72659437 | 4:90916631 | A / T | 0.05 | 0.06 | 0.05 |
| rs17201659 | 4:90916807 | A / G | 0.34 | 0.40 | 0.39 |
| rs964776 | 4:90917626 | T / G | 0.34 | 0.40 | 0.39 |
| rs992731 | 4:90917853 | A / T | 0.35 | 0.40 | 0.40 |
| rs11940898 | 4:90920067 | T / C | 0.34 | 0.40 | 0.39 |
| rs11097241 | 4:90920339 | C / T | 0.32 | 0.38 | 0.36 |
| rs12649663 | 4:90923250 | C / T | 0.22 | 0.20 | 0.21 |
| rs34886505 | 4:90924425 | T / C | 0.42 | 0.39 | 0.39 |
| rs10516855 | 4:90924478 | C / T | 0.11 | 0.08 | 0.09 |
| rs13113259 | 4:90928959 | G / A | 0.33 | 0.39 | 0.37 |
| rs6855896 | 4:90929621 | A / G | 0.42 | 0.38 | 0.39 |
| rs6855649 | 4:90929794 | A / G | 0.33 | 0.38 | 0.37 |
| rs6856201 | 4:90929905 | C / T | 0.33 | 0.39 | 0.37 |
| rs7682709 | 4:90930982 | T / A | 0.33 | 0.38 | 0.37 |
| rs6822365 | 4:90932208 | G / A | 0.33 | 0.39 | 0.37 |
| rs13148131 | 4:90933505 | A / C | 0.33 | 0.38 | 0.37 |
| rs1838224 | 4:90933830 | C / T | 0.33 | 0.38 | 0.37 |
| rs12639954 | 4:90934573 | A / G | 0.22 | 0.21 | 0.21 |
| rs56131942 | 4:90934896 | T / C | 0.42 | 0.38 | 0.39 |
| rs13132936 | 4:90935105 | T / C | 0.36 | 0.41 | 0.40 |
| rs13125618 | 4:90936165 | C / A | 0.42 | 0.38 | 0.39 |
| rs13139518 | 4:90937853 | T / A | 0.42 | 0.38 | 0.39 |
| rs4568207 | 4:90938255 | G / C | 0.33 | 0.38 | 0.37 |
| rs11724585 | 4:90938460 | G / T | 0.16 | 0.15 | 0.14 |
| rs11728457 | 4:90938575 | G / A | 0.25 | 0.24 | 0.24 |
| rs2120374 | 4:90938621 | A / G | 0.36 | 0.41 | 0.40 |
| rs11736934 | 4:90938815 | T / G | 0.22 | 0.21 | 0.21 |
| rs1443801 | 4:90939397 | C / A | 0.45 | 0.41 | 0.43 |
| rs34868237 | 4:90939451 | A / G | 0.23 | 0.21 | 0.22 |
| rs13102162 | 4:90939567 | G / A | 0.32 | 0.37 | 0.36 |
| rs7688860 | 4:90940988 | A / G | 0.24 | 0.22 | 0.22 |
| rs7667181 | 4:90940990 | T / C | 0.33 | 0.38 | 0.37 |
| rs1838223 | 4:90941048 | G / A | 0.33 | 0.38 | 0.37 |
| rs1838222 | 4:90941051 | T / C | 0.33 | 0.38 | 0.37 |
| rs17016426 | 4:90941351 | G / A | 0.23 | 0.22 | 0.22 |
| rs17202137 | 4:90941366 | C / T | 0.42 | 0.38 | 0.39 |
| rs1443800 | 4:90941508 | G / A | 0.44 | 0.41 | 0.42 |
| rs1822527 | 4:90941644 | A / G | 0.32 | 0.37 | 0.36 |
| rs13144206 | 4:90941790 | A / T | 0.44 | 0.41 | 0.42 |
| rs12641504 | 4:90941822 | G / A | 0.33 | 0.38 | 0.37 |
| rs12644715 | 4:90941931 | C / T | 0.23 | 0.22 | 0.22 |
| rs12642299 | 4:90942633 | C / G | 0.32 | 0.37 | 0.36 |
| rs12645478 | 4:90942641 | C / T | 0.44 | 0.41 | 0.42 |
| rs6812321 | 4:90942824 | G / A | 0.33 | 0.38 | 0.37 |
| rs12642395 | 4:90942826 | A / G | 0.42 | 0.38 | 0.39 |
| rs12642404 | 4:90943038 | T / C | 0.42 | 0.38 | 0.39 |
| rs17016436 | 4:90943444 | A / G | 0.23 | 0.22 | 0.22 |
| rs7662081 | 4:90943750 | G / T | 0.33 | 0.38 | 0.37 |
| rs7661577 | 4:90943785 | G / A | 0.44 | 0.41 | 0.42 |
| rs10516857 | 4:90944370 | C / T | 0.44 | 0.41 | 0.42 |
| rs34343229 | 4:90944806 | A / G | 0.42 | 0.38 | 0.39 |
| rs4561894 | 4:90945456 | T / A | 0.32 | 0.37 | 0.36 |
| rs1443798 | 4:90945779 | G / A | 0.44 | 0.41 | 0.42 |
| rs7670600 | 4:90946314 | T / A | 0.32 | 0.37 | 0.36 |
| rs7671006 | 4:90946548 | T / C | 0.33 | 0.38 | 0.37 |
| rs7693326 | 4:90946665 | A / G | 0.43 | 0.40 | 0.41 |
| rs59746776 | 4:90946729 | T / A | 0.23 | 0.22 | 0.22 |
| rs13121853 | 4:90947396 | A / G | 0.42 | 0.38 | 0.39 |
| rs13148977 | 4:90947538 | C / T | 0.44 | 0.41 | 0.42 |
| rs17808648 | 4:90947793 | G / T | 0.42 | 0.38 | 0.39 |
| rs12648916 | 4:90948523 | A / C | 0.33 | 0.38 | 0.37 |
| rs7660182 | 4:90948929 | A / C | 0.42 | 0.38 | 0.39 |
| rs7685355 | 4:90949156 | C / T | 0.44 | 0.41 | 0.42 |
| rs4626162 | 4:90949562 | T / C | 0.32 | 0.37 | 0.36 |
| rs17016446 | 4:90950230 | A / G | 0.23 | 0.22 | 0.22 |
| rs6830014 | 4:90950726 | G / A | 0.42 | 0.38 | 0.39 |
| rs6830566 | 4:90950762 | G / T | 0.32 | 0.37 | 0.36 |
| rs1982293 | 4:90951276 | A / G | 0.42 | 0.38 | 0.39 |
| rs17808829 | 4:90951958 | A / G | 0.44 | 0.40 | 0.41 |
| rs10516858 | 4:90952424 | A / G | 0.42 | 0.38 | 0.39 |
| rs10516859 | 4:90952582 | T / C | 0.22 | 0.21 | 0.21 |
| rs13107514 | 4:90953022 | C / A | 0.44 | 0.41 | 0.42 |
| rs34064092 | 4:90953974 | T / C | 0.42 | 0.38 | 0.39 |
| rs723992 | 4:90954550 | G / A | 0.44 | 0.41 | 0.42 |
| rs13122257 | 4:90954829 | G / T | 0.32 | 0.37 | 0.36 |
| rs6532211 | 4:90955516 | A / G | 0.33 | 0.38 | 0.37 |
| rs62312738 | 4:90955581 | T / C | 0.42 | 0.38 | 0.39 |
| rs56300941 | 4:90956803 | G / T | 0.32 | 0.37 | 0.36 |
| rs55663876 | 4:90956847 | T / G | 0.42 | 0.38 | 0.39 |
| rs72659455 | 4:90956877 | C / A | 0.23 | 0.22 | 0.22 |
| rs13149395 | 4:90956988 | G / A | 0.42 | 0.38 | 0.39 |
| rs13150124 | 4:90957016 | C / G | 0.42 | 0.38 | 0.39 |
| rs6842029 | 4:90957655 | G / A | 0.33 | 0.38 | 0.37 |
| rs17016459 | 4:90958046 | T / C | 0.23 | 0.22 | 0.22 |
| rs13113035 | 4:90958920 | G / C | 0.30 | 0.35 | 0.34 |
| rs66513408 | 4:90960077 | T / G | 0.23 | 0.22 | 0.22 |
| rs12651123 | 4:90960235 | C / T | 0.42 | 0.38 | 0.39 |
| rs12639636 | 4:90960297 | C / T | 0.42 | 0.38 | 0.39 |
| rs11733658 | 4:90960574 | C / G | 0.23 | 0.22 | 0.22 |
| rs6815240 | 4:90961117 | G / A | 0.33 | 0.38 | 0.37 |
| rs6814978 | 4:90961283 | G / A | 0.44 | 0.41 | 0.42 |
| rs7664476 | 4:90962031 | T / A | 0.44 | 0.40 | 0.42 |
| rs13141175 | 4:90962743 | G / A | 0.44 | 0.40 | 0.42 |
| rs13117400 | 4:90962828 | T / C | 0.33 | 0.38 | 0.37 |
| rs724454 | 4:90963446 | C / A | 0.23 | 0.22 | 0.22 |
| rs724455 | 4:90963720 | T / A | 0.32 | 0.37 | 0.36 |
| rs2120373 | 4:90963742 | C / T | 0.30 | 0.35 | 0.34 |
| rs17809148 | 4:90964088 | C / T | 0.07 | 0.09 | 0.08 |
| rs55686873 | 4:90964584 | T / C | 0.42 | 0.38 | 0.39 |
| rs13104173 | 4:90964718 | C / A | 0.32 | 0.37 | 0.36 |
| rs10516860 | 4:90965087 | G / A | 0.44 | 0.41 | 0.42 |
| rs62312743 | 4:90965967 | C / T | 0.42 | 0.38 | 0.39 |
| rs35197902 | 4:90966183 | A / T | 0.42 | 0.38 | 0.39 |
| rs59913189 | 4:90966849 | T / A | 0.23 | 0.22 | 0.22 |
| rs6532212 | 4:90967670 | A / C | 0.44 | 0.40 | 0.42 |
| rs1443797 | 4:90968156 | G / A | 0.31 | 0.37 | 0.36 |
| rs7675130 | 4:90968757 | A / G | 0.33 | 0.38 | 0.37 |
| rs11929845 | 4:90969565 | A / G | 0.23 | 0.22 | 0.22 |
| rs11929774 | 4:90969583 | C / T | 0.33 | 0.38 | 0.37 |
| rs12639819 | 4:90970261 | T / C | 0.23 | 0.22 | 0.22 |
| rs1443796 | 4:90971423 | G / T | 0.44 | 0.40 | 0.42 |
| rs722937 | 4:90971745 | G / T | 0.31 | 0.37 | 0.36 |
| rs722936 | 4:90971800 | A / C | 0.15 | 0.15 | 0.13 |
| rs12500465 | 4:90971832 | A / G | 0.44 | 0.40 | 0.42 |
| rs13124133 | 4:90972536 | A / G | 0.44 | 0.41 | 0.42 |
| rs13151441 | 4:90972844 | C / T | 0.44 | 0.41 | 0.42 |
| rs13106409 | 4:90973406 | C / T | 0.46 | 0.42 | 0.43 |
| rs6532213 | 4:90974046 | T / A | 0.46 | 0.42 | 0.43 |
| rs6819755 | 4:90974286 | A / G | 0.44 | 0.41 | 0.42 |
| rs1348444 | 4:90975188 | T / C | 0.44 | 0.41 | 0.42 |
| rs72874327 | 4:90975849 | A / G | 0.21 | 0.20 | 0.19 |
| rs1443795 | 4:90976115 | A / T | 0.31 | 0.37 | 0.36 |
| rs13114320 | 4:90976413 | A / G | 0.44 | 0.41 | 0.42 |
| rs6858220 | 4:90977830 | A / C | 0.44 | 0.41 | 0.42 |
| rs11097243 | 4:90979609 | G / T | 0.07 | 0.05 | 0.06 |
| rs1443794 | 4:90979682 | G / A | 0.46 | 0.41 | 0.43 |
| rs78327649 | 4:90980332 | C / G | 0.08 | 0.08 | 0.07 |
| rs13143583 | 4:90980778 | G / A | 0.44 | 0.40 | 0.42 |
| rs13143794 | 4:90980880 | C / A | 0.45 | 0.40 | 0.42 |
| rs6837222 | 4:90981731 | A / G | 0.44 | 0.40 | 0.42 |
| rs17016481 | 4:90982868 | A / G | 0.21 | 0.20 | 0.19 |
| rs13107286 | 4:90983045 | T / A | 0.44 | 0.40 | 0.42 |
| rs13113863 | 4:90983475 | A / G | 0.42 | 0.38 | 0.39 |
| rs1562034 | 4:90983617 | T / C | 0.45 | 0.41 | 0.43 |
| rs12650363 | 4:90983713 | C / T | 0.44 | 0.40 | 0.42 |
| rs2120372 | 4:90983723 | G / A | 0.44 | 0.41 | 0.42 |
| rs12647171 | 4:90983772 | T / G | 0.44 | 0.40 | 0.42 |
| rs12647266 | 4:90984231 | T / C | 0.44 | 0.40 | 0.42 |
| rs1975488 | 4:90984432 | C / T | 0.34 | 0.38 | 0.38 |
| rs57845937 | 4:90984929 | C / T | 0.21 | 0.20 | 0.20 |
| rs13122493 | 4:90985018 | G / A | 0.34 | 0.38 | 0.38 |
| rs6819992 | 4:90988236 | C / T | 0.45 | 0.41 | 0.43 |
| rs6831759 | 4:90989319 | T / C | 0.34 | 0.38 | 0.38 |
| rs72874355 | 4:90989833 | A / G | 0.16 | 0.15 | 0.14 |
| rs13119778 | 4:90989980 | T / C | 0.44 | 0.40 | 0.42 |
| rs13127674 | 4:90991026 | G / C | 0.40 | 0.45 | 0.46 |
| rs6819864 | 4:90991373 | C / T | 0.40 | 0.45 | 0.46 |
| rs894803 | 4:90993172 | C / A | 0.35 | 0.40 | 0.41 |
| rs1031125 | 4:90993342 | A / T | 0.40 | 0.45 | 0.46 |
| rs1808952 | 4:90993888 | G / A | 0.40 | 0.45 | 0.46 |
| rs3919723 | 4:90996034 | T / C | 0.40 | 0.44 | 0.45 |
| rs35049792 | 4:90996189 | C / T | 0.40 | 0.45 | 0.46 |
| rs35474064 | 4:90996192 | A / G | 0.40 | 0.44 | 0.45 |
| rs13108338 | 4:90997150 | C / T | 0.43 | 0.38 | 0.39 |
| rs10516862 | 4:90997171 | T / G | 0.43 | 0.38 | 0.39 |
| rs12649987 | 4:90997355 | C / G | 0.43 | 0.38 | 0.39 |
| rs6826127 | 4:90998124 | C / A | 0.40 | 0.44 | 0.45 |
| rs6838592 | 4:90999570 | G / A | 0.17 | 0.17 | 0.15 |
| rs6816256 | 4:90999635 | T / G | 0.40 | 0.45 | 0.46 |
| rs6839192 | 4:90999769 | C / T | 0.36 | 0.40 | 0.41 |
| rs2197291 | 4:91000334 | C / T | 0.40 | 0.45 | 0.46 |
| rs11931255 | 4:91001043 | G / A | 0.40 | 0.44 | 0.45 |
| rs10516863 | 4:91001818 | T / C | 0.17 | 0.16 | 0.15 |
| rs7655785 | 4:91003238 | C / T | 0.36 | 0.40 | 0.41 |
| rs6818979 | 4:91004112 | C / T | 0.40 | 0.45 | 0.46 |
| rs6844974 | 4:91004271 | T / C | 0.40 | 0.44 | 0.45 |
| rs6824062 | 4:91004427 | G / A | 0.17 | 0.16 | 0.15 |
| rs7668607 | 4:91005062 | T / C | 0.43 | 0.38 | 0.39 |
| rs4383587 | 4:91005096 | A / T | 0.40 | 0.44 | 0.45 |
| rs62313871 | 4:91007345 | T / A | 0.43 | 0.38 | 0.39 |
| rs34243515 | 4:91008837 | C / A | 0.47 | 0.43 | 0.44 |
| rs17204072 | 4:91008877 | T / C | 0.43 | 0.38 | 0.39 |
| rs72874369 | 4:91009715 | T / G | 0.21 | 0.20 | 0.19 |
| rs72659470 | 4:91009758 | A / T | 0.47 | 0.43 | 0.45 |
| rs6844776 | 4:91010920 | C / A | 0.16 | 0.15 | 0.14 |
| rs7671248 | 4:91011689 | T / A | 0.35 | 0.40 | 0.40 |
| rs73832568 | 4:91012162 | C / T | 0.10 | 0.10 | 0.08 |
| rs12648872 | 4:91013383 | G / A | 0.43 | 0.39 | 0.40 |
| rs6849100 | 4:91014732 | C / T | 0.16 | 0.15 | 0.14 |
| rs7684512 | 4:91015463 | G / T | 0.37 | 0.42 | 0.42 |
| rs13132730 | 4:91015597 | G / T | 0.43 | 0.38 | 0.39 |
| rs13104940 | 4:91015600 | G / A | 0.47 | 0.43 | 0.44 |
| rs17204198 | 4:91015888 | T / C | 0.47 | 0.43 | 0.44 |
| rs17016538 | 4:91016797 | T / C | 0.16 | 0.15 | 0.14 |
| rs6855891 | 4:91017611 | T / C | 0.16 | 0.15 | 0.14 |
| rs13120745 | 4:91018145 | G / A | 0.43 | 0.38 | 0.39 |
| rs36047807 | 4:91018641 | T / C | 0.46 | 0.42 | 0.44 |
| rs78785032 | 4:91021009 | G / A | 0.07 | 0.05 | 0.06 |
| rs7672024 | 4:91021291 | C / A | 0.46 | 0.43 | 0.44 |
| rs7677966 | 4:91021712 | A / G | 0.43 | 0.38 | 0.39 |
| rs7677884 | 4:91021879 | A / C | 0.43 | 0.38 | 0.39 |
| rs7438673 | 4:91022130 | G / A | 0.43 | 0.38 | 0.39 |
| rs1838227 | 4:91023312 | T / C | 0.43 | 0.38 | 0.39 |
| rs1838226 | 4:91023371 | G / A | 0.37 | 0.41 | 0.41 |
| rs1838225 | 4:91023473 | G / A | 0.43 | 0.38 | 0.39 |
| rs1031124 | 4:91024383 | T / G | 0.16 | 0.15 | 0.14 |
| rs1031123 | 4:91024525 | G / A | 0.37 | 0.41 | 0.41 |
| rs58507725 | 4:91025637 | T / C | 0.16 | 0.15 | 0.14 |
| rs34001515 | 4:91025850 | G / T | 0.43 | 0.38 | 0.39 |
| rs10516864 | 4:91027048 | G / A | 0.47 | 0.43 | 0.44 |
| rs6824746 | 4:91029192 | C / T | 0.21 | 0.20 | 0.19 |
| rs11932400 | 4:91029513 | G / A | 0.21 | 0.20 | 0.19 |
| rs62313888 | 4:91030144 | T / C | 0.43 | 0.38 | 0.39 |
| rs76735357 | 4:91032538 | A / G | 0.12 | 0.11 | 0.10 |
| rs7674520 | 4:91033300 | C / T | 0.37 | 0.41 | 0.41 |
| rs116761960 | 4:91033521 | G / A | 0.07 | 0.05 | 0.06 |
| rs58617553 | 4:91034241 | G / A | 0.16 | 0.15 | 0.14 |
| rs57352504 | 4:91034415 | T / G | 0.16 | 0.15 | 0.14 |
| rs76793825 | 4:91039202 | A / G | 0.16 | 0.14 | 0.14 |
| rs7679917 | 4:91039783 | C / T | 0.41 | 0.46 | 0.47 |
| rs2870040 | 4:91039894 | T / C | 0.21 | 0.20 | 0.19 |
| rs12644131 | 4:91046719 | C / T | 0.21 | 0.20 | 0.19 |
| rs115406248 | 4:91051522 | C / A | 0.04 | 0.05 | 0.05 |
| rs60520837 | 4:91051688 | C / T | 0.16 | 0.15 | 0.14 |
| rs13132515 | 4:91052890 | C / T | 0.43 | 0.38 | 0.39 |
| rs1963874 | 4:91053985 | C / G | 0.16 | 0.15 | 0.14 |
| rs17016540 | 4:91055884 | C / T | 0.16 | 0.15 | 0.14 |
| rs13146071 | 4:91056544 | T / A | 0.43 | 0.38 | 0.40 |
| rs13116571 | 4:91059512 | G / A | 0.20 | 0.20 | 0.19 |
| rs78032083 | 4:91061671 | G / A | 0.07 | 0.05 | 0.06 |
| rs35545901 | 4:91062485 | A / G | 0.16 | 0.15 | 0.14 |
| rs12650934 | 4:91063703 | T / C | 0.15 | 0.14 | 0.13 |
| rs12650984 | 4:91063747 | A / G | 0.20 | 0.20 | 0.19 |
| rs11947970 | 4:91064462 | T / C | 0.16 | 0.16 | 0.14 |
| rs7677289 | 4:91065352 | C / T | 0.20 | 0.20 | 0.19 |
| rs62313890 | 4:91065700 | G / T | 0.43 | 0.38 | 0.39 |
| rs17016560 | 4:91066125 | T / C | 0.16 | 0.16 | 0.14 |
| rs72874395 | 4:91067136 | A / G | 0.16 | 0.16 | 0.14 |
| rs1946567 | 4:91067333 | C / T | 0.16 | 0.16 | 0.14 |
| rs7667087 | 4:91068074 | G / A | 0.20 | 0.20 | 0.19 |
| rs61308573 | 4:91068594 | A / G | 0.20 | 0.20 | 0.19 |
| rs57291587 | 4:91068613 | T / A | 0.16 | 0.16 | 0.14 |
| rs60347355 | 4:91068669 | T / A | 0.16 | 0.16 | 0.14 |
| rs7697471 | 4:91068675 | T / C | 0.35 | 0.40 | 0.40 |
| rs17016562 | 4:91068924 | C / T | 0.16 | 0.16 | 0.14 |
| rs72874402 | 4:91069070 | G / C | 0.16 | 0.16 | 0.14 |
| rs17016565 | 4:91069629 | G / A | 0.16 | 0.16 | 0.14 |
| rs6837278 | 4:91069888 | G / A | 0.16 | 0.16 | 0.14 |
| rs6838176 | 4:91070086 | A / G | 0.20 | 0.20 | 0.19 |
| rs56054231 | 4:91070977 | A / G | 0.16 | 0.16 | 0.14 |
| rs17016575 | 4:91071042 | C / T | 0.20 | 0.20 | 0.19 |
| rs11935738 | 4:91071069 | C / T | 0.16 | 0.16 | 0.14 |
| rs17016581 | 4:91071661 | T / G | 0.20 | 0.20 | 0.19 |
| rs6834950 | 4:91072701 | C / T | 0.16 | 0.16 | 0.14 |
| rs72876108 | 4:91073774 | C / G | 0.16 | 0.16 | 0.14 |
| rs1812076 | 4:91073943 | A / G | 0.43 | 0.38 | 0.39 |
| rs78969706 | 4:91076862 | A / G | 0.16 | 0.15 | 0.14 |
| rs74413418 | 4:91076866 | A / C | 0.16 | 0.15 | 0.14 |
| rs115709735 | 4:91077522 | A / T | 0.10 | 0.10 | 0.08 |
| rs35212317 | 4:91078066 | A / T | 0.43 | 0.38 | 0.39 |
| rs72876110 | 4:91079739 | A / T | 0.16 | 0.16 | 0.14 |
| rs1835524 | 4:91080308 | C / T | 0.16 | 0.16 | 0.14 |
| rs1835523 | 4:91080598 | G / A | 0.16 | 0.16 | 0.14 |
| rs11732065 | 4:91080624 | C / T | 0.43 | 0.38 | 0.39 |
| rs1835522 | 4:91080676 | G / T | 0.20 | 0.20 | 0.19 |
| rs55924200 | 4:91081279 | A / G | 0.20 | 0.20 | 0.19 |
| rs11937976 | 4:91081489 | T / C | 0.16 | 0.16 | 0.14 |
| rs12645259 | 4:91081909 | A / G | 0.16 | 0.16 | 0.14 |
| rs11944177 | 4:91082781 | C / T | 0.16 | 0.16 | 0.14 |
| rs1835521 | 4:91083051 | G / A | 0.43 | 0.38 | 0.39 |
| rs5006831 | 4:91083582 | G / C | 0.20 | 0.20 | 0.19 |
| rs1835519 | 4:91083630 | C / G | 0.16 | 0.16 | 0.14 |
| rs1835518 | 4:91083645 | T / C | 0.17 | 0.16 | 0.14 |
| rs34146382 | 4:91085477 | A / T | 0.43 | 0.38 | 0.39 |
| rs12645059 | 4:91087140 | A / G | 0.15 | 0.14 | 0.12 |
| rs11945007 | 4:91088473 | C / T | 0.20 | 0.20 | 0.19 |
| rs4403004 | 4:91088562 | A / C | 0.20 | 0.20 | 0.19 |
| rs4339173 | 4:91088742 | T / A | 0.20 | 0.20 | 0.19 |
| rs17016594 | 4:91090849 | A / T | 0.20 | 0.20 | 0.19 |
| rs7655515 | 4:91091035 | G / A | 0.43 | 0.38 | 0.39 |
| rs10026036 | 4:91091770 | C / A | 0.37 | 0.42 | 0.42 |
| rs17016596 | 4:91092938 | G / A | 0.20 | 0.20 | 0.19 |
| rs1986976 | 4:91093620 | A / G | 0.20 | 0.20 | 0.19 |
| rs11947949 | 4:91094329 | A / G | 0.16 | 0.16 | 0.14 |
| rs17205805 | 4:91094931 | A / G | 0.42 | 0.38 | 0.39 |
| rs6532214 | 4:91095875 | G / A | 0.16 | 0.16 | 0.14 |
| rs11934796 | 4:91096995 | G / A | 0.16 | 0.16 | 0.14 |
| rs6532215 | 4:91097819 | G / A | 0.20 | 0.20 | 0.19 |
| rs61394011 | 4:91099139 | T / C | 0.16 | 0.16 | 0.14 |
| rs6813752 | 4:91099481 | T / A | 0.37 | 0.41 | 0.41 |
| rs7680617 | 4:91100130 | T / G | 0.16 | 0.16 | 0.14 |
| rs2726 | 4:91102650 | G / A | 0.16 | 0.16 | 0.14 |
| rs60047025 | 4:91102704 | G / A | 0.16 | 0.16 | 0.14 |
| rs10014272 | 4:91103640 | T / C | 0.37 | 0.42 | 0.42 |
| rs6829376 | 4:91109167 | T / C | 0.12 | 0.11 | 0.11 |
| rs12331460 | 4:91109576 | G / A | 0.15 | 0.14 | 0.14 |
| rs9307081 | 4:91110326 | A / G | 0.15 | 0.14 | 0.14 |
| rs78876299 | 4:91111063 | C / A | 0.06 | 0.06 | 0.05 |
| rs17016622 | 4:91111631 | G / A | 0.12 | 0.11 | 0.11 |
| rs7698120 | 4:91112815 | A / G | 0.12 | 0.11 | 0.11 |
| rs72876151 | 4:91113471 | C / G | 0.12 | 0.11 | 0.11 |
| rs57711133 | 4:91113668 | C / G | 0.15 | 0.14 | 0.14 |
| rs13103611 | 4:91114142 | G / T | 0.44 | 0.39 | 0.41 |
| rs7664833 | 4:91114979 | T / C | 0.39 | 0.37 | 0.38 |
| rs6840859 | 4:91116003 | C / G | 0.47 | 0.49 | 0.49 |
| rs6818163 | 4:91116065 | T / C | 0.48 | 1.00 | 0.50 |
| rs6840905 | 4:91116126 | A / C | 0.47 | 0.49 | 0.50 |
| rs6841431 | 4:91116156 | A / G | 0.09 | 0.10 | 0.09 |
| rs62313893 | 4:91116462 | A / G | 0.15 | 0.12 | 0.13 |
| rs963207 | 4:91117003 | G / A | 0.47 | 0.49 | 0.49 |
| rs7694967 | 4:91117474 | A / G | 0.48 | 1.00 | 0.50 |
| rs7695158 | 4:91117476 | C / T | 0.48 | 1.00 | 0.50 |
| rs6854773 | 4:91117965 | G / T | 0.48 | 1.00 | 0.50 |
| rs78694949 | 4:91119554 | G / A | 0.07 | 0.05 | 0.06 |
| rs13146102 | 4:91121536 | T / A | 0.51 | 0.49 | 0.49 |
| rs7676382 | 4:91121900 | A / G | 0.51 | 0.49 | 0.49 |
| rs12645744 | 4:91122378 | A / G | 0.51 | 0.49 | 0.49 |
| rs7681376 | 4:91122398 | G / A | 0.51 | 0.49 | 0.49 |
| rs10461167 | 4:91123414 | C / T | 0.51 | 0.49 | 0.49 |
| rs10461168 | 4:91123665 | T / G | 0.51 | 0.49 | 0.49 |
| rs13109951 | 4:91123874 | G / A | 0.24 | 0.21 | 0.22 |
| rs4428251 | 4:91124595 | A / G | 0.48 | 1.00 | 0.50 |
| rs6532216 | 4:91126063 | C / T | 0.48 | 1.00 | 0.50 |

**Table S3. All variants found in full sequencing of *SNCA* with targeting next generation sequencing.**

| **Variant Name** | **Position** | **Allele effect/reference** | **Freq. Cases** | **Freq. Controls** | **Variant Type** |
| --- | --- | --- | --- | --- | --- |
| rs529553259 | 4:90645459 | A / T | 0.003561 | 0.001771 | UTR3 |
|  | 4:90645578 | T / G | 0.0005165 | 0 | UTR3 |
| rs774934945 | 4:90645632 | C / A | 0.0005086 | 0.0005903 | UTR3 |
|  | 4:90645641 | A / G | 0.0005086 | 0 | UTR3 |
| rs1045722 | 4:90645671 | A / T | 0.08494 | 0.06553 | UTR3 |
| rs3857053 | 4:90645674 | T / C | 0.08494 | 0.06494 | UTR3 |
|  | 4:90645943 | G / A | 0.001017 | 0 | UTR3 |
|  | 4:90646085 | A / C | 0.0005133 | 0 | UTR3 |
|  | 4:90646095 | T / C | 0.0005128 | 0 | UTR3 |
|  | 4:90646101 | T / C | 0 | 0.001181 | UTR3 |
|  | 4:90646118 | C / A | 0.0005086 | 0 | UTR3 |
|  | 4:90646230 | A / G | 0.0005258 | 0 | UTR3 |
| rs752617075 | 4:90646289 | G / A | 0.0005247 | 0 | UTR3 |
|  | 4:90646298 | A / G | 0.0005258 | 0 | UTR3 |
|  | 4:90646300 | T / G | 0.0005258 | 0 | UTR3 |
|  | 4:90646301 | A / G | 0.0005258 | 0 | UTR3 |
| rs560896156 | 4:90646302 | G / A | 0.0005258 | 0 | UTR3 |
|  | 4:90646303 | A / G | 0.0005258 | 0 | UTR3 |
|  | 4:90646327 | G / A | 0.0005086 | 0 | UTR3 |
|  | 4:90646384 | T / G | 0.0005086 | 0 | UTR3 |
|  | 4:90646395 | C / T | 0.0005086 | 0 | UTR3 |
|  | 4:90646410 | T / G | 0.001017 | 0 | UTR3 |
| rs915091114 | 4:90646639 | A / G | 0.0005086 | 0.001181 | UTR3 |
|  | 4:90646676 | T / C | 0.0005086 | 0 | UTR3 |
| rs561544802 | 4:90646677 | A / G | 0.001017 | 0 | UTR3 |
|  | 4:90646681 | T / A | 0.001526 | 0 | UTR3 |
| rs886059723 | 4:90646701 | T / TA | 0.0005086 | 0 | UTR3 |
| rs938774312 | 4:90646709 | * / G | 0.001017 | 0 | UTR3 |
|  | 4:90646712 | A / G | 0.0005086 | 0 | UTR3 |
| rs577490090 | 4:90646919 | T / A | 0.003052 | 0.001771 | UTR3 |
|  | 4:90647027 | A / G | 0.001117 | 0 | UTR3 |
|  | 4:90647270 | A / AAC | 0 | 0.0005903 | UTR3 |
|  | 4:90647271 | * / A | 0 | 0.0005903 | UTR3 |
| rs17016074 | 4:90647278 | A / G | 0.0117 | 0.009445 | UTR3 |
| rs193289281 | 4:90647506 | A / G | 0 | 0.0005903 | UTR3 |
|  | 4:90647599 | A / G | 0.0005086 | 0 | UTR3 |
| rs10024743 | 4:90647640 | C / A | 0.0005086 | 0 | UTR3 |
| rs144511886 | 4:90647663 | A / G | 0 | 0.0005903 | UTR3 |
| rs145304567 | 4:90647702 | T / G | 0.00763 | 0.01004 | UTR3 |
|  | 4:90647704 | G / A | 0.0005086 | 0 | UTR3 |
| rs76642636 | 4:90647794 | A / G | 0.003052 | 0.001771 | synonymous SNV |
| rs1358566725 | 4:90650365 | T / C | 0.0005086 | 0 | nonsynonymous p.A124T |
| rs749476922 | 4:90650370 | G / T | 0 | 0.001181 | nonsynonymous p.N122S |
| rs145138372 | 4:90650386 | T / G | 0.0005086 | 0 | nonsynonymous p.P117S |
|  | 4:90650465 | C / T | 0.0005086 | 0 | intronic |
|  | 4:90743413 | C / T | 0.0005086 | 0 | nonsynonymous p.K97R |
| rs762146888 | 4:90743555 | T / A | 0 | 0.0005903 | intronic |
| rs200475984 | 4:90743572 | A / G | 0.0005086 | 0 | intronic |
| rs771528614 | 4:90749256 | C / T | 0 | 0.0005903 | intronic |
| rs141859659 | 4:90749310 | T / C | 0.001017 | 0 | synonymous SNV |
| rs745412490 | 4:90756825 | C / T | 0 | 0.0005903 | UTR5 |
| rs2583986 | 4:90757947 | A / T | 0.1915 | 0.27 | UTR5 |
| rs548276692 | 4:90758106 | T / C | 0.0005531 | 0.0005959 | intronic |
|  | 4:90758116 | CC / TC | 0.0005501 | 0 | intronic |
|  | 4:90758152 | A / G | 0.0005102 | 0 | intronic |
|  | 4:90758172 | T / G | 0.0005102 | 0 | intronic |
|  | 4:90759407 | C / T | 0 | 0.0005903 | ncRNA splicing |
